# A MYT1L Syndrome mouse model recapitulates patient phenotypes and reveals altered brain development due to disrupted neuronal maturation

**DOI:** 10.1101/2020.12.17.423095

**Authors:** Jiayang Chen, Mary E. Lambo, Xia Ge, Joshua T. Dearborn, Yating Liu, Katherine B. McCullough, Raylynn G. Swift, Dora R. Tabachnick, Lucy Tian, Kevin Noguchi, Joel R. Garbow, John N. Constantino, Harrison W. Gabel, Keith B. Hengen, Susan E. Maloney, Joseph D. Dougherty

## Abstract

Human genetics have defined a new autism-associated syndrome caused by loss-of-function mutations in *MYT1L*, a transcription factor known for enabling fibroblast-to-neuron conversions. However, how *MYT1L* mutation causes autism, ADHD, intellectual disability, obesity, and brain anomalies is unknown. Here, we develop a mouse model of this syndrome. Physically, *Myt1l* haploinsufficiency causes obesity, white-matter thinning, and microcephaly in the mice, mimicking clinical phenotypes. During brain development we discovered disrupted gene expression, mediated in part by loss of *Myt1l* gene target activation, and identified precocious neuronal differentiation as the mechanism for microcephaly. In contrast, in adults we discovered that mutation results in failure of transcriptional and chromatin maturation, echoed in disruptions in baseline physiological properties of neurons. This results in behavioral anomalies including hyperactivity, muscle weakness and fatigue, and social alterations with more severe phenotypes in males. Overall, our findings provide insight into the mechanistic underpinnings of this disorder and enable future preclinical studies.

## Introduction

Genetic mutations in a variety of highly constrained human genes have been strongly associated with Intellectual Disability (ID) and other developmental disorders including Autism Spectrum Disorder (ASD), providing a great opportunity to understand these disorders. Many forms of ID/ASD are caused by mutations in chromatin regulators/transcription factors (TFs), such as *FOXP1, MECP2, SETD5* and *CHD8*, and rodent models have been essential to defining the CNS consequences of these mutations (Anderson et al., 2020; Gompers et al., 2017; Guy et al., 2001; Katayama et al., 2016; Sessa et al., 2019). Now, recent human genetic studies discovered *de novo* mutations in another TF, *MYT1L*, to be strongly associated with ID (Blanchet et al., 2017; de Ligt et al., 2012; Loid et al., 2018; Windheuser et al., 2020a) and ASD (De Rubeis et al., 2014; Sanders, 2015; Satterstrom et al., 2018). Most individuals with these *MYT1L* mutations (or larger deletions of the entire 2p25.3 region) suffer from ID with a subset also showing ASD and/or ADHD (Blanchet et al., 2017; Mansfield et al., 2020; Windheuser et al., 2020a). Other consequences described by these groups include microcephaly, white-matter thinning, obesity, epilepsy, and neuroendocrine disruptions, as well as partially penetrant physical abnormalities (clinodactyly and strabismus). However, histological, cellular, and molecular consequences of germline *MYT1L* mutation are not yet defined. A better understanding of the function of MYT1L in the developing brain may clarify the pathobiology of this syndrome.

Prior studies, primarily in cell culture, have proposed some molecular and developmental functions for MYT1L. MYT1L is a CCHC zinc finger transcription factor that is highly expressed in the developing brain (Kim et al., 1997; Matsushita et al., 2014; Weiner and Chun, 1997). MYT1L is most well-known from overexpression studies, where high levels of MYT1L, along with BRN2 and ASCL1, reprogram fibroblasts directly into neurons. This indicates it can, in combination with other factors, play an instructive role in regulating neuronal differentiation (Pang et al., 2011; Vierbuchen et al., 2010). ChIP-seq studies in this system indicated MYT1L binds specific targets, especially promoters. Comparison of these targets with RNA-seq changes in fibroblasts following MYT1L overexpression led to the conclusion that MYT1L was a new kind of repressor specifically of non-neuronal gene expression, thereby restricting cell potential away from non-neuronal fates (Mall et al., 2017). This fits with an earlier report indicating that at least an isolated central domain of MYT1L can bind the repressor SIN3B (Romm et al., 2005) and MYT1L can repress gene expression (Manukyan et al., 2018). The related *MYT1* gene was shown in the same work to recruit histone deacetylases - proteins which have the capacity to close chromatin. However, work on synthetic reporters showed MYT1L tended to activate gene expression 4-5 fold overall (Jiang et al., 1996; Manukyan et al., 2018), and this could be mapped to the N-terminal domain (Manukyan et al., 2018). Yet, while overexpression of MYT1L in u87 glioma lines both increased and decreased gene expression, the reported MYT1L binding motif (AAA[C/G]TTT) was enriched primarily in the promoters of repressed genes, and luciferase activity for 3 of 5 endogenous targets showed repression (Manukyan et al., 2018). Thus, whether MYT1L activates gene expression through a different motif, or through cooperativity with other transcription factors is not clear. Further, the direct impact of MYT1L knockdown or overexpression on chromatin accessibility has not been assessed.

As opposed to overexpression as highlighted in the cell culture studies above, most *MYT1L* variants associated with developmental disabilities are predicted to be heterozygous loss-of-function, suggesting haploinsufficiency as the primary mechanism in disease. Yet, there is no mammalian system to study the consequences of *MYT1L* germline haploinsufficiency or complete loss *in vivo*. Likewise, there have been no comprehensive studies to define the normal role of MYT1L in developing and mature brains. Evidence from neuronal differentiation of neural progenitor cells suggests that, rather than only repress non-neuronal genes, MYT1L was also necessary for activating neuronal genes (Kepa et al., 2017). Specifically, upon shRNA knockdown, far more genes were decreased than increased, consistent with loss of an activator, and these corresponded to synaptic proteins that mark neuronal maturation. Indeed, with MYT1L shRNA, neurons failed to reach sufficient electrophysiological maturation to enable action potentials (Mall et al., 2017). Further, a morpholino study in zebrafish followed by *in situ* hybridization for two neuropeptides (Blanchet et al., 2017), showed a loss of these transcripts, both known to express late in maturation (Almazan et al., 1989). Thus, overall, the human brain development phenotypes could either be related to a loss of a repressor (i.e., ectopic expression of non-neuronal genes) as suggested by overexpression, or a loss of an activator that promotes neuronal differentiation, as suggested by shRNA. Determining whether loss of MYT1L levels results in more opening or closing of chromatin, and the corresponding consequence on gene expression, could inform whether it functions as an activator, repressor, or both *in vivo.* Furthermore, characterizing a mammalian model for the downstream consequences on brain development, neurophysiology, and behavioral circuit function would provide insight into the conserved roles of this protein, as well as provide a tool for future studies of the disease.

Therefore, we developed a mouse model to understand the consequences of MYT1L mutation *in vivo*, inspired by a local patient who had an exon 10 stop gain resulting in ASD, ADHD, ID and other characteristics. We find MYT1L haploinsufficiency alters chromatin accessibility and corresponding gene expression during development leading to precocious neuronal differentiation and smaller brains, though without obvious ectopic non-neuronal gene expression. Postnatally, genomic studies reveal a neuronal maturation failure, along with corresponding electrophysiological abnormalities, finally resulting in clinically-relevant muscle weakness and fatigue, obesity, hyperactivity, and social orientation deficits as revealed by a novel social motivation assay. This new model provides mechanistic insights into MYT1L function *in vivo,* and preclinical opportunities for novel therapeutic development for MYT1L Syndrome.

## Results

### MYT1L is expressed in neuronal lineages across key developmental windows

To establish where MYT1L functions, we first defined its expression across development. First, we looked at its temporal expression in mice, which would guide spatial expression studies afterwards. We found *MYT1L* mRNA increased across neurogenesis and peaked on postnatal day (P)1 yet sustained low levels into adulthood (Fig. 1A), paralleling human expression (Fig. S1A). Further, MYT1L maintained similar protein levels from embryonic day (E)14 to P1 then declined (Fig. 1B). Spatially, initial studies highlighted expression in new neurons of the developing brain (Kim et al., 1997), with an absence in glia. In contrast, a recent report proposed expression in oligodendroglia, promoting their fate (Shi et al., 2018). To resolve this inconsistency, we next investigated MYT1L’s cellular expression.

**Figure 1:**
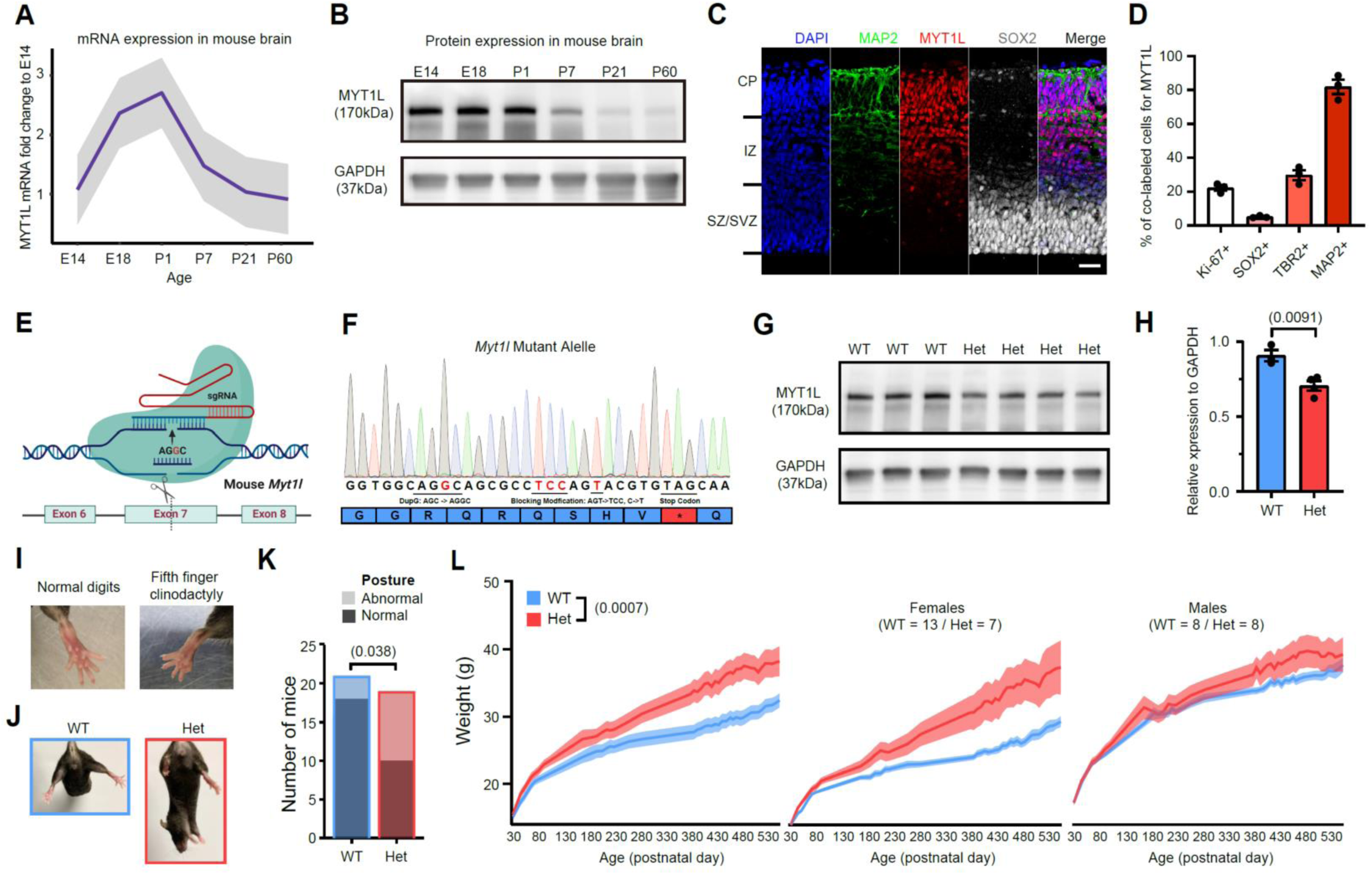
MYT1L frameshift mutation results in protein haploinsufficiency, physical anomalies, and obesity. qRT-PCR revealed trajectory of MYT1L mRNA expression across mouse brain development (n = 3 for each time point). (**B**) Western blot showed a parallel trajectory of protein levels. **(C)** Immunofluorescence of MYT1L protein (red) revealed expression in Map2+ (green) Cortical Plate (CP), intermediate zone (IZ), and a few in SOX2+ (white) progenitors in the Ventricular Zone/SubVentricular Zone (VZ/SVZ). Scale bar, 50 μm. **(D)** Quantification of MYT1L+ fraction within four different cell types (n = 3). **(E)** Schematics for MYT1L KO mouse line generation using CRISPR-Cas9 and donor oligonucleotide. **(F)** Sanger sequencing confirmed c.3035dupG mutation, which leads to frameshift and premature stop codon gain on MYT1L mutant allele. **(G)** Western blot on P1 whole brain lysates confirmed MYT1L protein reduction in Het mice with **(H)** quantification. In physical examination, a subset of Het mice displayed **(I)** fifth finger clinodactyly, **(J,K)** abnormal hindlimb posture representation with **(K)** quantification. **(L)** Also, Het mice weighed significantly more than WTs as adults, which was more pronounced in females. We also observed the expected overall greater weight in males compared to females. *Data were represented as mean ± SEM. See also* Figure S1 *and* ***Table S5*** *for statistical test details*.

Immunofluorescence (IF) during peak cortical neurogenesis (E14), with a knockout-validated MYT1L antibody (Fig. S1J), revealed MYT1L’s gradient of expression in the cortex and medial ganglionic eminence: almost absent in the progenitor layers (SOX2+) and highest in the upper cortical plate (CP; Fig. 1C,D, S1B,C), mirroring prior studies (Kim et al., 1997; Matsushita et al., 2014; Weiner and Chun, 1997). This parallels neuronal maturation gradients, with dim intermediate zone (IZ) expression, where immature neurons are found, and strongest expression in CP. A small portion of proliferating cells (Ki-67+), or intermediate progenitors (TBR2+) expressed MYT1L (Fig. 1D, S1C) at the IZ border. This suggests MYT1L may be expressed as progenitors exit the cell cycle. In neonates, MYT1L was in postmitotic neurons (BRN2+ and CTIP2+) and small portion of radial glia (SOX2+) but not in oligodendroglia (OLIG2+; Fig. S1D,E). In adults, MYT1L was expressed in neurons (NEUN+) across all regions examined (Fig. S1F,G). MYT1L was not found in astrocytes (GFAP+) nor oligodendroglia (OLIG2+; Fig. S1H). Collectively, our expression studies indicate MYT1L’s function commences concurrently with final proliferation of neuronal progenitors, and its expression in all postmitotic neurons across development implies MYT1L haploinsufficiency potentially influences any neuron type. Further, the timeline suggests a peak function during neuronal maturation, but does not rule out a sustained role in adult neurons. Finally, we replicated observations (Kim et al., 1997; Mall et al., 2017) that MYT1L is not expressed in glia, and thus impacts on glia (i.e., white-matter thinning) must be indirect. In addition to establishing where and when MYT1L deficiency might act, we also generated a mouse to determine the consequences of its loss.

### Generation and characterization of *Myt1l* knockout mice

Germline mutants of *Myt1l* would enable studies of its role in CNS development, its function on chromatin, gene expression, and the cellular, physiological, and behavioral phenotypes of haploinsufficiency. Therefore, we used CRISPR-mediated homology-directed repair (HDR) to generate mice with a mutation on exon7 (chr12:29849338, c.3035dupG, S710fsX; Fig. 1E), inspired by a *MYT1L* patient mutation in the homologous exon 10 (**Table S1**), resulting in frameshift and a predicted stop-gain (Fig. 1F). As we found *Myt1l* homozygous mutant (KO) mice die at birth, we confirmed *Myt1l* transcripts and protein decreased in heterozygous mice (Het; Fig. 1G,H, S1K), and IF showed complete MYT1L protein loss in KO E14 mouse cortex (Fig. S1J). No truncated protein (est. 80.63 kDa) was produced by the mutation (Fig. S1I). Further sequencing of the cDNA from Hets revealed a depletion of the mutant mRNA compared to genomic controls, consistent with nonsense mediated decay (Fig. S1L). Thus this mutation appears to result in haploinsufficiency.

Next, we assessed mice for physical abnormalities, including obesity, reported in patients. We observed clinodactyly (Fig. 1I) and early age cataracts. In addition, we observed abnormal hindlimb posture: transient hyperflexions of one or both hindlimbs (Fig. 1J,K), reflected not in coordinated clasping, but in holding limb(s) at midline. Fifth finger clinodactyly and eye issues (strabismus) have been reported in patients. Finally, we also observed obesity in Hets. There was an initial separation of group weights at P45 which became statistically significant at P94, and was more pronounced in females than males (Fig. 1L). Thus, *Myt1l* mutation results in physical alterations and obesity in mice and humans.

### MYT1L haploinsufficiency results in microcephaly and thinned white-matter

Almost half of patients have CNS malformations like microcephaly, hydrocephaly and thinned white-matter. Therefore, we investigated structural abnormalities in P60 Hets with Nissl staining (Fig. 2A,B). Brain organization was grossly normal, yet Hets showed decreased brain weight and smaller cortical volume (Fig. 2C,D) with no change in cortex/brain ratio (Fig. S2A) compared to WTs. For white-matter, there was a trend towards reduced corpus callosum volume (Fig. S2B). In addition, there was no cell density change in Het cortex (Fig. S2C), indicating that microcephaly in Hets corresponds to fewer cells, rather than less parenchyma. We next investigated mouse brains with magnetic resonance (MR)-based Diffusion Tensor Imaging (DTI), a more sensitive, *in vivo,* clinically-translatable technique that can provide both structural and functional information (Fig. 2E). From maps of apparent diffusion coefficient (ADC; Fig. S2D) and fractional anisotropy (FA; Fig. 2F), we segmented several brain regions and performed 3D reconstruction (Fig. 2G, S2E,F). By MRI, Hets again had smaller brain volumes, with no size change in the ventricular system (Fig. 2H, S2G). With segmentation of FA maps (Fig. 2F,G), we found Hets had a smaller corpus callosum volume (Fig. 2I). Functionally, FA values were unchanged in white matter and cortex, suggesting that remaining axons were not abnormal (Fig. S2H). As MYT1L was not expressed in oligodendroglia (Fig S1E,H), this suggests the white matter decrease reflected a decreased number of axons rather than oligodendroglial dysfunction (e.g. dysmyelination). Overall, *Myt1l* mutation results in both decreased brain size and smaller specific white-matter tracts.

**Figure 2:**
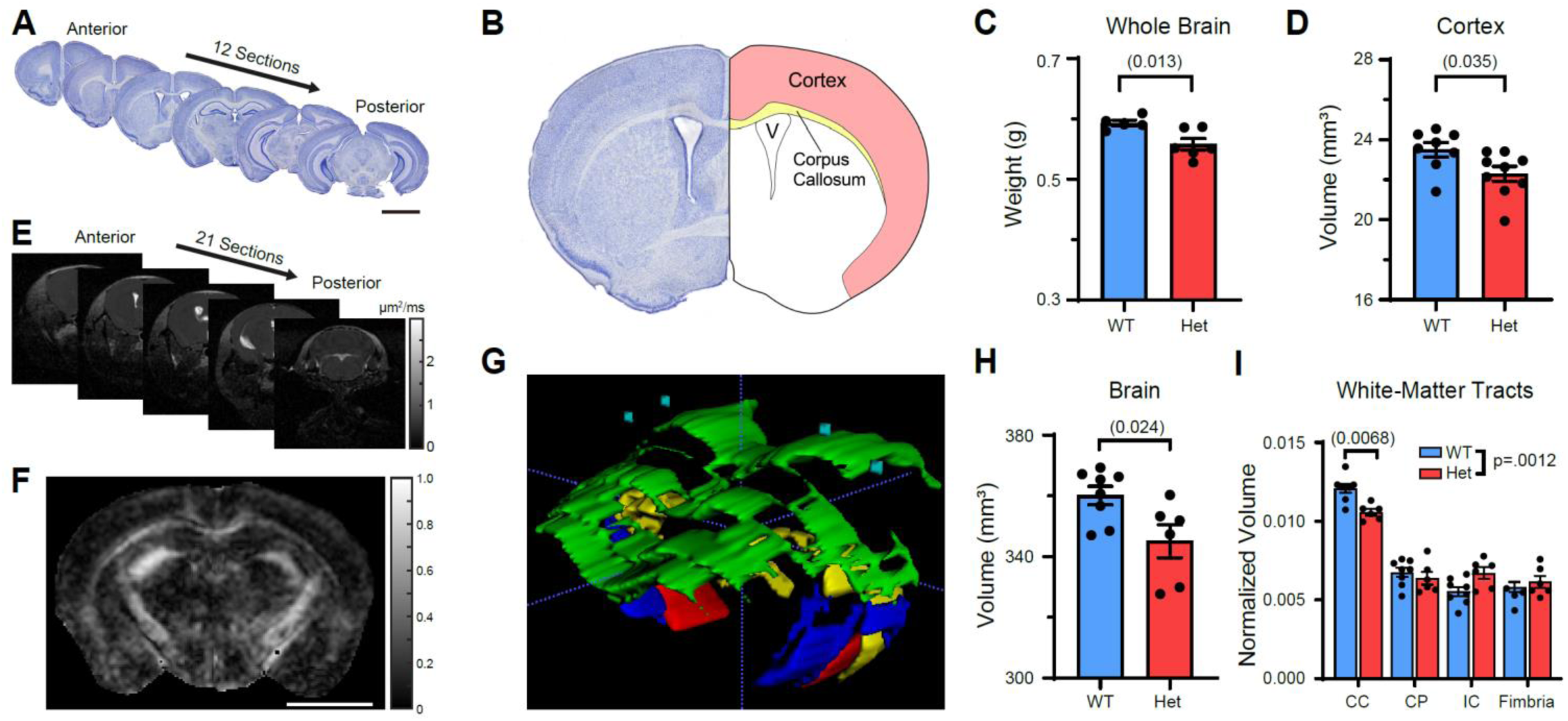
MYT1L haploinsufficiency causes microcephaly and white-matter thinning in corpus callosum. **(A)** Sectioning strategy for Nissl staining. Scale bar, 3 mm. **(B)** Diagram of different brain structures examined. **(C)** Adult Het mice had decreased brain weight and **(D)** decreased cortical volume. **(E)** Coronal images acquired from DTI. **(F)** Fractional anisotropy (FA) map for visualization of white-matter tracts. Scale bar, 0.5 cm. **(G)** 3D reconstruction of different white-matter tracts via FA mapping, including corpus callosum (CC, green), cerebral peduncle (CP, red), internal capsule (IC, blue), fimbria (yellow), and cortex (blue). **(H)** DTI recapitulated smaller brain phenotype in Het mice. **(I)** Histogram showed adult Het mice had decreased corpus callosum volume with other white-matter tracts unchanged. and **(J)** *Data were represented as mean ± SEM. See also* Figure S2 *and* ***Table S5*** *for statistical test details*.

### MYT1L loss alters chromatin state during early mouse brain development

We next conducted genomic studies in the developing brain to 1) determine the function of MYT1L in the embryonic brain, and 2) understand the developmental deficits that might cause the structural phenotypes. We focused on E14, when MYT1L begins expression (Fig 1), and to leverage previous chromatin ChIP-seq analysis from E13.5 (Mall et al., 2017) to examine the consequences of MYT1L loss at direct binding targets. At E14, we could also assay KOs which may further potentiate any molecular phenotypes, and with Hets allow for identification of regions that respond linearly to gene dose.

First, we performed Assay for Transposase Accessible Chromatin(ATAC)-seq (6 WT, 5 Het, and 6 KO E14 cortex, Fig. S3A) to determine how MYT1L loss alters chromatin accessibility. MTY1L is thought to modulate chromatin (Romm et al., 2005), with recent overexpression studies highlighting a repressor role (Mall et al., 2017). We sought here to determine if it has the same role during normal brain development. We identified 1965 (FDR<.05, 4837 FDR<.1, **Table S2**) differentially accessible regions (DARs) in mutants (Het and KO; Fig. 3A,B, S3B,C). Interestingly, KO mice showed smaller changes than Hets in terms of DARs decreasing accessibility after *Myt1l* mutation (Fig. 3A), indicating a possible compensatory mechanism triggered by complete loss of MYT1L resulting in more activated chromatin structures. Motif analysis on DARs revealed that regions losing accessibility in mutants were enriched for motifs of stem-cell TFs (*Lhx2*, *Sox2*), as well as the key neurogenic TF *Ascl1*. More-accessible DARs are enriched for motifs of pro-differentiation TFs (*NF-1* and *Olig2*; Fig. 3C **& Table S2**). A Gene Ontology (GO) analysis on DARs located in transcriptional start sites (TSS) (**Table S4**) revealed less-accessible TSS were enriched for cell cycle and neurogenesis pathways (Fig 3D).

**Figure 3:**
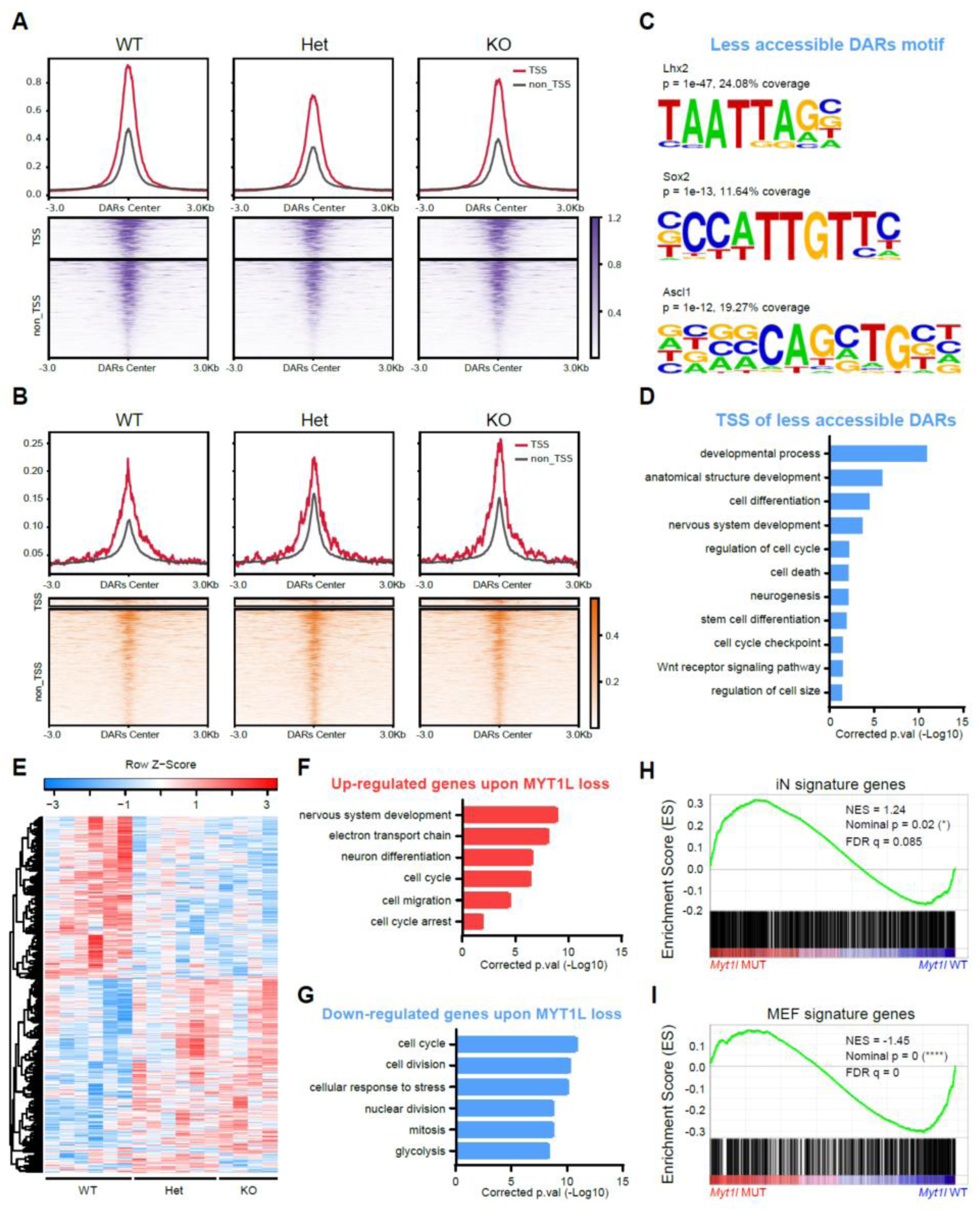
Chromatin Accessibility and RNA-seq analysis define molecular consequences of MYT1L loss in the developing brain. **(A)** Less and **(B)** more-accessible regions in *MYT1L* mutant E14 mouse cortex identified by ATAC-seq (FDR < .1). **(C)** Homer motif analysis on less-accessible DARs over more-accessible DARs. **(D)** GO analysis on less-accessible DARs associated genes showed the disruption of neurodevelopmental programming in mutants. **(E)** Heatmap for differential gene expression in mutant (FDR < .1). **(F)** GO analysis on DEGs revealed an upregulation of early neuronal differentiation pathways and **(G)** a downregulation of cell proliferation programs. **(H)** GSEA analysis found iN signature genes increased expression while **(I)** MEF genes decreased expression in mutants cortex. *See also* Figure S3 *and* ***Table S5*** *for statistical test details*.

Second, we examined MYT1L binding targets defined from E13.5 brain and reprogramming fibroblasts (Mall et al., 2017). On the balance, we found MYT1L loss decreased the accessibility of bound regions (Fig. S3D), suggesting loss of an activator. Also, there were more ChIP targets overlapped with less-accessible DARs than those overlapped with more-accessible DARs (*χ*^2^(1, *N*=203)=11.48, *p*=.0007), further arguing MYT1L’s direct function as an activator to open the chromatin during CNS development. However, only a small subset of ChIP targets were DARs (3.62% of 6652 ChIP peaks). Thus, chromatin accessibility changes in mutants can be attributed to both direct and indirect effects. In sum, MYT1L loss alters chromatin accessibility, including directly bound targets, which likely leads to neurodevelopmental gene expression deficits (Fig. 3D).

### MYT1L loss alters gene expression during early mouse brain development

To understand the transcriptional consequences of this altered chromatin, we conducted RNA-seq on E14 mouse cortex (6 WT, 6 Het, and 4 KO E14 cortex, Fig. S3E). Among 13846 measurable genes, we identified 1768 differentially expressed genes (DEGs; Fig. 3E **& Table S3**). Fold changes of DEGs correlated well between Het and KO datasets, with larger effects in KOs (Fig. S3F,G). This is consistent with a dose-dependence for MYT1L transcriptional regulatory activity at many targets.

Decreased gene expression can be caused by TSS closure, so we plotted the ATAC-seq fold changes for TSS of all DEGs. Indeed, there was a concordance between ATAC-seq TSS and RNA-Seq changes (Fig. S3H). In addition, unlike in neuronal reprogramming where MYT1L overexpression mostly suppressed the expression of its ChIP-seq targets, those targets showed subtle decreased expression in our E14 RNA-seq upon MYT1L loss (Fig. S3I,J). Generally, changes in DEGs identified from MYT1L mouse mutants did not correlate with changes seen in prior RNA-seq of MYT1L overexpression (OE) or knockdown (KD) experiments in cultured MEFs and neurons, respectively (Mall et al., 2017)(Fig. S3M-P). We also categorically defined ‘*in vitro* MYT1L repressed’ genes (downregulated by OE, upregulated by KD) and ‘*in vitro* MYT1L induced’ genes (upregulated by OE, downregulated by KD). In these categorical analyses, we found downregulated genes from our *in vivo* RNA-seq did include 33 *in vitro* MYT1L induced genes from cultures (p<.0005). However, our upregulated genes did not show significant overlap with MYT1L repressed genes (Fig. S3Q,R). Collectively, the loss of expression of MYT1L target genes with MYT1L mutation indicates MYT1L functions preponderantly as an activator during early brain development. This is distinct from the repressor of non-neuronal lineages function reported in direct conversion by OE or KD *in vitro* (Mall et al., 2017). Specifically, we observed no de-repression of the previously described non-neuronal lineage genes with MYT1L mutation (Fig. S3K), suggesting this proposed new class of repressor function (Mall et al., 2017) might be relevant in direct reprogramming, but not reflect the *in vivo* role.

Next, as adult structural abnormalities can be attributed to deficits during brain development, we examined the gestalt of the RNA-seq using GO analysis (**Table S4**). There was an upregulation of CNS development pathways (Fig. 3F) in mutants, driven by markers of neuronal differentiation. This suggests early differentiation in mutants. Likewise, examination of induced neuron (iN) and mouse embryonic fibroblast (MEF) signature genes in our RNA-seq with Gene Set Enrichment Analysis (GSEA) discovered a downregulation of MEF genes and upregulation of iN genes in E14 mutants (Fig. 3H,I), indicating mutant cortex shifted profiles towards early neuronal differentiation. This was further supported by GSEA analysis on pre-defined “mid-fetal” and “early-fetal” genes from the human brain, with mid-fetal genes precociously upregulated in mutants (Fig. S3S,T). Interestingly, this is the opposite expression pattern to *Chd8* mutants, who have macrocephaly rather than microcephaly (Katayama et al., 2016). We also looked at Wnt and Notch signaling pathways, which are suppressed by MYT1L in OE studies (Mall et al., 2017). However, we found no significant categorical upregulation in our *Myt1l* mutants (Fig. S3U,V). Surprisingly, MYT1L loss also impacted cell cycle pathway genes, with inhibitors (e.g. *Rb1*, *Cdkn1c*, *Gas1*) upregulated and mitosis genes (e.g. *Mcm5*, *Cdca5*, *Ccnf*) downregulated (Fig. 3F,G, **Table S2**). In addition, an upregulation of electron transport chain (e.g. *Mt-nd2*, *Mt-co3*) and downregulation of glycolysis (e.g. *Gpi1*, *Ldha*) indicate mitochondria dysfunction upon MYT1L loss, which could further affect neuronal development (Khacho et al., 2017; Klein Gunnewiek et al., 2020). We further compared gene expression between Het and KO mice and found a further upregulation in KOs of genes associated with chromatin activation (e.g. *Setd2, Dpf3*; Fig. S3L). We speculate these changes might represent compensation for complete MYT1L loss and could explain the more open chromatin in the KOs than Hets (Fig 3A). Overall, the results suggest MYT1L mutation leads to precocious early neuronal programs and perturbs cell proliferation programs during brain development, which might contribute to microcephaly in adults.

### MYT1L loss impairs cell proliferation in developing mouse cortex

Precocious neuronal differentiation can potentially reduce the progenitor pool and thus alter cell-type proportions in the developing brain. Further, decreased progenitor numbers can reduce overall cell production and eventually result in smaller brain size. To validate our prediction that MYT1L loss affects cell differentiation and proliferation, we first stained cell-stage markers in E14 cortex (Fig. 4A) to assess the proportions of different cell populations including progenitors. We found KOs have decreased apical progenitor (SOX2+) density with normal intermediate progenitor (TBR2+) and postmitotic neuron (TBR1+) density compared to Hets and WTs (Fig. 4C-E, S4B-C). After normalizing SOX2+ cells to total cell number, there was still a trend towards fewer SOX2+ cells in KOs (Fig. S4A, p = 0.0528), indicating smaller APs density can be independent of decreased total cell number (Fig. 4B). We also found the ratio between TBR2+ and SOX2+ but not between TBR1+ and TBR2+ was increased in both Het (p = 0.066) and KO (p = 0.0026) mice (Fig. S4D,E). Overall, when considering all three cell types, we saw a shift from early cell fate (SOX2+) to late cell fate (TBR2+/TBR1+) in *Myt1l* mutants (Fig. 4F), supporting the precocious cell differentiation program hypothesized from the RNA-seq. Proliferating cells (Ki-67+) were also decreased in KOs after normalizing to total cell number cell number (Fig. 4G-I), suggesting MYT1L loss affects cell proliferation. Therefore, we performed EdU labeling experiments to measure proliferation rates (Fig. 4J). We found that, within a 1.5-hour window, both Het and KO cortices have significantly fewer EdU+ cells (Fig. 4K), highlighting a slower cell proliferation rate in the mutant developing cortex and a potential mechanism for the adult microcephaly.

**Figure 4:**
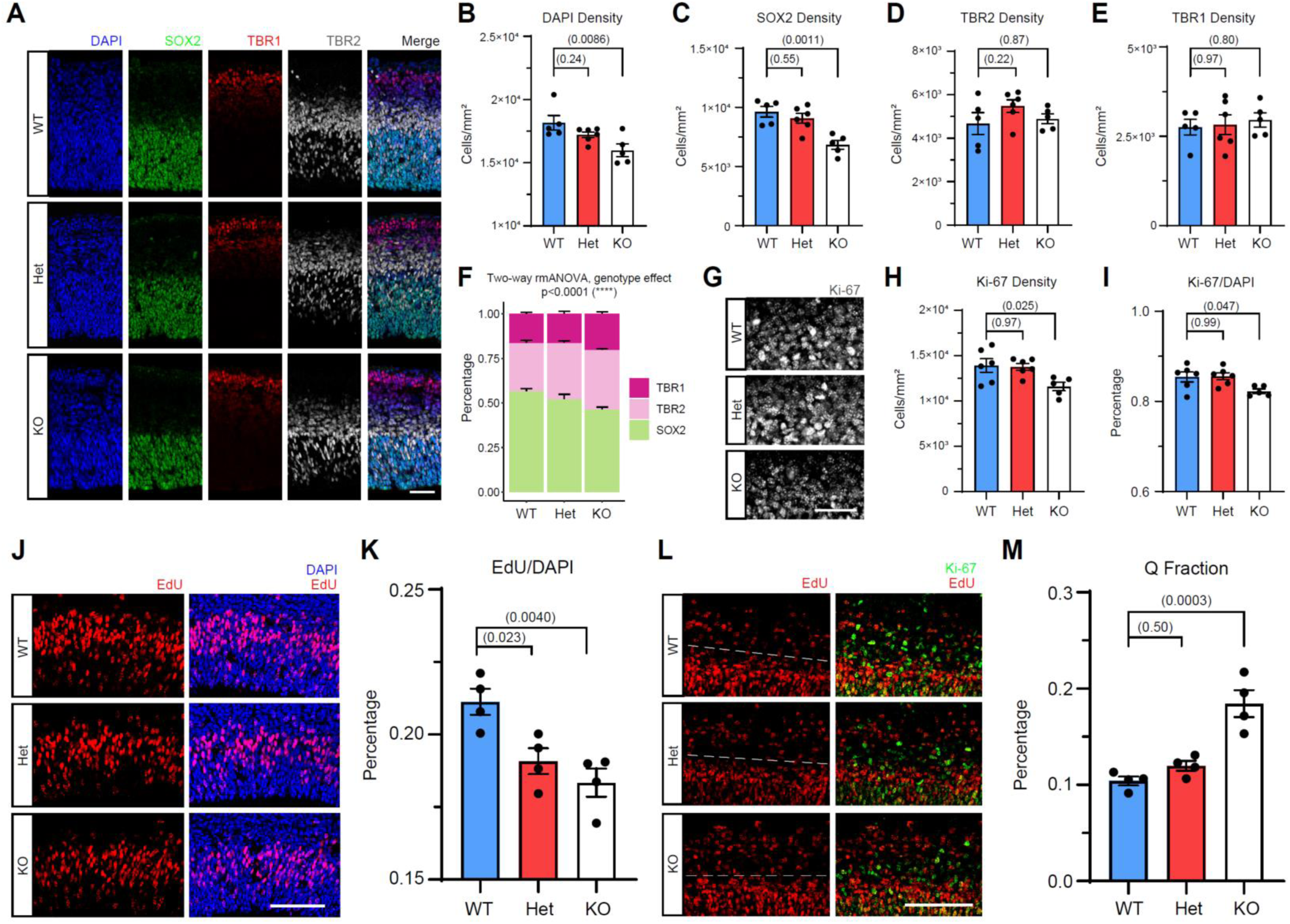
MYT1L loss disrupts progenitor proliferation by precocious cell cycle exit. **(A)** Immunostaining for nuclei (DAPI, blue), apical progenitors (SOX2, green), intermediate progenitors (TBR2, grey) and postmitotic neurons (TBR1, red) in the E14 mouse cortex. **(B)** KO mouse cortex had significantly less cellular density, **(C)** less apical progenitors, with normal **(D)** intermediate progenitors and **(E)** postmitotic neurons. **(F)** *Myt1l* mutants have significantly more early cell (SOX2+ and TBR2+) populations but less later cell (TBR1+) population. **(G-H)** KO mice have less proliferating cells compared to Het and WT littermates, **(I)** even after being normalized to total cell number. **(J)** EdU labeling for a 1.5-hour window found decreased cell proliferation rate in both Het and KO mouse cortex compared to WT, as quantified in **(K)**. **(L)** Co-staining for Ki-67 and EdU (20-hours after labeling) experiments found **(M)** a larger Q fraction value in KO but not in Het mouse cortex compared to WT. White dash lines in **(L)** indicates the border where proliferating cells started to exit the cell cycle and differentiate. *Data are represented as mean ± SEM. Scale bars, 25 μm in **A**, 50 μm in **G**, 100 μm in **J&L**. See also* Figure S4 *and* ***Table S5*** *for statistical test details*.

Following mitosis, daughter cells either re-enter cell cycle to expand the progenitor pool, or leave permanently and become neurons. Decreased proliferation could be driven by a greater number of cells exiting the cell cycle. Therefore, we quantified exiting by co-staining for recently proliferating cells (EdU 20 hours) which have lost Ki-67 (Q fraction; Fig. 4L) (Gompers et al., 2017). KOs had significantly larger Q fraction (Fig. 4M). These results show that MYT1L loss perturbs cell proliferation and enhances exit from the cell cycle. This corresponds well to the RNA-seq, and provides the most parsimonious explanation for the smaller brains: precocious differentiation of a fraction of neural progenitors results in overall less proliferating cells and a decreased final neuron number and brain size in adults.

### MYT1L haploinsufficiency results in sustained chromatin changes in adult brain

Germline mutants also enable investigation of MYT1L function in the adult brain. We next determined if the developmental molecular deficits continue, or if MYT1L serves a distinct role in adults. As ID and ASD are not well localized in the brain, we focused on the prefrontal cortex (PFC), known to be dysregulated in human ADHD (Yasumura et al., 2019). For ATAC-seq (6 WT, 6 Het PFC, Fig. S1A,B), we discovered 4988 DARs (FDR<0.05, 9756 DARs FDR<0.1; Fig. 5A,B, S5C,D, **and Table S2**), with no peak showing sex and genotype interaction (**Table S2**). Motif analysis on DARs found regions of lost accessibility in Hets are enriched for motifs of TFs involved in neuron projection (*Egr2*) and the ASD gene *Foxp1*, while those more-accessible regions had motifs for an early neuronal TFs (*Eomes*; Fig. 5C, S5F). GO analysis (**Table S4**) likewise highlighted disruption of neuronal projection development and synaptic transmission pathways (Fig. 5D, S5G). Similar to E14, CHIP-seq targets had less accessibility in adult Hets (Fig. S5E) and more ChIP targets overlapped with less-accessible DARs than those overlapped with more-accessible DARs (*χ*^2^(1, *N*=291)=143.94, *p*<.0001), again suggesting MYT1L often has a direct role as an activator *in vivo*.

**Figure 5:**
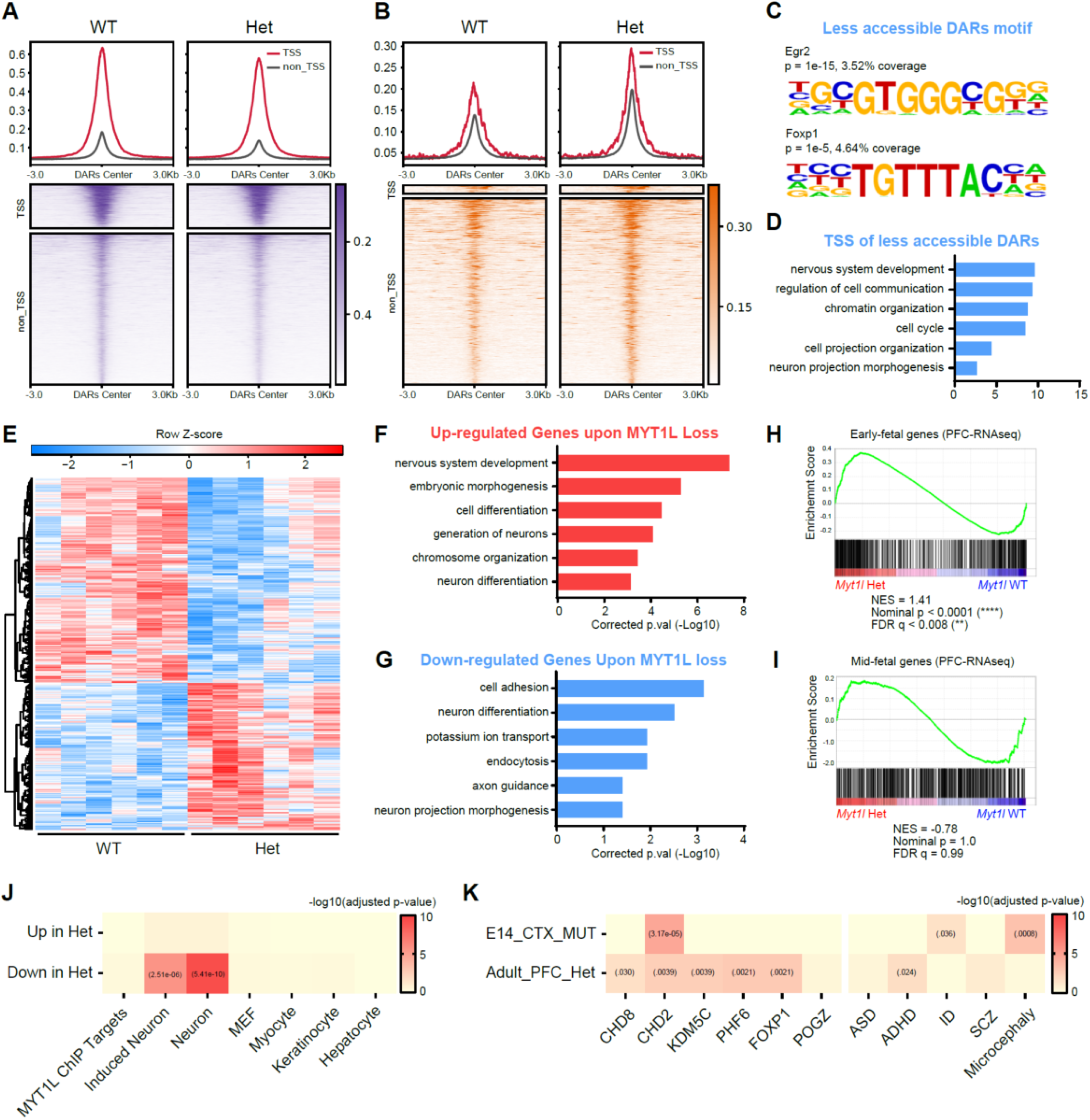
Long term MYT1L deficiency results in arrested maturation of neuronal chromatin and expression patterns. **(A)** Less and **(B)** more-accessible regions in adult Het mouse PFC identified by ATAC-seq (FDR < .1). **(C)** Homer motif analysis on less-accessible DARs over more-accessible DARs. **(D)** GO analysis on DAR associated genes showed the dysregulation of neurodevelopmental programming in adult Het mouse PFC. **(E)** Heatmap for differential gene expression in adult Het mouse PFC (FDR < .1). **(F)** GO analysis on DEGs revealed an upregulation of early neurodevelopmental pathways and **(G)** a down-regulation of neuron maturation and functions. **(H)** GSEA analysis found “early-fetal” genes increased their expression while **(I)** “mid-fetal” genes remained unchanged in adult Het mouse PFC compared to WT. **(J)** Repressed genes upon MYT1L loss in PFC significantly overlapped with induced neuron signature and neuronal signature genes. **(K)** MYT1L regulated genes were implicated in other ID/ASD mouse models and human genetic data sets. *See also* Figure S5 *and* ***Table S5*** *for statistical test details*.

We also performed RNA-seq (6 WT, 6 Het PFC, Fig. S5H) to determine transcriptional consequences. Of 14,104 measurable genes, we identify 533 DEGs in Het PFC (Fig. 5E **& Table S3**). And sex specific analysis showed limited number of genes with significant sex and genotype interaction, including genes related to activity (*Bdnf* and *Jun*) and cell adhesion (*Cdh3* and *Perp*) (**Table S3**). Mapped to ATAC-seq data, there was good correspondence between TSS accessibility and gene expression (Fig. S5I, **Table S3**). Opposite to E14 RNA-seq, CHIP-seq promoter-related genes displayed subtle *upregulation* upon MYT1L loss in adult RNA-seq (Fig. S5J), suggesting in adults it may be more often acting as a repressor (though more firm conclusions await MYT1L binding data in adult brains). When comparing DEGs’ expression between adult *in vivo* and prior *in vitro* RNA-seq, we still did not see correlation in fold changes (Fig. S5K,L). Interestingly, only upregulated genes from our *in vivo* RNA-seq significantly overlapped with ‘*in vitro* MYT1L repressed genes’ (defined as above), while our downregulated genes did not show any overlap with MYT1L induced genes (Fig. S5M,N). Also, the DEGs from E14 but not adult RNA-seq were significantly enriched in CHIP-seq targets (Fig. 5J, S3K). These results suggest that MYT1L has distinct targets in adult brain, and perhaps different roles than in E14. In E14 the preponderance of evidence fit a model of a loss of an activator. In the adult, the results were more mixed with ATAC suggesting loss of an activator, while the RNA-seq contrasted somewhat.

### MYT1L haploinsufficiency results in failed transcriptional development

To define a role of MYT1L in the adult brain, we performed GO analysis on DEGs (**Table S4**). This revealed that genes from early phases of CNS development (neurogenesis, neuronal differentiation genes, e.g. *Eomes*, *Dlx2*, *Dcx*), were up-regulated in Hets (Fig. 5F). Interestingly, these are genes expressed in immature neurons, again indicating a shift in timing of transcriptional maturation. To systematically evaluate this, we performed GSEA and confirmed increased expression of “early-fetal” genes with no expression change of “mid-fetal”, Wnt signaling, and Notch signaling genes in Hets (Fig. 5H,I, S5O,P). Persistent activation of developmental programs suggests that adult Het brains are trapped in an immature state. Indeed, genes downregulated upon MYT1L loss were significantly enriched in neuronal genes, showing an impaired mature neuronal identity (Fig. 5J). Likewise, GO analysis in Hets showed downregulation of neuronal projection development (e.g. *Epha7, L1cam)*, ion homeostasis (e.g. *Kcnt2*, *Kcne4*, *Kctd13)*, and synaptic transmission (*Gipc1, Vamp2*, Fig. 5G), echoing this immaturity, and potentially a disrupted neuronal function.

Finally, since MYT1L Syndrome is one of several forms of ID/ASD caused by TF mutation, we tested whether DEGs are dysregulated in related models. DEGs from adult RNA-seq significantly overlapped with DEGs from *Chd8*, *Chd2*, *Kdm5c*, *Phf6*, *Foxp1,* and *Pogz* KO mouse models (Fig. 5K). DEGs from E14 were enriched in the *Chd2* and *Chd8* datasets (Fig. 5K, S5Q). Interestingly, post hoc analysis showed genes were dysregulated in an opposite direction between *Myt1l* mutant mice and other ID/ASD mouse models (**Fig S5Q,R**). This suggests genes implicated in different ID/ASD models are pathogenic when dysregulated in either direction.

Further, compared to human data, DEGs derived from PFC of Het mice were enriched in ADHD and ASD associated genes, but not in human IDD, SCZ, or microcephaly genes (Fig. 5K, S5R). Conversely, DEGs from E14 significantly overlapped with human ID and microcephaly but not ASD, ADHD, or SCZ genes (Fig. 5K, S5Q). Together, these findings highlight some convergence between MYT1L Syndrome and other developmental disorders.

### MYT1L haploinsufficiency disrupts postnatal neuronal physiology and spine maturity

Het mice showed deficits in transcriptional and epigenetic chromatin states, with a failure to achieve the mature profile alternation in axonal development programming. Therefore, we asked whether this manifested in neurophysiological changes at the level of cellular excitability or synaptic transmission. We first examined the passive membrane properties and cell morphology of layer 2/3 pyramidal neurons in the primary visual cortex (V1) at P21-23, an extensively studied system with similar cell types and mesoscale circuit connectivity to PFC. Early postnatal development drives a series of changes in synaptic and membrane properties of cortical neurons, which are collectively necessary for normal function(Desai et al., 2002; Kasper et al., 1994; Kroon et al., 2019; Maravall et al., 2004). Compared to age-matched WT neurons, Het neurons exhibited significantly depolarized resting membrane potentials (Fig. 6A), and significantly decreased membrane resistance (Fig. 6B), changes that affect membrane excitability in opposite directions. We also observed a smaller time constant in Hets that was explained by the decrease in membrane resistance and capacitance (Fig. 6C,D), which could arise from a decrease in total cell surface area or altered ion channel composition. In total, MYT1L haploinsufficiency disrupts the passive physiological properties of pyramidal neurons. To ask whether the change in capacitance was a direct result of cell surface area, we examined the somatic size of the patched neurons. A previous shRNA study on differentiating NPCs revealed larger cell bodies yet decreased neurites (Kepa et al., 2017). Here, with controlled haploinsufficiency *in vivo*, we found that MYT1L loss changed neither neuron soma size (Fig. 6E, S6A,B) nor dendrite morphological properties, including length, nodes, as well as complexity (Fig. S6D-G). There was a small decrease of total dendrite numbers in Het neurons (Fig. S6C), but it was not significant (p = 0.054). Further branch analysis revealed no difference between Hets and WTs for branch order numbers or length (Fig. S6H,I). To assess detailed dendritic complexity, we conducted a Sholl analysis and still found no difference in spatial aspects of dendritic morphology across genotypes (Fig. 6F, S6J). These results indicate altered passive properties of Het neurons is not caused by morphological changes.

**Figure 6:**
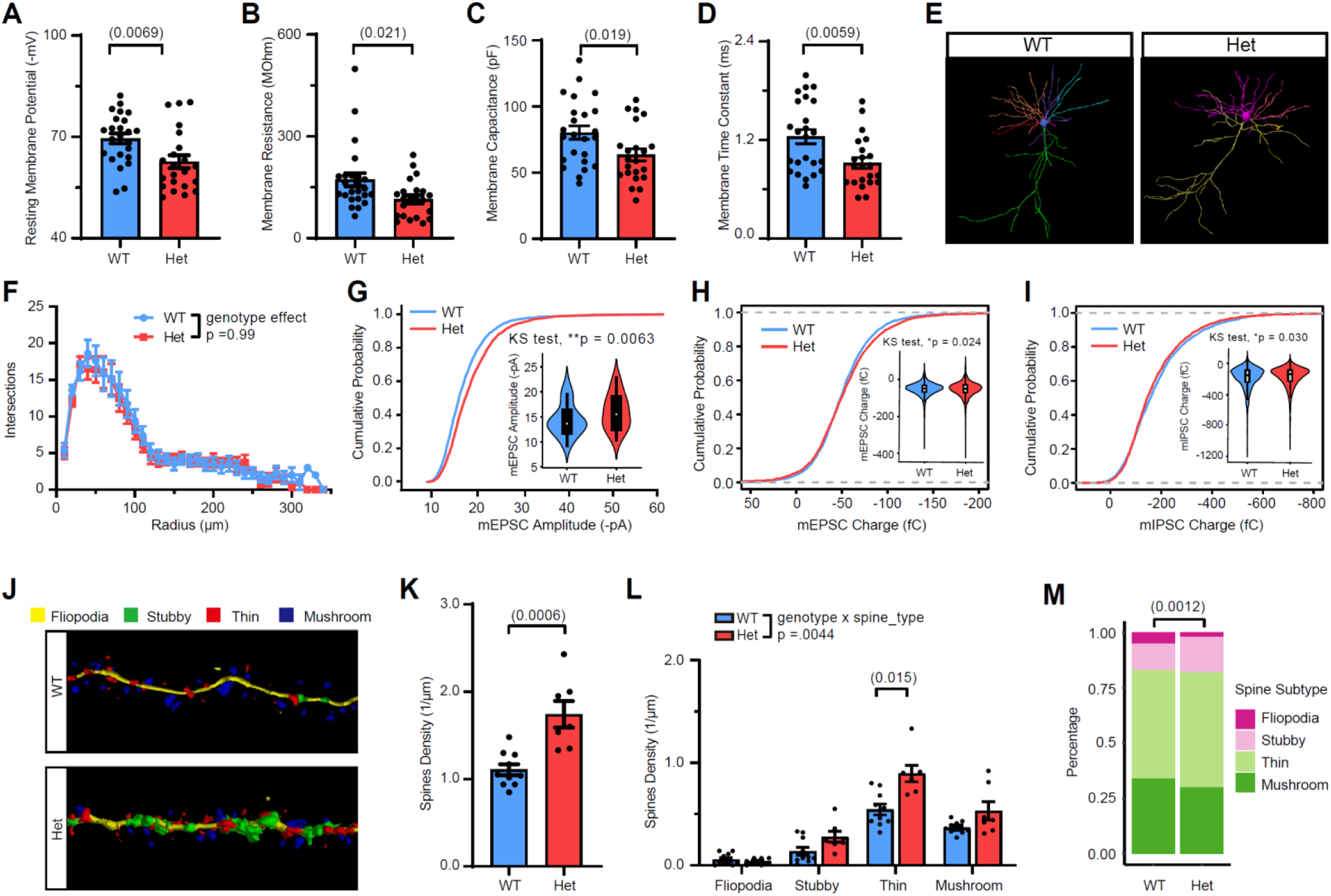
MYT1L haploinsufficiency disrupts baseline neuronal properties and dendritic spine maturity but not neuronal morphology. **(A)** MYT1L loss led to less negative membrane potential, **(B)** reduced membrane resistance, **(C)** decreased membrane capacitance, and **(D)** smaller membrane time constant in cortical pyramidal neurons. **(E)** Neuronal soma and dendrites tracing in Neurolucida. **(F)** Sholl analysis found no dendrite complexity change across genotypes. **(G)** Het neurons showed increased mEPSC amplitudes distribution compared to WT neurons. **(H)** Analysis on individual events of mEPSC and mIPSC found that the charges of Het neurons’ mEPSC are slightly larger, **(I)** while mIPSC are slightly smaller. **(J)** Representative images of spine tracing and subtypes identification using Neurolucida. **(K)** Het neurons had more apical spines compared to WT neurons with **(L)** general increase in different spine subtypes. **(M)** Het neurons had a higher percentage of less immature spines (Stubby, Thin) and less mature spines (Mushroom) compared to WT. *Data are represented as mean ± SEM. See also* Figure S6 *and* ***Table S5*** *for statistical test details*.

We next asked whether MYT1L haploinsufficiency affects the number or strength of synapses onto cortical pyramidal neurons. To do this, we measured miniature excitatory postsynaptic currents (mEPSCs, Fig. S6K). We saw no change in the frequency of mEPSCs(Fig. S6L). However, we did see a trend towards an increase in the mean amplitude of the mEPSCs across cells (Fig. S6M). More immature cortical neurons have been shown to have larger mEPSCs (Desai et al., 2002). Investigating all individual mEPSC events revealed they were indeed shifted towards larger currents (Fig. 6G). Since excitation/inhibition (E/I) balance is often disrupted in neurodevelopmental disorders including ASD (Gogolla et al., 2009; Nelson and Valakh, 2015), we also measured miniature inhibitory postsynaptic currents (mIPSC, Fig. S6N) to examine E/I balance in Het mice. With no change in mIPSC amplitude, there was a small decrease of mIPSC frequency, though not significant (p = 0.081)(Fig. S6O,P). We further looked at mEPSC and mIPSC charge and found that the distribution of charge carried by all individual postsynaptic current events shift towards increased excitation (p=.024) and decreased inhibition (p=.030) in Het neurons compared to WT (Fig. 6H,I). These results suggest MYT1L loss leads to increased E/I ratio in the mouse brain. Morphologically, microscopic investigation of apical dendritic spine density and morphological maturity (Fig. 6J) revealed increased spine density (Fig. 6K) with decreased mature spines(mushroom) but increased immature spines (thin and stubby) in Het neurons (Fig. 6L,M). Neurons tend to generate excessive spines during early development and spine numbers decrease via pruning process after postnatal development (Bhatt et al., 2009). Thus, increased spine density again indicated disrupted maturation of Het neurons. However, we did not see mEPSC frequency increase in Het neurons, suggesting extra spines were non-functional or other compensatory mechanisms offset the impact of increased spine density.

### MYT1L haploinsufficiency persistently impairs muscle strength and endurance, and elevates activity and arousal

We determined behavioral circuit consequences of the sustained molecular anomalies resulting from MYT1L haploinsufficiency. We evaluated Hets for features related to developmental delays, ADHD, ASD, and ID present in human MYT1L deletion patients and our index patient by conducting a comprehensive behavioral characterization (Fig. 7A,B). We note that the index patient was a male who had been diagnosed with ASD in early childhood; he exhibited sustained pathognomonic features of the condition including repetitive thinking, subtle stereotypic motor mannerisms, deficiency in eye gaze, and interpersonal aloofness when focused on the objects of his own mental pursuits, fully consistent with DSM5 level 1 severity of impairment in function. These ASD symptoms were out of proportion to social impairments that would be attributable to the comorbid conditions of ADHD and mild ID with which he was also diagnosed, and he displayed the compensatory strength of a distinct affability (at times to the point of joviality) and enjoyment of social interaction despite the quality and consistency of social interaction being compromised by his ASD symptoms. Further details of his symptom profile are provided in **Table S1**.

**Figure 7:**
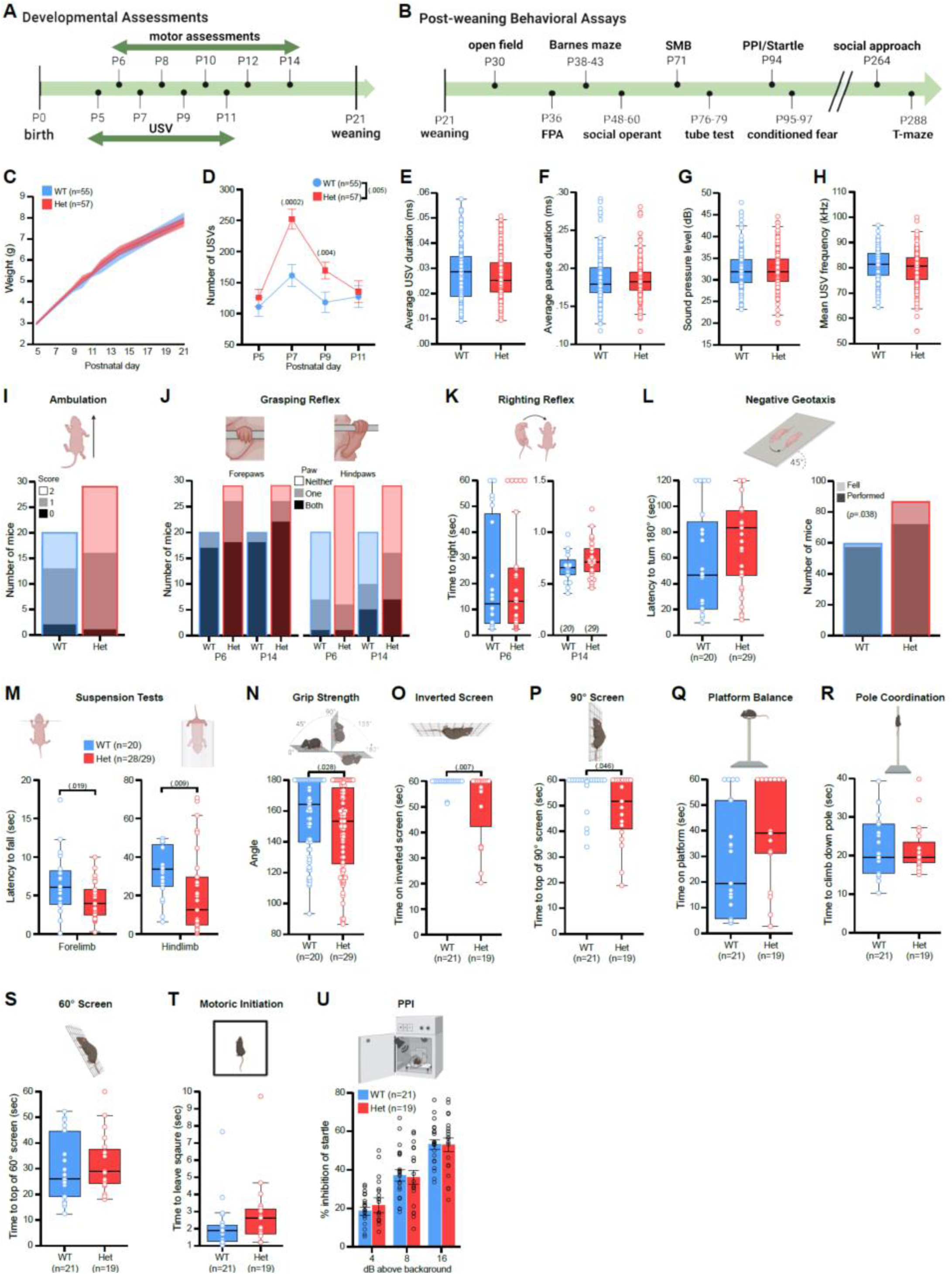
*Myt1l* haploinsufficiency results in heightened USV production and muscle weakness and fatigue. **(A)** Timeline for developmental assessments. **(B)** Timeline for post-weaning behavioral assays. **(C)** Early postnatal weight trajectories were comparable for WT and Hets. **(D)** Hets produced fewer USVs from P5-P11 than WT. **(E-H)** Het calls did not differ from WT calls on temporal (call duration **(E)**, pause duration **(F)**, sound pressure level **(G)**) or spectral (mean frequency **(H)**) features. **(I)** MYT1L loss was not associated with ambulation scores at P8. **(J)** MYT1L loss was not associated with grasping reflex in the forepaws or hindpaws. **(K)** Hets exhibited latency to righting reflex similar to WTs at P6 and P14. **(L)** Latency to exhibit negative geotaxis was comparable between Hets and WTs at P10, however, MYT1L loss was associated with increased falls from the inclined apparatus. **(M)** Hets were unable to remain suspended by fore or hindlimbs as long as WTs. **(N)** Hets fell from the grip strength mesh screen at a narrower angle than WTs. **(O)** As adults, Hets hung on an inverted screen for a shorter latency than WTs. **(P)** Hets exhibited a longer latency than WTs to climb to the top of a 90° screen. **(Q)** Time to balance on an elevated platform was similar between Hets and WTs. **(R)** Hets climbed down a pole at a comparable latency to WTs. **(S)** Hets climbed to the top of a 60° wire mesh screen at a comparable latency to WTs. **(T)** Hets initiated movement at a similar latency to WTs. **(U)** Percent inhibition of startle following a pre-pulse was similar in Hets and WTs. *For panels C, D, and U, grouped data are presented as means ± SEM. For panels E-H, K, L (left), and M-T grouped data are presented as boxplots with thick horizontal lines respective group medians, boxes 25^th^ – 75^th^ percentiles, and whiskers 1.5 x IQR. Individual data points are open circles. See also* ***Table S5*** *for statistical test details*.

Language and motor delay are universal in MYT1L deletion patients (Blanchet et al., 2017), therefore, we assessed Hets for gross developmental, communication, and motor delay (Fig. 7A). Physically, we found that Hets did not exhibit signs of gross developmental delay: they matched WT ages at pinnae detachment and eye opening, and postnatal weight gain (Fig. 7C). We examined early communicative interaction by recording ultrasonic vocalizations (USV) emitted by isolated pups. Hets exhibited an increase in USV rate compared to WT littermates (Fig. 7D) following maternal separation, that is likely independent of altered respiratory muscle function (Fig. 7E-H). Rather than delayed communicative behavior, this elevated rate suggests an anxiety-like phenotype or, since USV rate also reflects arousal levels, a heightened arousal that may reflect a hyperactive phenotype.

Possible motor delay was assessed with a battery of tasks conducted during the first two weeks postnatal (Feather-Schussler and Ferguson, 2016), which examined the acquisition of motor function, including ambulation, posture, reflexes, and muscle strength and endurance. Hets exhibited normal acquisition of ambulation, grasping reflex (Fig. 7I,J), and comparable latencies for righting and negative geotaxis reflexes (Fig. 7K,L). However, *Myt1l* mutation was associated with an inability to hold position during the negative geotaxis test (Fig. 7L), indicating Hets had a difficult time holding themselves in place. Hets were also unable to remain suspended as long on other strength tasks including fore- and hindlimb suspension (**Fig. N**), and grip strength (Fig. 7O) compared to WTs. While these tasks are not exhaustive, the results suggest no gross motor delay was present. Yet, the strength and endurance deficits suggest hypotonia, a feature reported often in MYT1L deletion patients (Blanchet et al., 2017; Doco-Fenzy et al., 2014; Windheuser et al., 2020b).

In an independent cohort assessed from P30 through adulthood (Fig. 7B) we also observed phenotypes consistent with reduced muscle strength and endurance in Hets on sensorimotor tasks. Hets demonstrated decreased strength and endurance on the inverted screen test (Fig. 7P) and difficulty climbing a 90° wire screen (Fig. 7Q), which requires strength and coordination. Hets were largely normal on the remaining sensorimotor tasks for balance, coordination and movement initiation (Fig. 7R-U). In addition, we found comparable pre-pulse inhibition (PPI) between groups in the sensory gating startle/PPI task (Fig. 7V). Coupled with the neonatal data, these findings indicate MYT1L loss resulted in muscle weakness suggestive of hypotonia, yet future studies of body composition and muscle pathology will be necessary to confirm this as a model of MYT1L-dependent hypotonia.

As patients show ID, we examined spatial learning and memory and Pavlovian fear conditioning as assessments of learning in mice. Hets displayed normal spatial acquisition and memory retention in the Barnes maze (**Fig S7A,B**). However, Hets failed to show typical contextual and cued fear conditioning (Fig. S7C), suggesting decreased associative memory. In the same cohort of mice, we examined activity levels for ADHD-like features at P30. Regardless of sex, Hets were hyperactive in the open-field task, traveling a greater distance than WT littermates (Fig. S8A). This hyperactivity replicated in subsequent assays, where activity variables were also available: in distance traveled in the social operant task and in heightened baseline force measurements in the startle task (a measurement of movement in the apparatus in the absence of startle stimuli; Fig 8S, S8B). This hyperactive phenotype confounds the interpretation of the conditioning data above because it can mirror a conditioning deficit in this task. Thus, further investigations are necessary to understand any learning deficits in this model. Finally, we assessed the center variables of the open field task for anxiety-related behaviors (thigmotaxis), and found no increase in anxiety-related behavior in Hets as measured in this task (Fig. S8C). The hyperactivity phenotype in the absence of anxiety-related markers sheds more light on the heightened USV data. It further supports our interpretation that we were observing an arousal-related increase in call rate during the first two weeks postnatal.

**Figure 8.**
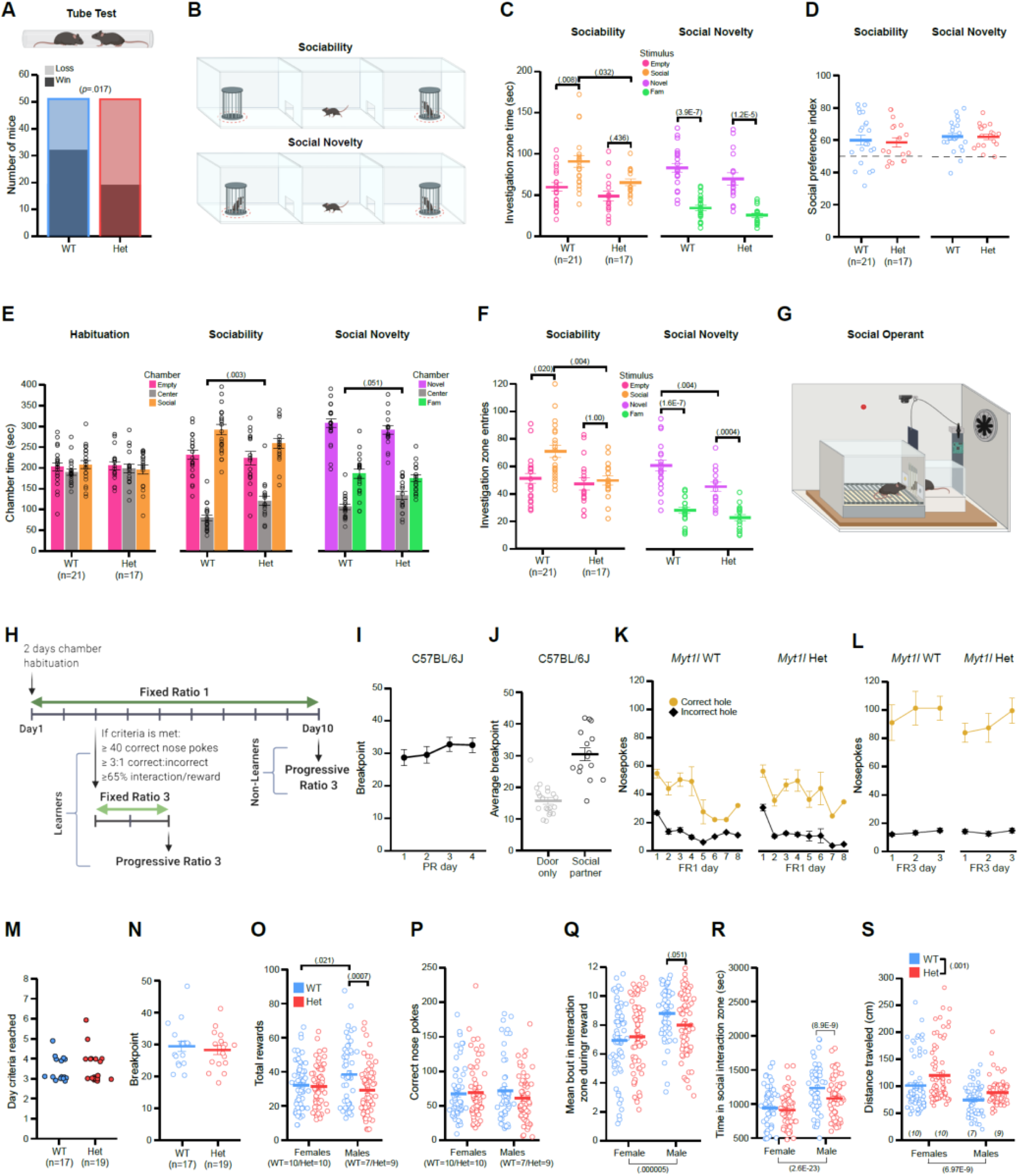
*Myt1l* haploinsufficiency altered social behaviors. **(A)** MYT1L loss was associated with more losses in the social dominance assay. **(B)** Assay schematics for social approach test trials. Investigation zones demarcated by the dotted red lines. **(C)** During the sociability trial, Hets spent less time investigating with the social stimulus than WTs and failed to show an increase in time spent in the social versus empty investigation zones. No difference between genotypes was observed in social novelty. **(D)** Sociability and social novelty preference scores were comparable between Hets and WTs. **(E)** Hets spent more time in the center chamber during both the sociability and social novelty trials of the social approach task compared to WTs. **(F)** During the sociability trial, Hets entered the zone surrounding the social stimulus fewer times compared to WTs, and failed to show an increase in entries into the social cup zone versus empty cup zone. During the social novelty trial, Hets entered the zone surrounding the novel mouse less than WT mice. **(G-H)** Social operant assay and timeline schematics. **(I)** C57BL/6J mice show consistency across multiple test days in the max level of effort they will exert for access to social interaction reward, demonstrating performance in the social operant test is reproducible in the same individuals across test days. **(J)** This max effort is driven by the social aspect of the reward as demonstrated by the difference in performance between mice that receive the social interaction reward versus mice that received only the raising of the door yet no access to a stimulus mouse. **(K)** The time series of task acquisition demonstrates that *Myt1l* WT and Het mice learn to discriminate between correct versus incorrect holes for access to a social interaction reward during FR1 training. **(L)** Both *Myt1l* WT and Het mice that meet learning criteria are motivated to work harder for the social interaction reward when more effort is required during the FR3 testing. **(M)** Day to reach criteria during social operant training was not different between Hets and WTs. **(N)** Breakpoint reached during social operant PR3 testing was not different between Hets and WTs. **(O)** Het males achieved less social rewards compared to WT males. **(P)** Het males and females exhibited a comparable number of correct nosepokes to WT littermates. **(Q)** During a reward, Het males trended towards less total time in the social interaction zone compared to WT males. Regardless of genotype, males spent more time in the social interaction zone compared to females. **(R)** Het males spent less total time in the social interaction zone than WT males. Regardless of genotype, males spent more time in the social interaction zone compared to females. **(S)** Female and male Hets traveled farther distances during 1-hr social operant trials compared to WTs. Overall, females traveled farther distances than males during social operant trials. *For panels C-F, I-L, and N-S, grouped data are presented as means ± SEM. Individual data points are open circles. See also* Figure S7, 8 *and* ***Table S5*** *for statistical test details*.

### MYT1L haploinsufficiency results in ASD-related social impairments particularly robust in males

We also investigated multiple behaviors related to the index patient’s ASD diagnosis. First, we investigated cognitive inflexibility, sensory sensitivity, repetitive behaviors and stereotypies across multiple assays. In the spontaneous alternation T-maze, male and female Hets exhibited comparable percent alternation to WTs and alternation rates different from chance (50%; Fig. S8D), indicating no preservation in this task. To assess sensory sensitivities, we quantified responses to stimulation of the plantar surface of the paw with von Frey filaments. Het mice exhibited an overall reduced sensitivity to this tactile stimulation (Fig. S8E). Examination of open field movement plots revealed sharp vertical movements in the perimeter, suggestive of jumping. Therefore, we re-analyzed the video-data with cutting-edge pose estimation software DeepLabCut (Mathis et al., 2018) coupled with SimBA (Nilsson et al., 2020) (Fig. S8F) to generate supervised machine-learning behavioral predictive classifiers for automated quantification of jumping behavior (**Movie S1**). Despite hyperactivity displayed by both male and female Hets (Fig. S8A), female Hets alone exhibited significantly more jumping compared to female WTs and male Hets (Fig. S8G). Therefore, this may be a female-specific overactivity trait. We also did not observe stereotyped behavior in the force-plate actometer (FPA) in the form of bouts of low mobility or movement during those bouts (Fig. S8H,I), which would be suggestive of repetitive grooming. Indeed, training a video classifier to specifically assess grooming in the open field task revealed that while there was an interesting sex difference in duration of grooming bouts (Fig. S8J), *Myt1l* mutation did not further modulate this behavior (Fig. S8K). Thus in the tasks used here, no behaviors related to repetitive/restrictive interests or stereotypies were observed.

Previous work suggested MYT1L promotes differentiation of oligodendroglia (Shi et al., 2018), which could impact myelination. Demyelination can result in a tremor in mice, as assessed by the FPA (Li et al., 2019). However, we did not observe any tremor in Hets at five weeks postnatal (Fig. S8L), suggesting the white matter anomalies we see do not reflect demyelination, consistent with the normal FA values (Fig. 3H).

Finally, we assayed multiple aspects of social behavior. To assess social hierarchy behavior, we used the social dominance tube test in which the dominant mouse will force a partner out of an interaction tube. MYT1L loss was associated with submission in this test (Fig. 8A). In the social approach task (Fig. 8B), Hets showed reduced sociability (less time investigating the novel conspecific compared to WTs) during both trials (Fig. 8C), though still exhibiting social preference (Fig. 8D). This is due to reduced investigation time overall, as Hets spent more time in the center chamber (Fig. 8E). These findings, coupled with reduced entries into the social investigation zone (Fig. 8F), indicate reduced sociability in the Hets compared to WTs.

Deficits in sociability may be due to reduced motivation to engage with a social partner. Social motivation requires both social reward circuits and social orienting circuits (i.e., attending to a social stimulus when presented) (Chevallier et al., 2012). Therefore, we used data from an adapted and extended social paradigm (Martin and Iceberg, 2015), to understand the effect of MYT1L loss on social motivation directly and parse these two possibilities. We adapted standard operant conditioning (Fig. 8G,H) to assess social motivation by rewarding nosepokes with an opportunity for transient social interaction (Fig. 8I-L). Social reward seeking is quantified by increasing the number of nosepokes required (work) to elicit each reward, and in parallel the animal’s social orienting can be assessed by tracking its behavior. Hets were normal on learning the task, including day to reach criteria based on correct to incorrect nosepokes (Fig. 8M) and appeared to show normal social reward-seeking defined by the maximum number (breakpoint) of correct nosepokes made for a reward (Fig. 8N). However, during training male Hets achieved fewer social rewards compared to WT males (Fig. 8O) despite exhibiting a comparable number of correct nosepokes (Fig. 8P). This suggested the Het males continued to poke despite the presentation of a social reward. Indeed, we found that Het males tended to spend less time at the door during a reward (Fig. 8Q), and showed a significant decrease in overall time in the interaction zone (Fig. 8R). This reduction is not secondary to increased activity levels of male Hets as both males and female Hets show increased distance traveled (Fig. 8S). Together, these data indicate that Het males failed to cease holepoking and attend to a social stimulus at the WT rate. This suggests MYT1L loss might lead to ASD phenotypes through disrupting social orienting, possibly linked to inappropriate perseveration on non-social stimuli.

## Discussion

Here, we generated a model of *Myt1l* Het mutation to address the role of MYT1L protein during CNS development, and to comprehensively characterize a model of this ID-associated syndrome. We confirmed that the frameshift mutation results in haploinsufficiency, ruling out a truncated protein mechanism. The lowered protein level leads to physical and behavioral anomalies, many of which reflect observations in patients, including microcephaly, thinned white-matter, muscle weakness, obesity, hyperactivity, and social deficits. This indicates these mice are a robust model of the disorder, and will enable preclinical and mechanistic studies that are not possible in humans.

Along these lines, molecular and neuropathologic studies defined a mechanism for aspects of the syndrome. Specifically, the syndrome’s microcephaly appears to be due to an increased rate of cell cycle exit and precocious differentiation from progenitor to immature neurons. The most parsimonious interpretation is that loss of proliferating progenitors results in insufficient expansion progenitor pools and thus a correspondingly smaller brain.

These same molecular studies clarify the role of MYT1L protein levels in normal brain development. In both Het and KOs, ATACseq revealed substantial change in chromatin accessibility across the genome, with both increases and decreases apparent. Given the shift in cell proportions from precocious differentiation, this represents a mix of direct and indirect effects. Focusing on the likely direct effects (i.e., at CHIPseq peaks), mutants showed a disproportionate loss of accessibility, suggesting MYT1L more often functions as an activator *in vivo*. Our RNAseq findings mirror these observations.

A role primarily as an activator during normal brain development agrees with some prior data, but does contrast with the specific role proposed for MYT1L during transdifferentiation studies. Prior studies defined both N-terminal activating domains and repressive domains (Manukyan et al., 2018) suggesting MYT1L may have distinct functions in different contexts. Further, the lack of binding motifs near activated transcripts following MYT1L overexpression led Manukyan et al. to speculate MYT1L’s activating effects involved either a novel motif or indirect recruitment via other TFs. Our data offer some support for the latter conclusion, with ∼20% each of reduced accessibility regions showing ASCL1 and LHX motifs, but no enrichment of the MYT1L motif. We also saw some evidence of repressive function for MYT1L, as some regions opened chromatin upon its loss. However, our findings *in vivo* during development contrast with the role MYT1L was proposed to serve *in vitro* during directed transdifferentiation of fibroblasts to neurons (Mall et al., 2017), where overexpression MYT1L corresponded to a loss of fibroblast gene expression. They concluded MYT1L served as a novel ‘repressor of all lineages save neurons,’ almost as an opposite to the classic REST TF, known to repress the expression of neuron-specific genes in all non-neuronal cells (Chong et al., 1995). We reasoned that if MYT1L had this role *in vivo*, deletion should lead to enhanced non-neuronal gene expression. However, decrease or loss of MYT1L did not result in ectopic expression of other lineages’ genes in E14 brain, suggesting such a role is not a major function during normal brain development.

Yet, with regards to later function on neuronal maturation our adult studies agree in a general way with prior shRNA data in primary neurons & NPCs (Kepa et al., 2017; Mall et al., 2017) that decreasing MYT1L levels disrupts neuronal maturation. Like these studies, we saw a decrease in mature neuronal markers, and we highlight an aberrant higher expression of immature neuronal markers such as *Eomes* and *Dlx2*. Correspondingly, Het neurons exhibited markedly abnormal passive membrane properties, specifically a depolarized resting potential, decreased membrane resistance, decreased capacitance and smaller time constant. In addition, we observed excessive dendritic spines with immature morphology and increased mEPSC amplitudes in Hets. This physiological effect was not as severe as was seen following shRNA from Mall, where action potentials were completely lost, nor Kepa, where cell body size was doubled and neurites decreased by half. Perhaps these more robust effects reflect a stronger knockdown (e.g, 90% for Kepa), and may explain why KO mice are not viable after birth. Nonetheless, synaptic and membrane dynamics are key determinants of neuronal computation, thus the changes observed *in vivo* indicate a functional mechanism by which MYT1L haploinsufficiency-induced changes in transcription and chromatin state may undermine circuit function in Hets.

This has lasting behavioral consequences as well, including muscle weakness, hyperactivity and social deficits, echoing patient prevalence of hypotonia, and the diagnosis in a subset of ADHD and ASD (Blanchet et al., 2017). Mutants were hyperactive across numerous tasks, including open field, social operant and prepulse inhibition/startle, and arguably USVs. The mice also had altered sociality, shown in the standard social approach task where they spent a decreased amount of time with stimulus mice, but had normal preference compared to an object. A common theory of ASD posits social motivation deficits are secondary either to deficits in social reward seeking or social orienting (Chevallier et al., 2012). We therefore adapted a protocol to specifically parse these possibilities: we coupled social operant conditioning to behavioral tracking and found that mutants, specifically males, learned to holepoke for a social reward, but tended to continue hole poking rather than reorienting to the social stimuli. This finding suggests this mutation might impact social orienting rather than social reward: a hypothesis that may be interesting to test with eye tracking in patients as well. Regardless, we believe this adaptation of the social operant protocol may be of use in subtyping deficits leading to social anomalies across different genetic models of ID with partially penetrant ASD.

Beyond mechanisms for the structural anomalies, the development of this new MYT1L Syndrome model will allow identification of molecular mechanisms mediating these behavioral anomalies as well. Of particular interest is understanding whether MYT1L acts on the same or different targets across CNS development. In addition, there is also an opportunity to define circuits involved in social orienting in mice, a relatively understudied area. Finally, the robust patient-related phenotypes allow for well-powered preclinical studies of potential therapeutics for MYT1L Syndrome.

## STAR * Methods

### Key Resources Table

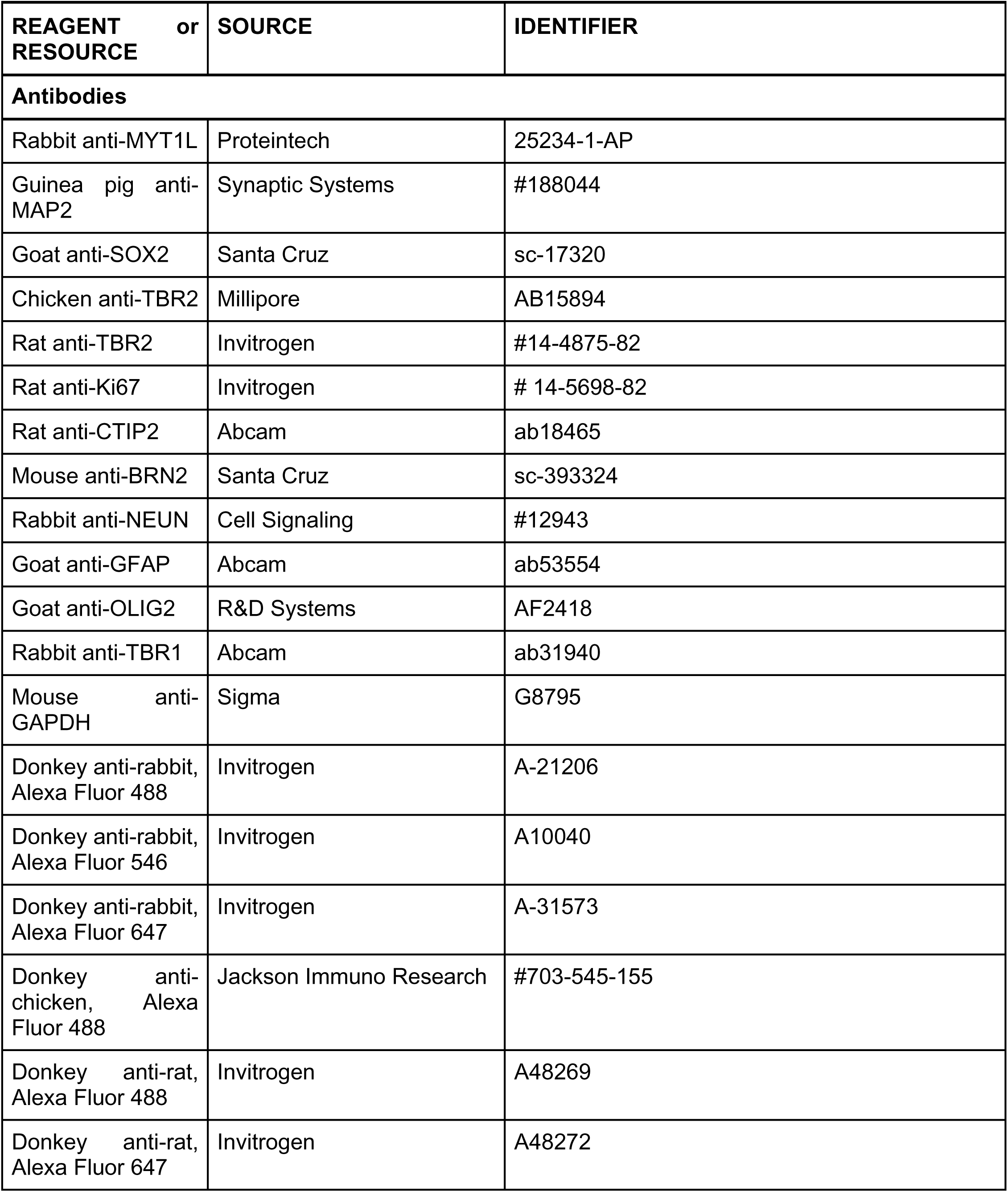
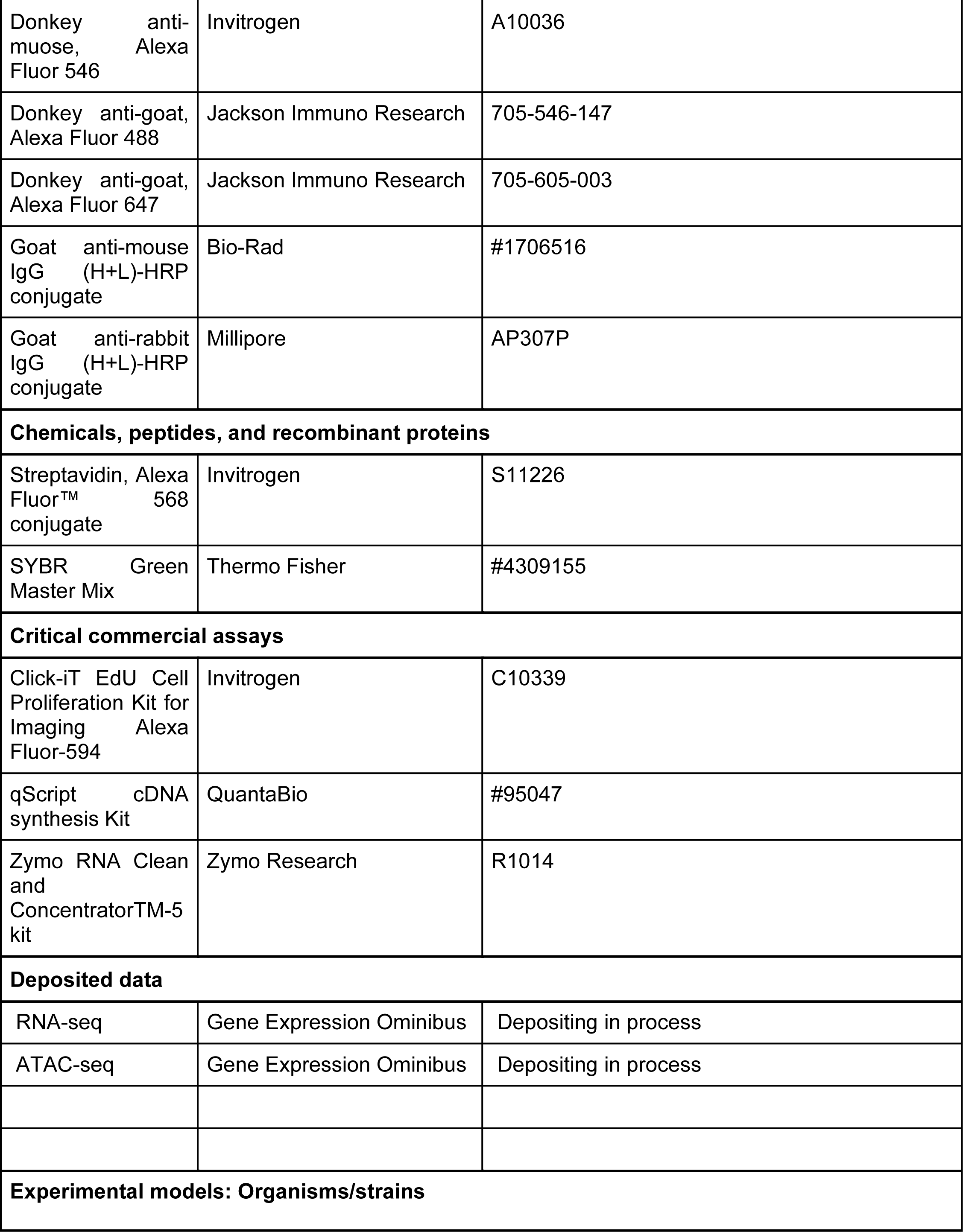
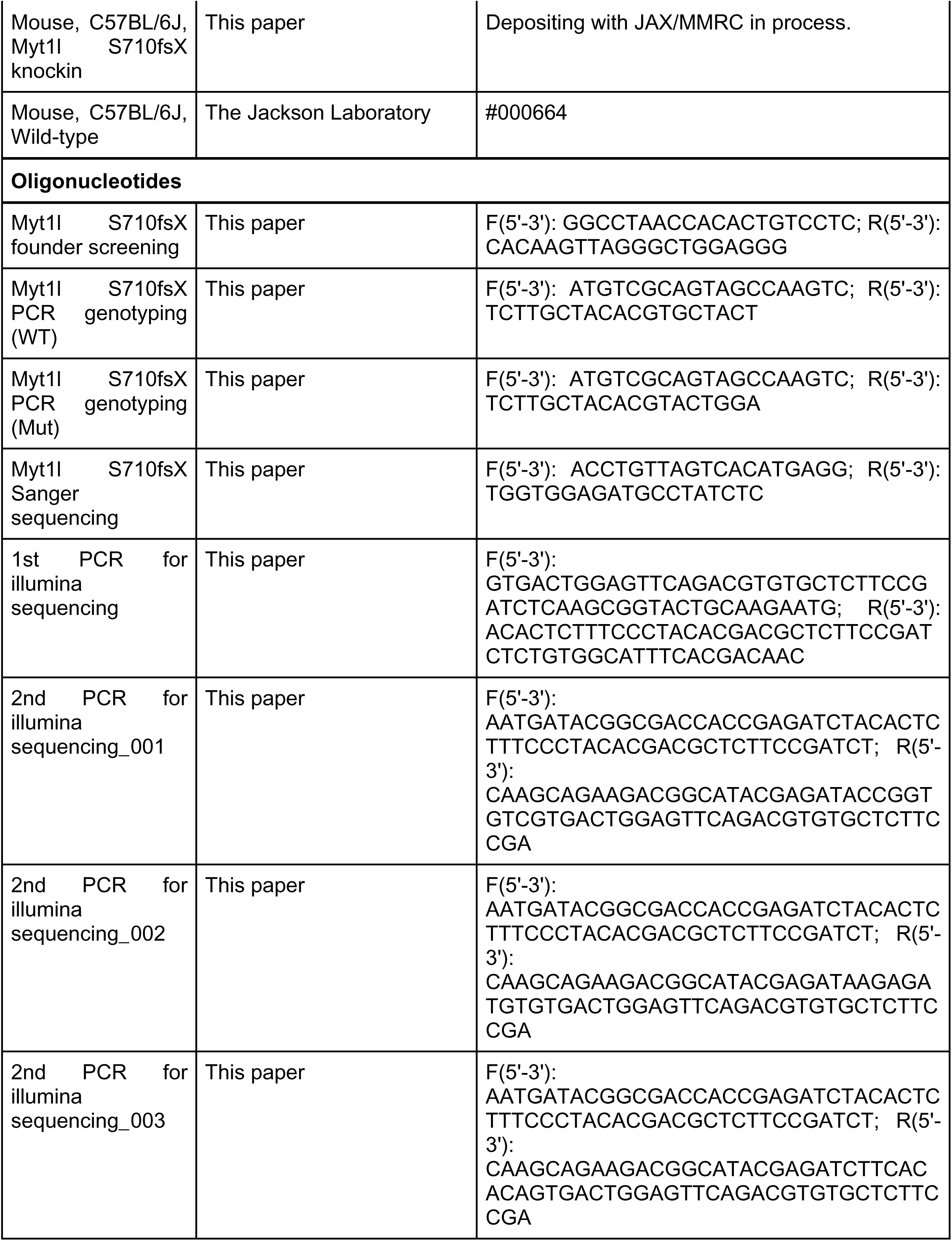
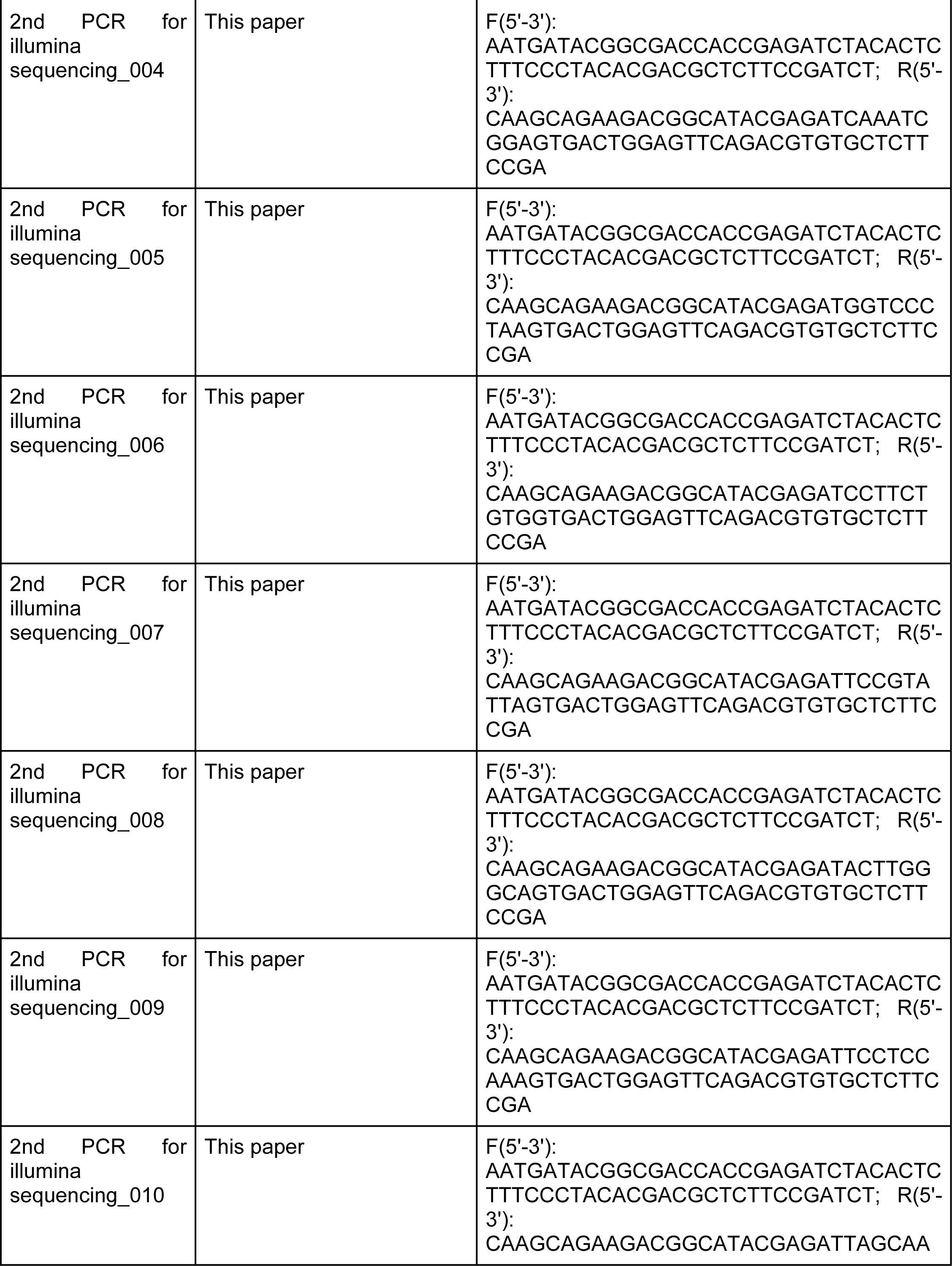
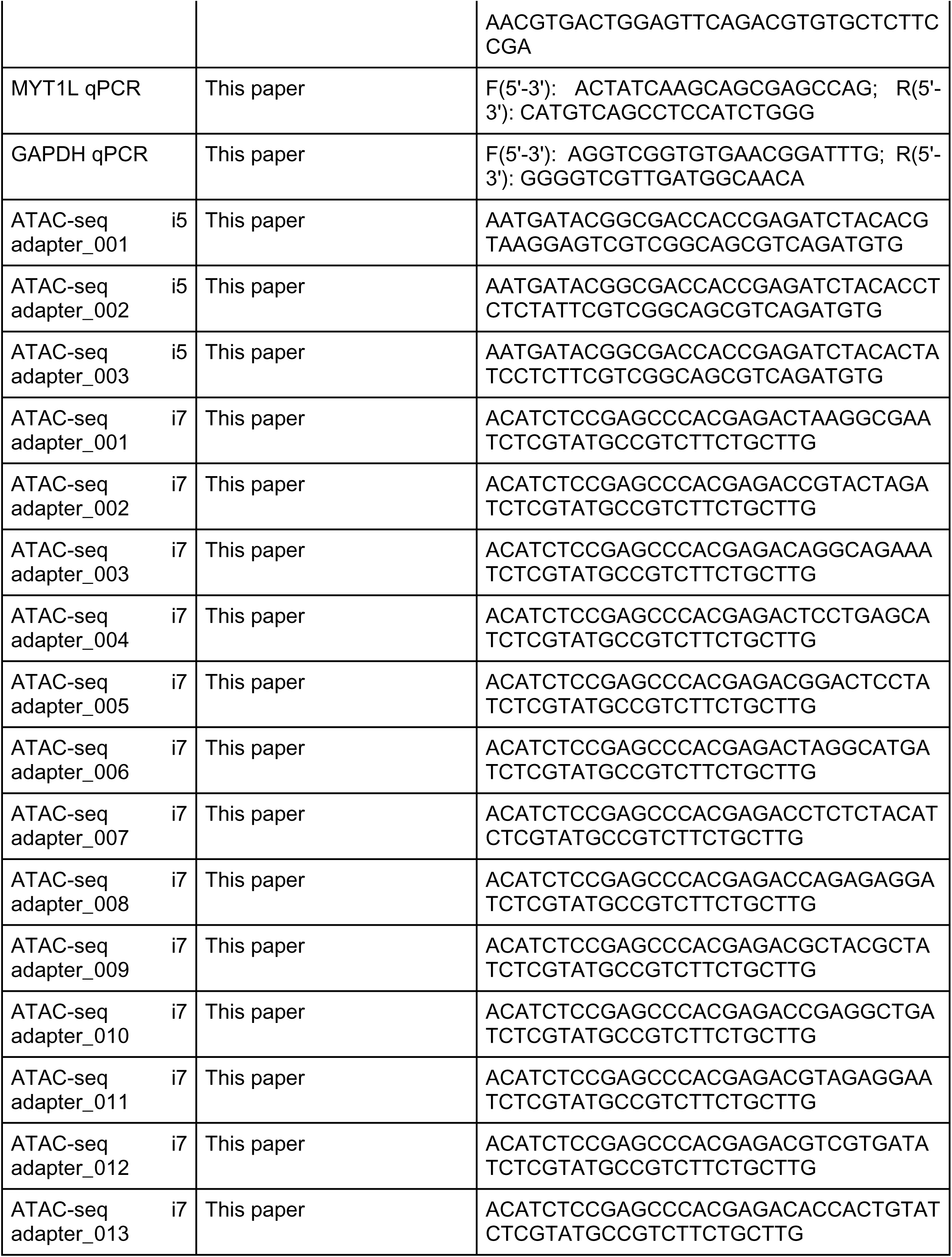
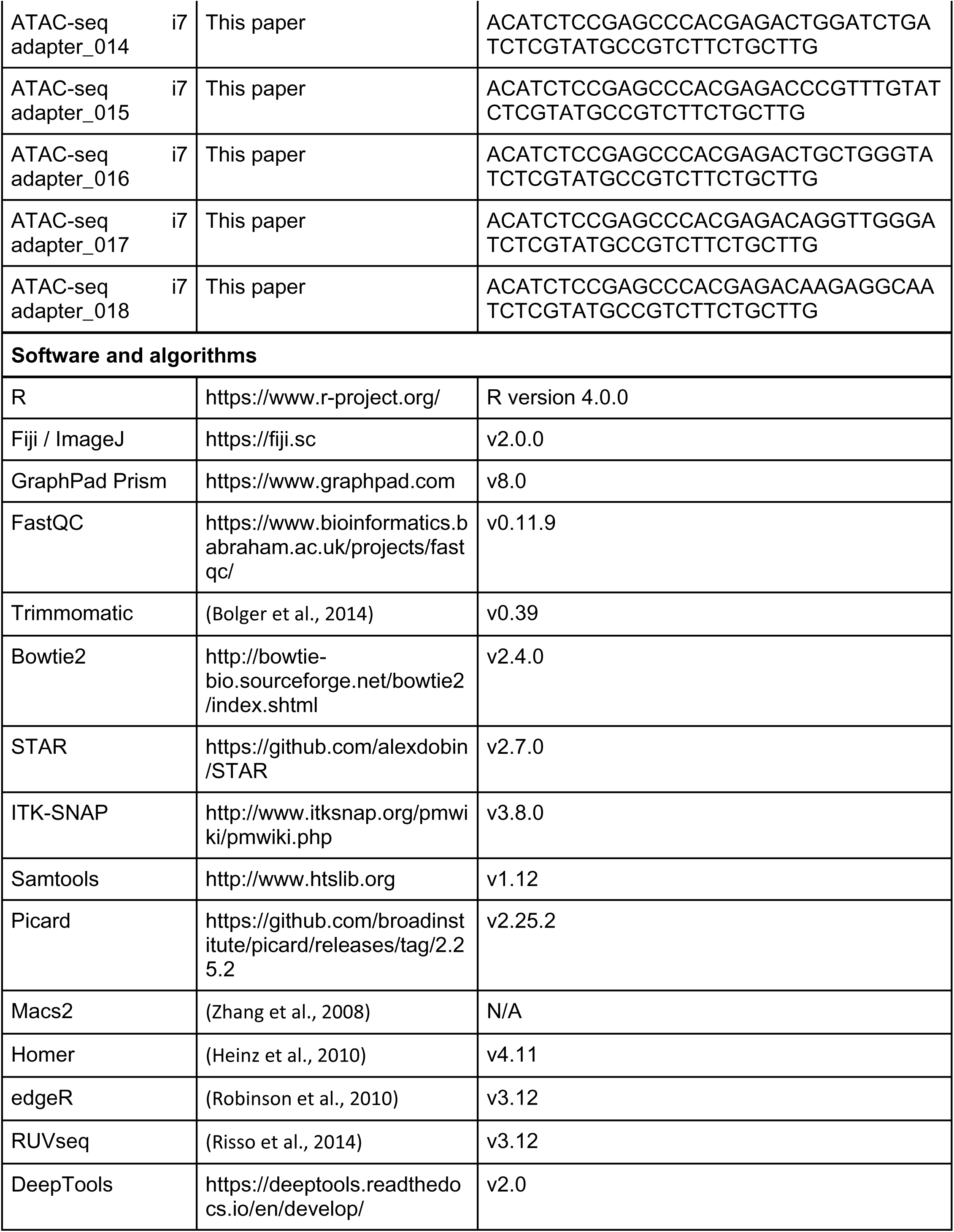
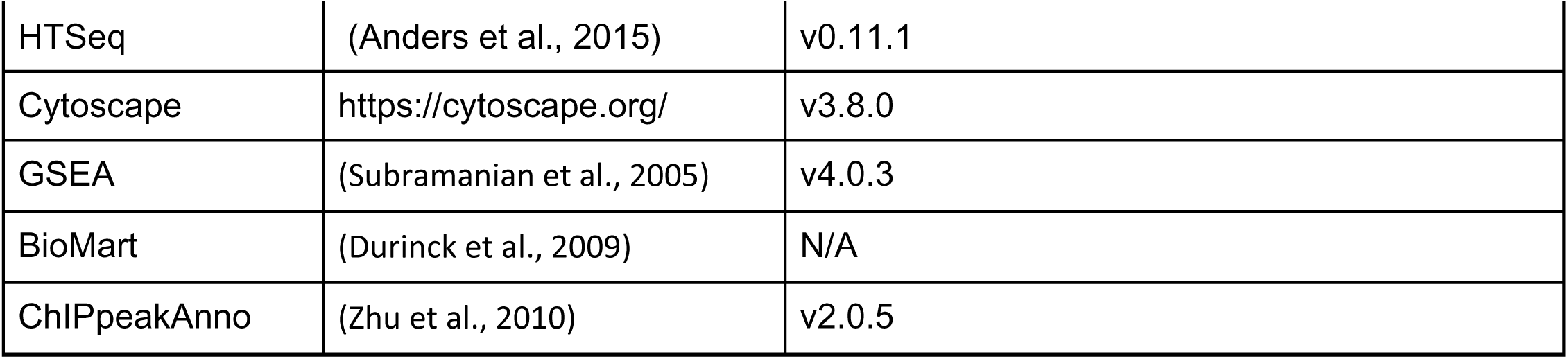

### Resource Availability

#### Lead Contact

Further information and requests for resources and reagents should be directed to and will be fulfilled by the lead contact, Dr. Joseph D. Dougherty.

#### Materials availability

Mouse line generated in this study is available upon request.

#### Data and code availability

The codes for analysing illumina sequencing, ATAC-seq, and RNA-seq generated in this study are available upon request.

The ATAC-seq and RNA-seq raws reads as well as counts data are available at…

### Experimental model and subject details

#### Human subjects

All procedures with human subjects were approved by the Washington University Institutional Review Board (201603131).

#### Animal models

All procedures using mice were approved by the Institutional Care and Use Committee at Washington University School of Medicine and conducted in accordance with the approved Animal Studies Protocol. All mice used in this study were bred and maintained in the vivarium at Washington University in St. Louis in individually ventilated (36.2 x 17.1 x 13 cm) or static (28.5 x 17.5 x 12 cm; post-weaning behavior only) translucent plastic cages with corncob bedding and *ad libitum* access to standard lab diet and water. Animals were kept at 12/12 hour light/dark cycle, and room temperature (20-22°C) and relative humidity (50%) were controlled automatically. For all experiments, adequate measures were taken to minimize any pain or discomfort. Breeding pairs for experimental cohorts comprised *Myt1l* Hets and wild type C57BL/6J mice (JAX Stock No. 000664) to generate male and female *Myt1l* Het and WT littermates. For embryonic ATAC-seq, RNA-seq, and EdU labeling, *Myt1l* Het x Het breeding pairs were used to generate *Myt1l* WT, Het and homozygous mutant littermates. Animals were weaned at P21, and group-housed by sex and genotype. Biological replicates for all experiments are sex and genitype balanced.

### Method details

#### Generation of MYT1L knockout mice

A Cas9 gRNA was designed to target the 7th exon of the mouse MYT1L gene (seq: 5’ GCTCTTGCTACACGTGCTACNGG 3’), similar to where a patient specific heterozygous *de novo* mutation had been defined by our clinical colleagues in human case with ASD (c.2117dupG). Cutting efficiency of reagents and homologous recombination was confirmed in cell culture. Then validated gRNA and Cas9 protein (IDT) were electroporated into fertilized C57BL6/J oocytes along with single stranded oligonucleotides carrying homology to the targeted region and the G mutation (Seq: 5’ accagcagctatgcacctagcagcagcagcaacctcagctgtggtggtggcagGcagcgccTCCagTacgtgtagcaagagcagcttt gacta cacacatgacatggaggccgcacacatggcagcc 3’) as well as blocking oligonucleotides (Seq: 5’ accagcagctatgcacctagcagcagcagcaacctcagctgtggtggtggcagcagcgccTCCagTacgtgtagcaagagcagctttgactacacacatgacatggaggccgcacacatggcagcc 3’) for the other strand to prevent homozygous mutation and presumptive embryonic lethality of founders. Eggs were cultured for 1-2 hours to confirm viability, then transferred to pseudopregnant surrogate dams for gestation. Pups were then screened for the targeted allele by amplicon PCR with mutation flanking primers followed by Illumina sequencing.

Founders carrying the appropriate allele were then bred with wild type C57BL/6J mice (JAX Stock No. 000664) to confirm transmission. F1 pups from the lead founder were genotyped by sequencing as above, then bred to generate experimental animals. Subsequent genotyping at each generation was conducted utilizing allele specific PCR using the MYT1L mutant primers and control primers, amplified using Phusion and the following cycling conditions: 98°C for 3 min, 98°C for 10 s, 61°C for 20 s, 72°C for 20 s, repeat 2-4 for 35 cycles, 72°C for 5 min, and hold at 4°C.

#### RNA extraction and RT-qPCR

Mice brains or cortex were dissected out at different developmental stages and homogenized in lysis buffer (10 mM Tris-HCl, pH 7.4, 10 mM NaCl, 3 mM MgCl2, 0.1% IGEPAL CA-630, 0.1% RNase inhibitor) on ice. Then lysates were mixed with Trizol LS and chloroform. After centrifugation, RNA was extracted from the aqueous layer with Zymo RNA Clean and ConcentratorTM-5 kit. cDNA libraries were prepared using qScript cDNA synthesis Kit (QuantaBio). RT-qPCR were performed using SYBR Green Master Mix (Thermo Fisher) on QuantStudio 6 Flex Real Time PCR System using primers in the **Key Resources Table**. We normalized cycle counts to GAPDH and calculated normalized relative gene expression using ΔΔCT. To compare MYT1L mRNA expression between genotypes, we put 6 WT and 8 Het brains into qPCR procedure. To understand MYT1L expression in human brain, we acquired normalized RNA-seq RPKM values of MYT1L in primary somatosensory cortex (S1C) from Allen Brain Atlas BrainSpan dataset (http://www.brainspan.org/) and plotted MYT1L mRNA temporal expression in R.

#### Western Blot

Mice brains or cortex were dissected out at different developmental stages and homogenized in lysis buffer (50 mM Tris-HCl, pH 7.4, 100 mM NaCl, 3 mM MgCl2, 1% IGEPAL CA-630, 10 mM NaF, 10 mM Na3Vo4 with Protease inhibitors). After centrifugation, supernatants were collected and protein concentration was measured by BCA assay. For each sample, 20 µg of protein was run on the 7.5% BioRad precast gel and transferred to the PVDF membrane. We blocked the membrane using TBST with 3% BSA for 2 hours at room temperature (RT). Then, the membrane was incubated with anti-MYT1L (1:500, 25234-1-AP, Proteintech) and anti-GAPDH (1:5000, G8795, Sigma) primary antibodies overnight at 4°C and then incubated with HRP conjugated anti-Mouse (1:2000, 1706516, BioRad) and anti-goat (1:2000, AP307P, Millipore) for one hour at RT. After washing, the membrane was developed in BioRad ECL Western Blotting Substrates and imaged with myECL Imager (Thermo Fisher). Fluorescent intensity was measured by ImageJ and MYT1L expression was normalized to GAPDH. To compare MYT1L protein expression between genotypes, we put 3 WT and 4 Het brains into Western Blot procedure.

#### Immunofluorescence

Mice brains were dissected out at different developmental stages and fixed in 4% paraformaldehyde (PFA) overnight at 4°C. After gradient sucrose dehydration and O.C.T. compound embedding, brains were sectioned using Leica Cryostat (15 µm for E14 brains and 30 µm for postnatal brains), Antigen retrieval was performed by boiling sections in 95°C 10 nM sodium citrate (pH 6.0, 0.05% Tween-20) for 10 mins. Then sections were incubated in the blocking buffer (5% normal donkey serum, 0.1% Triton X-100 in PBS) at RT for 1 hour. Primary antibodies, including anti-MYT1L (1:500, 25234-1-AP, Proteintech), anti-MAP2 (1:200, #188044, SYSY), anti-SOX2 (1:200, sc-17320, Santa Cruz), anti-TBR2 (1:400, AB15894, Millipore), anti-Ki-67 (1:500, #14-5698-82, Invitrogen), anti-CTIP2 (1:500, ab18465, Abcam), anti-BRN2 (1:500, sc-393324, Santa Cruz), anti-NEUN (1:500, #12943, Cell Signaling), anti-GFAP (1:500, ab53554, Abcam), anti-OLIG2 (1:200, AF2418, R&D Systems), and anti-TBR1 (1:500, ab31940, Abcam) were used to detect different cell markers. Next, sections were incubated in fluorescence conjugated secondary antibodies, including donkey anti-rabbit (Alexa 488, 546, and 647, Invitrogen), donkey anti-mouse (Alexa 546, Invitrogen), donkey anti-chicken(Alexa 488, Jackson ImmunoResearch), donkey anti-rat (Alexa 488 and 647, Invitrogen), and donkey anti-goat (Alexa 488 and 647, Jackson ImmunoResearch) at 1:500 dilution for 2 hours in RT. Images were captured under Zeiss Confocal Microscope or Zeiss Axio Scan Slide Scanner and cell counting was performed using ImageJ. In order to compare cell numbers of different cell types across genotypes, we had 5 WT, 6 Het, and 5 KO E14 brains for cell counting experiments (Fig. 4A). And we had 6 WT, 6 Het, and 5 KO E14 brains to quantify the Ki-67 positive cells (Fig. 4G,H).

#### Sanger Sequencing

Genomic DNA (gDNA) was extracted from mouse tissue by Qiagen Blood and Tissue Kit. a 2.2kb gDNA fragment flanking the G duplication site was amplified using the primers (**Key Resources Table**), Phusion, and following program: 98°C for 2 min, 98°C for 10 s, 60°C for 20 s, 72°C for 1 min, repeat 2-4 for 30 cycles, 72°C for 5 min, and hold at 4°C. PCR products were purified with QIAquick PCR Purification Kit and submitted for sanger sequencing at Genewiz. We used Snapgene to check and visualize sanger sequencing results.

#### Illumina Sequencing

gDNA and cDNA library from mice brains was generated as described in the above sections. To prepare sequencing libraries, we performed two-step PCR to first tag 200bp DNA fragments flaking the mutation site with illumina adapters (Taq, primers seen **Key Resources Table**, PCR program: 94°C for 3 min, 94°C for 10 s, 58°C for 20 s, 68°C for 1 min, repeat 2-4 for 30 cycles, 68°C for 5 min, and hold at 4°C) and then add unique index to individual samples (Taq, primers seen supplemental tables, PCR program: 98°C for 3 min, 98°C for 10 s, 64°C for 30 s, 72°C for 1 min, repeat 2-4 for 20 cycles, 72°C for 5 min, and hold at 4°C). Final PCR products were purified by gel extraction using Qiagen Gel Extraction Kit and submit for 2✕150 Illumina sequencing to CGSSB at Washington University School of Medicine. For each sample, we were able to get ∼80,000 reads. We conducted quality control on raw reads using Fastqc. Then reads were trimmed by Trimmomatic software and aligned to the mouse genome by STAR. We used VarScan and Samtools to determine the percentage of the mutation in gDNA(n = 8) and cDNA(n = 8) samples.

#### Nissl Staining

Following perfusion with 4% paraformaldehyde, the brains were removed, weighed (WT n = 5, Het n = 6), sectioned coronally using a vibratome at 70 μm, and then mounted onto gelatin coated slides (WT n = 8, Het n = 9). Sections were then rehydrated for 5 minutes in xylene, xylene, 100% ethanol, 100% ethanol, 95% ethanol, 70% ethanol, and deionized water. Using 0.1% cresyl violet at 60°C, sections were stained for two hours and rinsed with two exchanges of deionized water. Differentiation began with 30 second rinses in 70% ethanol, 80% ethanol, and 90% ethanol. Next, a two-minute rinse in 95% ethanol was done, checking microscopically for a clearing background. This was followed by a 30-second rinse in two exchanges of 100% ethanol, a 15-minute rinse using 50% xylene in ethanol, and a 1-hour rinse of xylene. Finally, the sections were mounted and coverslipped using DPX mountant. Whole and regional volumes were outlined by a rater blind to treatment using Stereoinvestigator Software (v 2019.1.3, MBF Bioscience, Williston, Vermont, USA) running on a Dell Precision Tower 5810 computer connected to a QImaging 2000R camera and a Labophot-2 Nikon microscope with electronically driven motorized stage.

#### In vivo Magnetic Resonance Imaging (MRI): data acquisition

All animal experiments were approved by Washington University’s Institutional Animal Care and Use Committee. MRI experiments were performed on a small-animal MR scanner built around an Oxford Instruments (Oxford, United Kingdom) 4.7T horizontal-bore superconducting magnet and equipped with an Agilent/Varian (Santa Clara, CA) DirectDrive^TM^ console. Data were collected with a laboratory-built, actively-decoupled 7.5-cm ID volume coil (transmit)/1.5-cm OD surface coil (receive) RF coil pair. Mice were anesthetized with isoflurane/O_2_ (1.2% v/v) and body temperature was maintained at 37±1°C *via* circulating warm water. Mouse respiratory rate (50-70 breaths/minutes) and body temperature (rectal probe) were monitored with a Small Animal Instruments (SAI, Stony Brook, NY) monitoring and gating unit.

T2-weighted transaxial images (T2W) were collected with a 2D fast spin-echo multi-slice (FSEMS) sequence: echo train length=4, kzero=4, repetition time (TR)=1.5 s, effective echo time (TE)=60 ms; field of view (FOV)=24 x 24 mm^2^, matrix size =192 x 192, slice thickness=0.5 mm, 21 slices, 4 averages. Co-registered T1-weighted images (T1W) were collected with a 2D spin-echo multi-slice (SEMS) sequence: TR=0.8 s, TE=11.3 ms, 2 averages.

Diffusion Tensor Imaging (DTI) measures the directional water movement along and perpendicular to axons (fractional anisotropy: FA) as a measure of white-matter integrity, and the same images can be used for structural assessments. DTI data were collected using a multi-echo, spin-echo diffusion-weighted sequence with 25-direction diffusion encoding, max b-value=2200 s/mm^2^. Two echoes were collected per scan, with an echo spacing of 23.4 ms, and combined offline to increase signal-to-noise ratio (SNR), resulting in a SNR improvement of 1.4x compared with a single echo. Other MR acquisition parameters were TR=1.5 s, TE=32 ms, length of diffusion-encoding gradients (δ)=6 ms, spacing between diffusion gradients (Δ)=18 ms, FOV = 24 mm x 24 mm, matrix size = 192 x 192, slice thickness=0.5 mm, 21 slices, 1 average. The total acquisition time was approximately 2 hours and 5 minutes.

#### DTI Data Analysis

DTI datasets were analyzed in MatLab (The MathWorks®, Natick MA). Following zero-padding of the k-space data to matrix size 384 x 384, the data were Fourier-transformed and the images from the two spin echoes were added together. A 3 x 3 Gaussian filter (Sigma = 0.7) was applied and the resulting images were fit as a mono-exponential decay using the standard MR diffusion equation (Stejskal and Tanner, 1965):

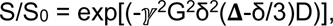

in which S is the diffusion-weighted signal intensity, S_0_ the signal intensity without diffusion weighting, γ is the gyromagnetic ratio, G is the gradient strength, and D is the diffusion coefficient. Eigenvalues (*λ*_1_,*λ*_2_,*λ*_3_) corresponding to the diffusion coefficients in three orthogonal directions were calculated and parametric maps of apparent diffusion coefficient (ADC), axial diffusion (D_axial_), radial diffusion (D_radial_), and fractional anisotropy (FA) were calculated according to standard methods (Basser and Pierpaoli, 2011; Mori, 2007). Parametric maps were converted into NIfTI (.nii) files for inspection and segmentation in ITK-SNAP (www.itksnap.org). We ended up analysing 8 WT mice and 6 Het mice.

#### ATAC-seq

ATAC-seq was performed as described before (Buenrostro et al., 2015). Briefly, mouse E14 cortex or adult PFC (P60-P70) was dissected and gently homogenized in cold nuclear isolation buffer (10 mM Tris-HCl, pH 7.4, 10 mM NaCl, 3 mM MgCl2, 0.1% IGEPAL CA-630). Embryonic tissues were pooled across sexes, adult tissues included both sexes, balanced for genotype. Lysates were filtered through 40 µm mesh strainer. After spinning down, 100,000 nuclei were put into the tagmentation reaction for each sample. We had 6 WT, 5 Het and 6 KO cortex for embryonic experiments. For adult PFC experiments, we put 6 WT and 6 PFC into the pipeline. Tagmentation reaction was performed using Illumina Tagment DNA TDE1 Enzyme and Buffer Kit with 30 min incubation time at 37°C. Immediately following the tagmentation, we purified DNA fragments using QIAquick PCR Purification Kit. We took half amount of purified DNA fragments and added Illumina Nextera i5+i7 adapters with unique index to individual samples by PCR reaction (Phusion, primers seen in **Key Resources Table**, PCR program: 72°C for 5 min, 98°C for 30 s, 98°C for 10 s, 63°C for 30 s, 72°C for 1 min, repeat 3-5 for 8-10 cycles, and hold at 10°C). Generated libraries were purified using AMpure beads (1:1.8 dilution). We ran Tapestation for libraries and checked the nucleosome peaks pattern as quality control. Finally, libraries were submitted to GTAC Washington University School of Medicine for Novaseq aiming for 50M reads per sample.

#### DAR analysis

Raw reads were trimmed by Trimmomatic software to remove adapter sequence. We used Fastqc to check reads quality before and after trimming. Then reads were mapped to mm10 genome by Bowtie2. We filtered out mitochondrial reads (Samtools), PCR duplicates (Picard), non-unique alignments (MAPQ > 30), and unmapped reads (Samtools). Then a series of QC metrics were examined to ensure ATAC experiments worked well, including insert size distribution, mitochondria reads percentage, non-redundant reads percentage, and TSS enrichment. To adjust reads start sites, we shifted reads aligned to + strand by +4bp and reads aligned to - strand by −5 bp by bedtools and awk. After shifting, we merged bam files for all samples in one specific time stage (E14 or adult) together and performed peak calling by MACS2 with q < 0.05. Peaks were annotated by Homer software. In order to perform differential accessible region analysis, we derived peaks read counts from individual sample’s shifted bam file using bedtools. With read counts, utilized edgeR package to identify DARs. Briefly, we first checked library size, read counts distribution, pearson correlation, and multidimensional scale plots and identified no obvious outlier sample. Then we normalized reads and removed unwanted variables using the RUVseq package. For E14 cortex ATAC-seq, we fitted the data into a nested interaction model to identify altered chromatin accessibility across all genotypes (WT, Het, and KO). And we considered peaks with the same significant fold change (FDR < .1) direction in Het and KO as true DARs. For adult PFC, a negative binomial generalized linear model was fitted and sex was counted as covariate when testing for DARs (FDR < .1). Heatmaps for DARs were generated by deepTools. TSS peaks were defined as ±1kb from TSS and all other peaks were considered non-TSS peaks. MYT1L ChIP targets from Mall et al.’s Table S2 were mapped to ATAC-seq data sets by bedTools and we defined overlapping peaks between the two with 1kb maximum gap. Motif analysis was performed using Homer software on DARs (FDR < .1). We used more-accessible regions as background when finding motifs for less-accessible regions and vice versa.

#### RNA-seq

Embryonic cortex and adult PFC (P60-P70) was dissected out and RNA was extracted as described above. Embryonic tissues were pooled across sexes, adult Adult tissues included both sexes, balanced for genotypeTotal RNA integrity was determined using Agilent 4200 Tapestation. Library preparation was performed with 10ng of total RNA with a RIN score greater than 8.0. ds-cDNA was prepared using the SMARTer Ultra Low RNA kit for Illumina Sequencing (Takara-Clontech) per manufacturer’s protocol. cDNA was fragmented using a Covaris E220 sonicator using peak incident power 18, duty factor 20%, cycles per burst 50 for 120 seconds. cDNA was blunt ended, had an A base added to the 3’ ends, and then had Illumina sequencing adapters ligated to the ends. Ligated fragments were then amplified for 15 cycles using primers incorporating unique dual index tags. Fragments were sequenced on an Illumina NovaSeq-6000 using paired end reads extending 150 bases. Again, raw reads were trimmed by Trimmomatic software to remove adapter sequence and we used Fastqc to check reads quality before and after trimming. rRNA reads were filtered out by Bowtie2. And filtered reads were mapped to the mouse mm10 genome by STAR. Read counts for genes were derived by HTSeq software for individual samples. We checked read counts distribution, junction saturation, library size, pearson correlation and multidimensional scale plots to rule out any outliers. In the end, we were able to put 6 WT, 6 Het, 4 KO E14 cortex and 6 WT, 6 Het adult PFC into the DGE analysis pipeline.

#### Differential Gene Expression analysis

Similar to DAR analysis, we normalized raw counts and removed unwanted variables with the edgeR and RUVseq package. A nested interaction model was fitted to identify differential gene expression across genotypes for E14 cortex RNA-seq. DEGs with the same significant fold change direction in both Het and KO samples were considered as true MYT1L regulated genes and were subjected to downstream analysis. For adult PFC RNA-seq, we fitted the data to a negative binomial generalized linear model with sex as covariates. We applied cut-off FDR <.1 to define DEGs. Heatmaps for DEGs were generated by heatmap.2 function in R.

#### GO analysis

To perform GO analysis on DARs, we assigned DARs (FDR < .1) located within ±1kb from TSS to corresponding genes. GO analysis was performed using BiNGO in Cytoscape. *p* values were adjusted by Benjamini-Hochberg FDR correction and FDR < 0.05 cut-off was used to determine significant enrichments. The same software and corrected p value cut-off was applied to GO analysis on DEGs (FDR < .1) in RNA-seq. Full GO analysis results can be seen in **Table S4**.

#### GSEA analysis

GSEA was performed as described before (Subramanian et al., 2005) using GSEA v4.0.3 (https://www.gsea-msigdb.org/gsea/index.jsp). We first examined gene set collections H (Hallmark gene sets) and C2 (curated gene sets of online pathway databases) to understand how MYT1L loss affects different cellular processes in a comprehensive manner. Then we tested the expression changes of MYT1L ChIP targets, human “early-fetal” and “mid-fetal” genes (Kang et al., 2011; Katayama et al., 2016), MEF signature genes, induced neuron signature genes on E14 cortex and adult PFC expression data (See **Table S3**), Wnt signaling genes (MGI GO:0016055), and Notch signaling genes (MGI GO:0007219). Human gene IDs were converted into mouse gene IDs by BioMart (https://www.ensembl.org/biomart). All analysis was performed with “gene_set” as permutation type and 1,000 permutations. Significant enrichment was determined by FDR <. 1 cut-off.

#### Comparison between in vivo and in vitro RNA-seq

In vitro RNA-seq data were obtained from Mall et al., 2017 studies on MYT1L overexpression (OE) in MEF and shRNA knockdown (KD) in primary hippocampal neuron cultures (Mall et al., 2017). We defined genes that showed upregulation in OE but downregulation in KD as MYT1L induced genes, while genes getting downregulated in OE but upregulated in KD were considered as MYT1L repressed genes. Then, the hypergeometric test was performed to determine whether there is significant overlapping between DEGs from our *in vivo* RNA-seq experiments and previously reported MYT1L targeted genes *in vitro*. We also used R to investigate linear regression of DEGs’ fold changes between *in vivo* and *in vitro* RNA-seq experiments.

#### Disease models and human genetic data sets enrichment

DEGs of different ID/ASD related mouse model were derived from CHD8 haploinsufficient cortex (*p* < .05 for E14.5, FDR < .1 for P77)(Gompers et al., 2017), KDM5C KO frontal cortex (*p* < 0.01)(Iwase et al., 2016), CHD2 haploinsufficient embryonic cortex (*p* < .05) and P30 hippocampus (FDR < .1)(Kim et al., 2018), PHF6 KO cortex (FDR < 0.05)(Cheng et al., 2018), FOXP1 KO hippocampus (FDR < 0.05)(Araujo et al., 2015), and POGZ cKO hippocampus (FDR < .05)(Suliman-Lavie et al., 2020). For human diseases genetic data sets, we downloaded ASD genes from SFARI (huamn module, gene score 1 and 2), ADHD genes from ADHDgene (http://adhd.psych.ac.cn/), ID genes from IDGenetics (http://www.ccgenomics.cn/IDGenetics/), SCZ genes from SZDB2.0 SNP data sets (http://www.szdb.org/), and Microcephaly genes from DisGeNET (https://www.disgenet.org/home/). Enrichment analysis was performed using the one-sided hypergeometric test and p values were adjusted by Benjamini-Hochberg correction.

#### EdU labeling

We performed intraperitoneal injection on E14 timed-pregnant females with EdU solution (50mg/kg). For the cell proliferation assay, we waited for 1.5 hours before harvesting embryonic brains. Brains were dissected and fixed with 4% PFA at 4°C overnight. Then we dehydrated and sectioned brains into 15 µm sections on glass slides as described in the immunofluorescence session. Those sections were subjected to EdU detection assay using Click-iT EdU Cell Proliferation Kit for Imaging Alexa Fluor-594 (Invitrogen) with manufacturer instructions. 4 animals per genotype were used for cell proliferation assay.

For the cell cycle existing assay, we waited for 20 hours before harvesting brains. The same procedure was conducted on fixed brains to get 15 µm sections. Then, antigen retrieval was performed by boiling sections in 95°C 10 nM sodium citrate (pH 6.0, 0.05% Tween-20) for 10 mins. Brain sections were first incubated with anti-Ki-67 primary antibody and Alexa488-fluorescence conjugated secondary antibody before EdU detection assay. EdU+/Ki67+ cells represent neuronal progenitors that still remained in the cell cycle, while EdU+/Ki67-cells represent differentiating progenitors that exited the cell cycle. We calculated Q fraction value as the ratio between EdU+/Ki67-cells and total EdU+ cells to assess the portion of cells starting differentiation within the 20-hour time window. All images were captured under Zeiss Confocal Microscope and cell counting was performed using ImageJ. 4 animals per genotype were used for cell cycle existing assay.

#### Slice Preparation

Coronal brain slices (325 μm) containing V1 were obtained as previously described (Lambo and Turrigiano, 2013) using chilled (1°C) standard artificial CSF (ACSF). ACSF was continuously oxygenated and contained the following (in mm): 126 NaCl, 3 KCl, 2 MgSO4, 1 NaHPO4, 25 NaHCO3, 2 CaCl2, and 14 Dextrose. Slices were cut on a Leica VT1000S vibratome and incubated on a semipermeable membrane covered by room temperature oxygenated standard ACSF.

#### Slice Electrophysiology

V1m was identified, and whole-cell patch-clamp recordings obtained from layer 2/3 pyramidal neurons, as previously described (Lambo and Turrigiano, 2013). In brief, V1m was identified using the mouse brain atlas after adjusting for the lambda-bregma distance for age. The shape and morphology of the white matter were used to identify V1m. Neurons were visualized with a 40× water-immersion objective using infrared-differential interference contrast optics. Internal recording solution contained (mm): 20 KCl, 100 K-gluconate, 10 HEPES, 4 Mg-ATP, 0.3 Na-GTP, 10 phosphocreatine, and 0.4% biocytin. For AMPA miniature EPSC (mEPSC) recordings, neurons were voltage-clamped to −70 mV in standard ACSF containing TTX (0.2 μm), APV (50 μm), and picrotoxin (20 μm) and warmed to 33°C. For AMPA miniature IPSC (mIPSC) recordings, internal recording solution contained (mM): 120 KCl, 10 HEPES, 4 Mg-ATP, 0.3 Na-GTP, 2.5 phosphocreatine, and 0.2% biocytin. Neurons were voltage-clamped to −70 mV in standard ACSF containing TTX (0.2 μm), APV (50 μm), and DNQX (20 μm). For all recordings, Neurons were excluded from analyses if the resting membrane potential was more positive than −50 mV, input resistance was <40 MΩ, series resistance was >20 MΩ, or if any of these parameters changed by >20% during the recording. Pyramidal neurons were identified by the presence of an apical dendrite and tear-drop shaped soma and morphology was confirmed by post hoc reconstruction of biocytin fills, as described previously (Desai et al., 2002). All physiology data were analyzed using Clampfit (Molecular Devices) and custom software written in Python (available at github.com/hengenlab). We recorded 24 neurons from 9 WT animals and 22 neurons from 9 Het animals to compare the passive properties as well as mEPSC (100 events for each recorded neuron) activities across genotypes. We also recorded the mIPSC of 17 neurons from 5 WT animals and 22 neurons from 5 Het animals to assess the E/I balance.

#### Neuronal Morphology Analysis

Brain slices from slice electrophysiology were subjected to histochemical analysis using NEUN antibody to confirm neuron identity and streptavidin Alex Fluor-568 (Invitrogen) to label injected biocytin for morphology assessment. Stained sections were mounted in cell gasket with SlowFade Diamond Antifade Mountant (Invitrogen). Images for neuronal body and dendrites were taken under Zeiss LSM 880 Airyscan Confocal Microscope. We used Neurolucida 360 (https://www.mbfbioscience.com/neurolucida360) to trace the neuronal body (15 neurons from 8 WT animals, 14 neurons from 8 Het animals) and dendrites (10 neurons from 5 WT animals, 10 neurons from 6 Het animals) and count different types of dendritic spines (10 neurons from 4 WT animals, 7 neurons from 4 Het animals). Branch analysis and sholl analysis were performed using Neurolucida Explorer (https://www.mbfbioscience.com/neurolucida-explorer). Then we exported measurements for soma surface area, soma volume, total dendrite number, total dendritic length, average dendrite length, dendrite node number, and complexity ([Sum of the terminal orders + Number of terminals] * [Total dendritic length / Number of primary dendrites]), branch number, branch length, total spine density, and density of different spine subtypes to compare neuron morphological maturation between Hets and WTs.

#### Behavioral Analysis

##### Animals and experimental design

The behavior phenotypes we investigated were chosen based on the symptom profile of the index patient and that of the greater MYT1L deletion population. We examined the phenotypes of two independent cohorts. The first cohort comprised 57 Het (26 female and 31 male) and 55 WT (29 female and 26 male) mice, and was used to assess the first three weeks of postnatal development for gross, motor and communicative delays (**Table S6**). The second cohort comprised 20 Het (10 female and 10 male) and 21 WT (13 female and 8 male) mice. One female Het died after social operant testing, two male Hets died one month after conditioned fear testing, and another male Het died before T-maze testing. A third cohort comprising 16 WT (8 female and 8 male) and 14 Het (8 female and 6 male) mice was assayed for cognitive inflexibility in the T-maze. These mice were characterized beginning as juveniles and continued through adulthood, and assessed for behavioral features related to the neuropsychiatric diagnoses of our index patient (**Table S6**). A fourth cohort comprising 23 WT (9 female and 14 male) and 19 Het (11 female and 8 male) mice was assessed for tactile sensitivity using the von Frey filaments. ASD-related repetitive and social behaviors were investigated in the force-plate actometer, spontaneous alternation T-maze, the von Frey assessment of tactile sensitivity, social operant task, social dominance task, and three-chambered social approach assay. ADHD-related hyperactivity was assessed specifically using the open field task, but we also examined general activity across any task in which we conducted subject tracking. We looked at behaviors relevant to ID in the Barnes maze and fear conditioning tasks. To assess mature sensory and motor function, we used a battery of sensory motor tasks and the prepulse inhibition/startle apparatus. Finally, we documented weight throughout the lifespan, and performed assessments of physical features and posture to identify any dysmorphia. A male experimenter conducted the ultrasonic vocalization recordings, and a female experimenter conducted the remainder of the behavioral testing. Each experimenter was blinded to experimental group designations during testing, which occurred during the light phase. Order of tests was chosen to minimize effects of stress. Animals were acclimated to the testing rooms 30 - 60 min prior to testing.

##### Developmental assessment

During the first three weeks postnatal, we assessed the *Myt1l* Het and WT littermates for signs of gross developmental delay, communicative delay or motor delay, which are universal in MYT1L deletion patients (Blanchet et al., 2017) (See **Table S6**). To evaluate gross development, the mice were weighed daily from P5 - P21, and evaluated for physical milestones of development including pinna detachment by P5 and eye opening by P14. While human language cannot be explored in mice, vocal communication behavior is conserved across taxa (Ehret, 1980). Mouse pups produce isolation calls as a way to attract the dam for maternal care (Haack et al., 1983), thus it is one of the earliest forms of social communication we can examine in mice. This behavior also has a developmental trajectory, beginning just after birth, peaking during first week postnatal and disappearing around P14, making it useful for examining delay in early social circuits. Ultrasonic vocalizations (USVs) were recorded on P5, P7. P9 and P11 following our previous methods (Maloney et al., 2018a). Briefly, the dam was removed from the nest and the litter placed in a warming cabinet. The surface temperature of each pup was recorded (HDE Infrared Thermometer; Het: *M*=35.4°C, *SD*=0.90; WT: *M*=35.2°C, *SD*=1.16), and then the pup was placed in an empty cage (28.5 x 17.5 x 12 cm) in a sound-attenuating chamber. USVs were recorded for three minutes using an Avisoft UltraSoundGate CM16 microphone, Avisoft UltraSoundGate 116H amplifier, and Avisoft Recorder software (gain = 3 dB, 16 bits, sampling rate = 250 kHz). The pup was then removed, weighed, tissue collected for genotyping (P5 only), and returned to the nest. Following recording of the last pup, the dam was returned to the nest. Frequency sonograms were prepared from recordings in MATLAB (frequency range = 25 kHz to 120 kHz, FFT size = 512, overlap = 50%, time resolution = 1.024 ms, frequency resolution = 488.2 Hz). Individual syllables and other spectral features were identified and counted from the sonograms as previously described (Holy and Guo, 2005; Rieger and Dougherty, 2016).

Possible motor delay was assessed with a battery of tasks conducted during the first two weeks postnatal (Feather-Schussler and Ferguson, 2016), which assess the acquisition of motor function, including ambulation, posture, reflexes, and muscle strength and endurance (See **Table S6**). A few of key reflexes appear in mouse pups in the first week, including the righting, grasping and negative geotaxis reflexes at about P5-P7. To assess surface righting reflex (P6 and P14), each pup was placed on its back in an empty cage lined with a plastic bench pad and the time to return to a prone position was recorded up to 60 sec (Fig. 7K). Three trials were averaged for analysis. Acquisition of grasping reflex was assessed (P6 and P14) by placing the blunt side of a razor blade against the palmar surface of each paw and recording the presence or absence of grasping (Fig. 7J). Negative geotaxis was evaluated (P10) by placing the pup facing downward on a 45° incline (Fig. 7L). The time up to two min the pup required to turn 180° was recorded. Three trials were averaged for analysis. Mice start to ambulate at P5 by crawling and are fully walking by P10. So we examined their ambulation at P8 to identify any delays (Fig. 7I). We also looked at the posture of their hindlimb – with maturation of ambulation, the hindlimb angle narrows. Each pup was placed in an empty cage (36.2 x 17.1 x 13 cm) and their ambulation was scored over a 3 min period using the following scale: 0 = no movement, 1 = crawling with asymmetric limb movement, 2 = slow crawling but symmetric limb movement, and 3 = fast crawling/walking. Video of ambulation was recorded at the same time and angle of the hindlimbs was measured with lines from mid-heel through middle digit across three separate frames, which were averaged for analysis. Muscle strength and endurance were assessed with forelimb and hindlimb suspension tests (P10 and P12, respectively) and grip strength (P10, 12 and 14). For forelimb suspension, each pup was allowed to grasp a wire strung across a pencil cup with felt padding with both forepaws (Fig. 7M). Latency to release from the wire was recorded across three separate trials that were averaged for analysis. One influential outlier (*z*=3.63) was excluded from analysis. Hindlimb suspension ability was measured by placing the pup facing downward into a 50 mL conical tube with the hindlimbs hung over the edge (Fig. 7M). Latency to release from the conical edge was recorded across three separate trials that were averaged for analysis. Grip strength was measured by placing each pup in the middle of a horizontal fiberglass wire screen, and slowly rotating the screen vertically until inverted 180° (Fig. 7N). The angle at which the pup fell from the screen onto a bench pad was recorded across three separate trials, which were averaged for analysis.

##### Open field

Locomotor ambulation was measured at P30 to assess activity, exploration, and anxiety-like levels in the open field assay similar to our previous work (Maloney et al., 2018b). Specifically, the behavior of each mouse was evaluated over a 1-hr period in a translucent acrylic apparatus measuring 59 x 39 x 22 cm (Fig. S8A), housed inside a custom sound-attenuating chamber (70.5 x 50.5 x 60 cm), under approximately 9 lux illumination (LED Color-Changing Flex Ribbon Lights, Commercial Electric, Home Depot, Atlanta, GA). A CCTV camera (SuperCircuits, Austin, TX) connected to a PC computer running the software program ANY-maze (Stoelting Co., Wood Dale, IL; http://www.anymaze.co.uk/) tracked the movement of the mouse within the apparatus to quantify distance traveled, and time spent in and entries into the center 50% and outer perimeter zones. The apparatus was cleaned between animals with a 0.02% chlorhexidine diacetate solution (Nolvasan, Zoetis, Parsippany-Troy Hills, NJ).

Pose estimation (DeepLabCut (Mathis et al., 2018) and machine learning classification (SimBA (Nilsson et al., 2020)) were used to further quantify behaviors of the mice in videos recorded during the open field test. Specifically, we used DeepLabCut to estimate the pose of eight body parts of the mice, including nose, left ear, right ear, center, lateral left, lateral right, tail base, and tail end. A random subset of frames from all 41 videos were used for the network training. The trained network was then applied to all videos, yielding pose tracking files. The video and the tracking file of a Het female mouse were input to SimBA to build classifiers for jumping (Fig. S8F), facial grooming, and body/tail grooming. A region of interest (ROI) defined as a rectangle covering the center area of the open field was appended to the machine learning features extracted from the tracking file. Then the training video was annotated for interesting behaviors using the SimBA event-logger. Random forest classifiers were trained using default hyperparameters, and classifier performances were evaluated. We set the discrimination threshold of jumping, facial grooming and body/tail grooming to 0.8, 0.444, and 0.521 respectively. The minimum behavior bout length (ms) for all behaviors was set to 200. In the end, the classifiers were applied to analyze all the videos. Facial grooming and body/tail grooming were combined for analyses. The descriptive statistics for each predictive classifier in the project, including the total time, the number of frames, total number of ‘bouts’, mean and median bout interval, time to first occurrence, and mean and median interval between each bout, were generated.

##### Force-plate actometer

At P36, the presence of stereotyped movements indicative of self-grooming and presence of tremor resulting from possible demyelination was assessed in the force-plate actometer (FPA; Fig. S8H), as previously described (Reddy et al., 2012; Tischfield et al., 2017). The custom made FPA consisted of a carbon fiber/nomex composite material load plate measuring 24 × 24 cm surrounded by a clear polycarbonate cage (15 cm high) with a removable clear polycarbonate top perforated with ventilation holes, and housed in a sound-attenuating cabinet measuring 70.5 x 50.5 x 60 cm. Force was measured by summing the signal from four transducers, which is then expressed as a percentage of body weight. Grooming only takes place during low mobility bouts, as previously defined (Reddy et al., 2012) and validated (Tischfield et al., 2017). Raw data was acquired with a DOS-based Free Pascal program and further processed using custom MATLAB scripts (Fowler et al., 2001). To identify any tremor, each force time series was Fourier transformed to identify unique frequencies and plotted as a continuous function or power spectra. Tremor was identified as the frequency at peak power.

##### Barnes maze

Spatial learning and memory was evaluated in the Barnes maze using methods adapted from previous work (Pitts, 2018). The Barnes maze apparatus consisted of a circular white acrylic platform measuring 122 cm in diameter, with 20 equally spaced holes (5 cm in diameter) around the perimeter 6.35 cm from the edge, elevated 80 cm from the floor (Fig. S7A,B). The maze was brightly lit with overhead lighting, and extra maze cues were used to aid learning. Testing comprised two acquisition trials separated by 45 minutes on each of 5 consecutive days. During acquisition trials, an escape box measuring 15.2 x 12.7 cm with an inclined entry was attached to the maze underneath one hole location (three escape locations were counterbalanced across mice). Prior to the first trial on the first day, each mouse was placed in the escape hole for 30 seconds covered by a clear acrylic tube. During each trial, a mouse was placed in the center of the maze facing a random direction, 75 dB white noise sounded until the mouse entered the escape box, which ended the trial. Each mouse was allowed to remain in the escape box for 30 seconds. If the mouse failed to enter the escape box, the trial would end after a maximum of three minutes and the mouse would be placed in the escape box for 30 seconds. On the sixth day, a three minute probe trial was conducted to assess each animal’s memory for the previously learned location of the escape box. The escape box was removed, and a mouse was placed in the center of the maze facing a random direction, and 75 dB white noise sounded until the end of the trial. A digital USB 2.0 CMOS Camera (Stoelting Co., Wood Dale, IL) connected to a PC computer running the software program ANY-maze (Stoelting Co., Wood Dale, IL; http://www.anymaze.co.uk/) tracked the movement of the mouse within the apparatus to quantify distance traveled, frequency and duration of visits to the escape box and to incorrect holes. All males were tested first, followed by the females. The apparatus was cleaned between animals with a 0.02% chlorhexidine diacetate solution (Nolvasan, Zoetis, Parsippany-Troy Hills, NJ).

##### Social operant

Social motivation, including social reward seeking and social orienting (Chevallier et al., 2012), was evaluated from P48-P60 using a social operant task adapted and extended from previous methods (Martin and Iceberg, 2015; Martin et al., 2014), adding continuous tracking to measure social reward seeking and social orienting in parallel. Standard operant chambers (Med Associates) enclosed in sound-attenuating chambers (Med Associates) were modified. A clear acrylic conspecific stimulus chamber (10.2 x 10.2 x 18.4 cm; Amac box, The Container Store) was attached to the side, separated from the operant chamber proper by a door opening (10.2 x 6 cm) with stainless steel bars (spaced 6mm apart), centered between the nosepoke holes (Fig. 8G). A 3D printed filament door was attached via fishing wire to a motor (Longruner) controlled by an Arduino (UNO R3 Board ATmega328P) connected to the Med Associates input panel. The chamber included a red cue light that illuminated at the beginning of the test trial and remained illuminated until the test trial ended. The rest of the chamber was illuminated with a puck light (Honwell) to achieve 54 lux. The operant chamber bottom tray was filled with one cup of fresh corn cob bedding, which was replaced between mice. Operant chambers and stimulus chambers were designated for males or females throughout the experiment. The operant chambers were cleaned with 70% ethanol and the stimulus chambers were cleaned with 0.02% chlorhexidine diacetate solution (acrylic; Nolvasan, Zoetis) between animals. One of the operant chamber holes was designated the “correct” hole, and the other the “incorrect” hole, which were counterbalanced across groups. A nosepoke into the correct hole triggered illumination of a cue light within that hole and the raising (opening) of the door between the operant and stimulus chambers. A nosepoke into the incorrect hole did not trigger an event. The experimental and stimulus animals were allowed to interact across the bars for 12 sec (social reward) and then the door was lowered (shut) and the correct hole cue light turned off. The operant chambers were connected to a PC computer via a power box (Med Associates). MED PC-V software quantified nosepokes as “correct”, “incorrect”, and “rewards” to measure social reward seeking behavior as part of social motivation. CCTV cameras (Vanxse) were mounted above the chambers and connected to a PC computer via BNC cables and quad processors. Any-Maze tracking software (Stoelting Co., Wood Dale, IL; http://www.anymaze.co.uk/) was used to track the experimental and stimulus animals’ behavior to quantify distance traveled, and time spent in and entries into the social interaction zone (6 x 3 cm zone in front of the door in each the operant and stimulus chamber). This allowed us to quantify the social orienting aspect of social motivation, defined as the experimental animal entering and spending time in the social interaction zone. Custom Java tools and SPSS syntax were used to align the Any-Maze tracking data with the timing of rewards in the Med Associates text data to extract presence or absence of each animal within the interaction zones during each second of every reward.

The operant paradigm comprised habituation, training, and testing trials (Fig. 8H). For all trials, sex- and age-matched, novel C57BL/6J mice served as conspecific stimulus mice. All mice, experimental and stimulus, were group housed by sex during the entirity of the operant paradigm. The stimulus mice were loaded into and removed from the stimulus chambers prior to the placement and after removal of the experimental mice into the operant chambers, respectively, to prevent the experimental animals from being in the chambers without a conspecific stimulus partner. Habituation consisted of a 30 minute trial on each of two consecutive days, during which the door remained opened, and the nosepoke holes were shifted to be inaccessible to prevent any nose-poking prior to training. This allowed the experimental mice to acclimate to the chamber and the presence of a stimulus partner in the adjoining chamber. Training consisted of 1-hr trials during which the fixed ratio 1 (FR1) reinforcement schedule was used to reward the mouse with a 12-sec social interaction opportunity following one correct nosepoke. During the 12-sec reward period, any additional correct nosepokes did not result in another reward. Each mouse received at least three days of FR1, after which achievement of learning criteria moved the mouse on to testing. Ten days of FR1 without reaching criteria resulted in “non-learner” status. Learning criteria included at least 40 correct nose pokes, a 3:1 correct:incorrect ratio, and at least 65% of rewards including a social interaction (defined as both experimental and stimulus mice in their respective social interaction zones simultaneously for at least 1 sec of the reward). Testing comprised a 1-hr trial on each of 3 consecutive days, during which the fixed ratio 3 (FR3) reinforcement schedule was used to reward the mouse with a 12-sec social interaction opportunity following three consecutive correct nosepokes. FR3 served to increase social reward seeking effort required to receive a social reward. Following completion of FR3 testing, breakpoint testing was conducted on the following day during a 1-hr trial. To measure the breakpoint, or maximum nosepokes or effort the animal would exhibit for a social reward, the progressive ratio 3 (PR3) reinforcement schedule was used to reward the mouse with a 12-sec social interaction opportunity following a progressive increase in required correct nosepokes by 3 (e.g. 3, 6, 9, 12, etc). Due to the limited number of testing chambers and the length of testing daily, we restricted the number of animals to 17 WTs (10 females, 7 males) and 19 Hets (10 females, 9 males) in order to fit all runs into one day. Task validation data was derived from a cohort of 40 male (n=20) and female (n=20) C57BL/6J adult mice (∼P60), which served as either experimental mice (n=20) that received a social partner interaction as a reward or control mice (n=20) that received only the opening of a door as a reward. The testing procedure was as stated above, except that all mice received four consecutive PR3 testing days to assess reliability of performance within individuals.

##### Sensorimotor battery

To assess sensorimotor capabilities, performance of the mice was measured at P71-P72 in the following series of tasks based on our previously published methods (Maloney et al., 2018b, 2019a). Walking initiation assessed the ability to initiate movement by placing the mouse on a flat surface in the middle of a taped square measuring 21 x 21 cm and recording the time up to 60 sec the animal took to cross the square with all four paws (Fig. 7T). Balance was assessed in the platform test, which requires the animal to balance up to 60 sec on a wooden platform measuring 1.0 cm thick and 3.3 cm in diameter and elevated 27 cm (Fig. 7Q). In the pole test, motor coordination was evaluated as the time the animal took up to 120 sec to turn 180° to face downward and climb down the 57.8 cm pole (Fig. 7R). The 60° and 90° screen tests assessed a combination of coordination and strength as each mouse was required to turn 180° to face upward while in the middle of a 52 cm long wire mesh screen angled 60° or 90° and climb to the top within 60 sec (**Fig. P,SI**). The inverted screen test required muscle strength and endurance for the animal to hang on an inverted wire mesh screen for up to 60 sec (Fig. 7O). The time for each test was manually recorded to the hundredths of a second using a stopwatch. Two trials were conducted for each test and the average of the two was used in analyses. To avoid exhaustion effects, the order of the tests during the first set of trials was reversed for the second set of trials. The order of the tests was not counterbalanced between animals so that every animal experienced each test under the same conditions. All males were tested first, followed by the females. All equipment was cleaned with 70% ethanol between animals.

##### Tube test of social dominance

Mice begin to develop social hierarchy behaviors at 6 weeks of age under laboratory conditions, which result in dominance ranks within their social groups (Hayashi, 1993). The tube test of social dominance was used to assess the social hierarchy behavior of the mice as previously described (Maloney et al., 2018b). Briefly, a pair of sex-matched *MYT1L* Het and WT mice were gently guided into a clear acrylic tube measuring 30 cm in length and 3.6 cm in diameter from either end (Fig. 8A). When the mice met in the center, a divider was lifted and the time for one mouse to back out of the tube as the bout loser/submissive partner up to 2 min was recorded. This was repeated once across three consecutive days for each animal with a novel sex-matched partner. Prior to testing, each mouse was habituated to the tube by gently guiding it through the tube from either end across two consecutive days. The tube was cleaned with 0.02% chlorhexidine diacetate solution (Nolvasan, Zoetis, Parsippany-Troy Hills, NJ) between each pair. Each trial was video recorded and subsequently scored for the dominant and submissive partner of each bout. Because testing required sex-matched genotype-mixed pairs, only a subset of 17 WTs (9 females, 8 males) and 17 Hets (9 females and 8 males) were used.

##### Prepulse inhibition/startle

Sensorimotor gating and reactivity were assessed at P94 in the prepulse inhibition (PPI) /acoustic startle task (Fig. 7U) as previously described (Dougherty et al., 2013). Briefly, PPI (response to a prepulse plus the startle pulse) and acoustic startle to a 120 dBA auditory stimulus pulse (40 ms broadband burst) were measured concurrently using computerized instrumentation (StartleMonitor, Kinder Scientific). A total of 65 trials were presented. Twenty startle trials were presented over a 20 min test period, during which the first 5 min served as an acclimation period when no stimuli above the 65 dB white noise background were presented (non-startle trials). The session began and ended by presenting 5 consecutive startle (120 db pulse alone) trials unaccompanied by other trial types. The middle 10 startle trials were interspersed with PPI trials, consisting of an additional 30 presentations of 120 dB startle stimuli preceded by prepulse stimuli of 4, 12, or 20 dB above background (10 trials for each PPI trial type). A percent PPI score for each trial was calculated using the following equation: %PPI = 100 × (startle pulse alone − prepulse + startle pulse)/startle pulse alone. The apparatus was cleaned with 0.02% chlorhexidine diacetate solution (Nolvasan, Zoetis, Parsippany-Troy Hills, NJ).

##### Fear conditioning

To assess associative memory to an aversive stimuli, we evaluated our mice in the fear conditioning paradigm as we previously described (Maloney et al., 2019a). In this task, freezing behavior was quantified as a proxy for the fear response. Briefly, the apparatus consisted of an acrylic chamber (26 x 18 x 18 cm) with a metal grid floor, an LED cue light and an inaccessible peppermint odorant that is housed in a sound-attenuating chamber (Actimetrics). The cue light turned on at the start of each trial and remained illuminated. The procedure (Fig. S7C) comprised a 5-minute training session, an 8-minute contextual memory test, and a 10 minute cued memory test across 3 consecutive days. During training an 80 dB tone (white noise) sounded for 20 sec at 100 sec, 160 sec and 220 sec. A 1.0 mA shock (unconditioned stimulus; UCS) was paired with the last two sec of the tone (new conditioned stimulus; CS). Baseline freezing behavior was measured during the first two minutes and the freezing behavior as the conditioned response (CR) to the presentation of tone and foot shock was measured during the last three minutes. Freezing behavior was quantified through the computerized image analysis software program FreezeFrame (Actimetrics, Evanston, IL). During contextual conditioning testing on day 2, no tones or shocks were presented allowing for the evaluation of freezing behavior (CR) in response to the contextual cues associated with the shock stimulus (UCS) from day 1. During cued conditioning testing on day 3 the context of the chamber was changed to an opaque acrylic-walled chamber containing a different (coconut) odorant. The 80 dB tone (CS) began at 120 sec and lasted the remainder of the trial. During the first two min baseline freezing behavior to the new context (pre-CS) was measured. During the remaining eight min, freezing behavior (CR) in response to the auditory cue (CS) associated with the shock stimulus (UCS) from day 1 was quantified. Sensitivity to footshocks was evaluated following testing as previously described (Maloney et al., 2019b), and no differences were observed between genotypes (data not shown).

##### Social approach

The three-chamber social approach task was used to test sociability and social novelty preference as previously described (Maloney et al., 2018b). Sociability is defined here as the preference to spend time with a novel conspecific over a novel empty cup. Social novelty is defined as the preference to spend time with a novel versus familiar conspecific. The clear acrylic apparatus measuring 60 x 39 x 22 cm is divided into three equal chambers each measuring 19.5 x 39 x 22 cm with two doors of 5 x 8 cm (Fig. 8B). During testing, an acrylic lid with four air holes is placed on top of the apparatus. Two stainless steel cages (Galaxy Pencil/Utility Cup, Spectrum Diversified Designs, Inc) measuring 10 cm tall and 10 cm in diameter with vertical bars served as conspecific stimulus cages and allowed for controlled, minimal contact interactions between experimental and stimulus mice. The apparatus is placed inside a custom-built sound-attenuating chamber (70.5 × 50.5 × 60 cm). Testing is completed under red light illumination of ∼11 lux provided by LED Flex Ribbon Lights (Commercial Electric, Home Depot). Video is captured by a CCTV camera (SuperCircuits) mounted in the top of each sound-attenuated chamber. A PC computer with the program ANY-maze (Stoelting Co., Wood Dale, IL; http://www.anymaze.co.uk/) recorded video and live tracked the nose, body and tail of the test mouse to produce variables for analysis: distance traveled, time spent in and entries into each chamber and investigation zone. An investigation zone is the area 2 cm outward from the perimeter of each conspecific cage. An entry into the investigation zone requires the nose-point to be within the zone, constituting a purposeful interaction by the test mouse. The social preference score was calculated as (time in social / (time in social + time in empty))*100. The novelty preference score was calculated as (time in novel / (time in novel + time in familiar))*100. Statistical analysis was as previously described (Nygaard et al., 2019).

Testing consists of four, consecutive 10-minute trials. Trials 1 and 2 habituate the test mouse to the center chamber and the whole apparatus, respectively. At the completion of trial 2 the mouse is gently guided back to the center chamber and doors closed. Trials 3 and 4 test sociability and social novelty preference, respectively. In trial 3, an unfamiliar, sex-matched conspecific (C57BL/6J) in a conspecific cage is added to one of outer chambers, and an empty conspecific cages is added to the other outer chamber. The conspecific cage locations were counterbalanced between groups. The test mouse was allowed to explore freely, and at the end of the trial was guided back to the center chamber. During trial 4, a new novel conspecific mouse (C57BL/6J) is added to the empty cage, the conspecific mouse from trial 3 remains in the same cage to serve as the familiar stimulus. After each test, the apparatus is cleaned with 0.02% chlorhexidine diacetate solution (Nolvasan, Zoetis). The conspecific cages were cleaned with 70% ethanol solution.

##### Spontaneous alternation T-maze

The spontaneous alternation T-maze was used to assess perseverative exploratory behavior with procedures adapted from our previous work (Maloney et al., 2018b). The apparatus is made of grey acrylic walls with a clear acrylic floor (Fig. S8D; Noldus). White paper is adhered to the underside of the floor to create distinction between coat color and the apparatus for contrast. A Start chamber (20 x 8.7 cm) is connected to two radiating arms (25 x 8.7 cm), each separated by a door that closes from the floor up. The doors for each arm and start chamber are controlled automatically by Ethovision XT 14 (Noldus) through air compression provided by an ultra-quiet air compressor (California Air Tools) located in an adjacent room. Video is captured by an IR camera (Basler acA1300) mounted above the apparatus, which is connected to a PC computer. Testing is completed in the dark with four IR LED lights (JC Infrared Illuminator) and consists of 10 consecutive trials. Prior to the start of the trial, the mouse is sequestered in the Start chamber for two minutes to habituate to this chamber. To begin the trial, the start door opens, and the mouse is free to explore. An arm choice is made when the whole body crosses the arm threshold located 11.1 cm beyond the door to the arm, and which triggers all doors to close, and the mouse is allowed to explore the chamber for 15 seconds. The door to that arm is then lowered, allowing the animal to move back to the Start chamber, triggering the closing of all doors. After 5s in the Start chamber, the doors all re-open, triggering the beginning of the nex trial. If no arm choice is made after two minutes, it is considered a non-choice trial, and the start of the next trial is triggered. Once all 10 trials are completed the mouse is returned to its home cage and the apparatus cleaned with 0.02% chlorhexidine diacetate solution (Nolvasan, Zoetis).

##### Tactile sensitivity assessment

Tactile sensitivity task assessed reflexive, mechanical sensitivity to a punctate stimulus (von Frey filaments), and was conducted as previously described (Maloney et al., 2018b). The testing apparatus consisted of a metal grid surface elevated 63.5 cm, which allowed access to the plantar surface of each animals’ paws. Each animal was housed in an individual acrylic box (10 cm x 10 cm x 10 cm) open on the bottom and opaque on three sides to prevent visual cues between animals. All mice were acclimated to the testing room 30 min prior to habituation and testing. On days 1 and 2, all mice were habituated to the testing apparatus for 1 hour. On day 3, mice were allowed to acclimate to the testing apparatus for 30 minutes prior to start of testing. Eight different von Frey hair filaments (applying 0.04-2 g of force; North Coast Medical and Rehabilitation Products) were applied to the plantar surface of each animal’s hind paw and withdrawal responses were recorded (Fig. S8E). Presentations started with the lowest filament strength (0.04 g) and increased to the maximum filament strength (2 g). Each filament was applied to the plantar surface of each hind paw five times, and the number of paw withdrawal responses was recorded as percentage of responses. To evaluate the changes in paw withdrawal responses to the whole range of filaments over the testing duration, the area of the curve (AUC) was calculated for each animal.

##### Weight, posture, and physical assessments

All mice from the second cohort were weighed continuously throughout the experiment, starting on P30, to assess obesity-related weight gain in the mice. In addition, on P86, the mice were assessed for posture and physical characteristics. Posture was assessed by picking up the animal by the base of its tail and evaluating the splay of the forelimbs and hindlimbs. Normal posture was defined as splay of both forelimbs and hindlimbs. Abnormal posture was defined as any deviation from this, including hyperflexion or grasping of limbs. Posture was analyzed as a binary measure: normal splayed posture or abnormal posture. The physical examination consisted of assessment of the condition of eyes (presence of debris or cataracts), whiskers (full, partial, pruned), fur (matted or clean), skin (presence of dermatitis), nose (presence of drainage), and anus (presence of prolapse), as well as presence of any seizure-like activity induced by handling or tumors.

### Quantification and statistical analysis

Statistical analyses and graph plottings were performed using IBM SPSS Statistics (v.26), GraphPad Prism (v.8.2.1), and R(v.4.0.0). Prior to analyses, data was screened for missing values and fit of distributions with assumptions underlying univariate analysis. This included the Shapiro-Wilk test on *z*-score-transformed data and qqplot investigations for normality, Levene’s test for homogeneity of variance, and boxplot and *z*-score (±3.29) investigation for identification of influential outliers. Means and standard errors were computed for each measure. Analysis of variance (ANOVA), including repeated measures or mixed models, was used to analyze data where appropriate. Sex was included as a biological variable in all analyses across all experiments. Simple main effects were used to dissect significant interactions. Where appropriate, the Greenhouse-Geisser or Huynh-Feldt adjustment was used to protect against violations of sphericity. Multiple pairwise comparisons were subjected to Bonferroni correction or Dunnett correction. One-sample *t*-tests were used to determine differences from chance. For data that did not fit univariate assumptions, non-parametric tests were used or transformations were applied. For mouse behavior data, the square root transformation was applied to the USV and fear conditioning data. Chi-square or Fisher’s exact tests were used to assess *Myt1l* mutation and sex association with categorical variables. Sex x genotype effects are reported where significant, otherwise data are reported and visualized collapsed for sex. The critical alpha value for all analyses was *p* < .05, unless otherwise stated. Figure schematics were generated using BioRender. The datasets generated and analyzed during the current study are available from the corresponding author upon reasonable request. All statistical data can be found in **Table S5**.

## Acknowledgements

We thank Dr. John Constantino for inspiration and advice, Dr. Monica Sentamet and the Genome Engineering and iPSC Center (GEiC) at Washington University in St. Louis for gRNA design and validation, Dr. Michael White and the transgenic services core for oocyte injection, Dr. Cheng Cheng, Dr. Lingchun Kong, Dr. Xiaoying Chen, Dr. Adam Clemens, Weijia Cao, Brant Swiney, Nicole Fuhler, Rena Silverman, Joelle Schneiderman for technical assistance, and Drs. Carla Yuede and David Wozniak for access to behavioral equipment, and the Mallinckrodt Institute of Radiology’s Small Animal Magnetic Resonance Facility and Washington University Center for Cellular Imaging for technical support. Funding was provided by The Jakob Gene Fund, the Mallinckrodt Institute of Radiology at Washington University School of Medicine, McDonnell International Scholars Academy, and the NIH: (R01MH107515, R01MH124808 to JDD, and NIH 5UL1TR002345 (ICTS) and P50 HD103525 (IDDRC)).

## Supplemental Figures

**Figure S1:**
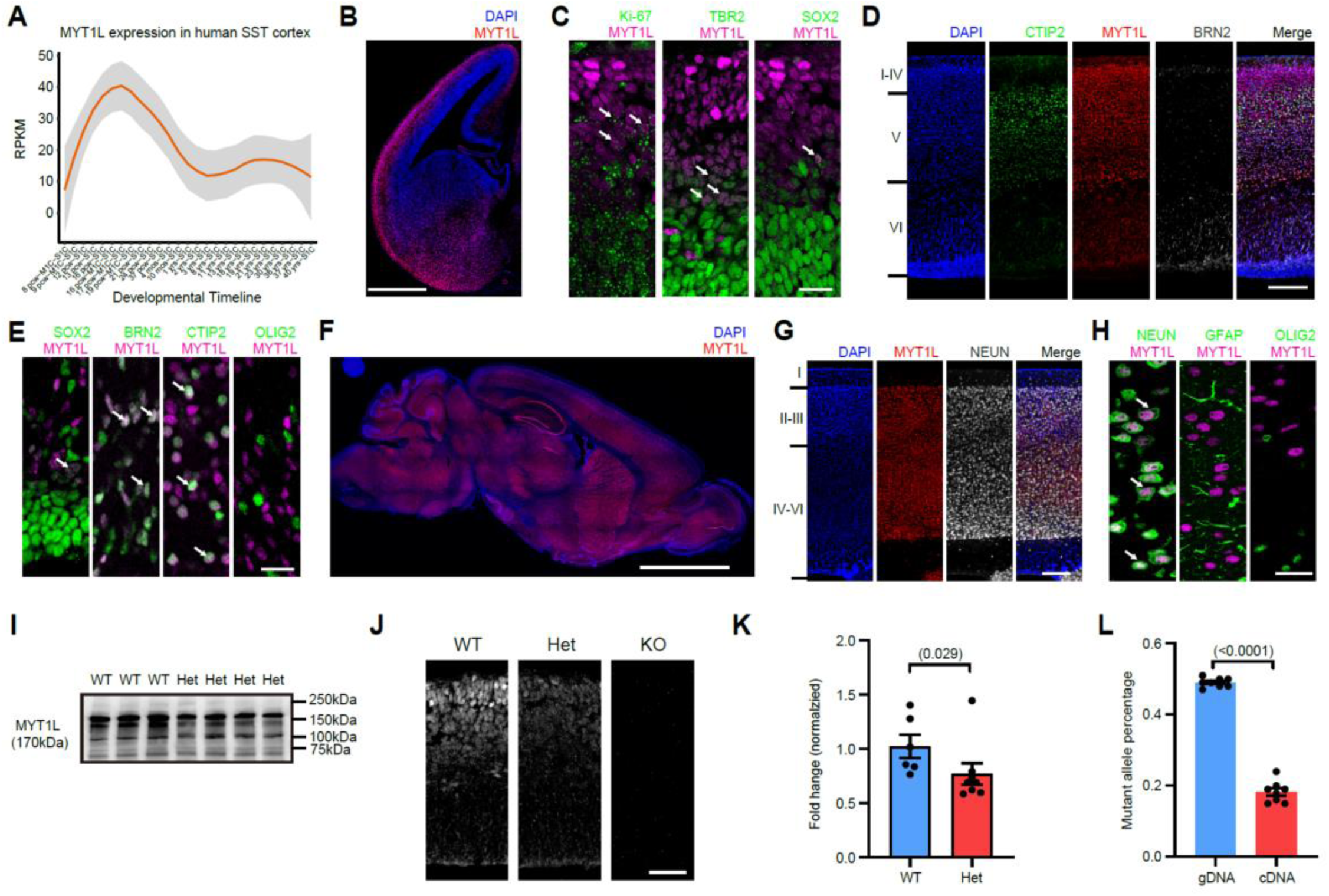
MYT1L protein is expressed in neuronal lineages, peaking during neuronal maturation. Related to Figure 1. **(A)** MYT1L expression across human brain development (somatosensory cortex, data from BrainSpan, https://www.brainspan.org/) also showed peak expression during neuronal maturation, yet sustained expression in adulthood. **(B)** Coronal section of E14 brain immunofluorescence showed MYT1L expression in zones of maturing neurons throughout the brain. **(C)** Immunofluorescence of (Intermediate Zone) in E14 showed transition from cycling (Ki-67+), SOX2, and TBR2 positive progenitors to MYT1L positive cells. Only a small portion of cells showed overlap of these makers. **(D)** MYT1L was expressed in neurons across upper (CTIP2+, green) and lower (BRN+, white) layers of the P1 mouse cortex, **(E)** but not in radial glia (SOX2+) and oligodendrocytes (OLIG2+). **(F)** Sagittal section of adult (P60) mouse brain immunofluorescence showed broad expression of MYT1L in cortex, hippocampus, hypothalamus, striatum, as well as cerebellum. **(G & H)** MYT1L staining in P60 mouse cortex showed its exclusive expression in neurons (NEUN+) but not in astrocytes (GFAP+) and oligodendrocytes (OLIG2+). **(I)** Long exposure of Western blot in Fig. 1G showed no truncated protein produced by *MYT1L* c.3035dupG mutation. **(J)** Immunofluorescence on E14 mouse cortex further validated antibody specificity and protein loss in *MYT1L* KO mice. **(K)** RT-qPCR revealed *MYT1L* relative mRNA expression to GAPDH decreases in P1 Het whole brain lysates (WT n = 6, Het n = 7). **(L)** Illumina sequencing on gDNA and cDNA from P1 *MYT1L* Het mouse brain showed mutant allele-specific loss in cDNA (n = 8), consistent with nonsense mediated decay. *Scale bars, 500 μm in **B**, 20 μm in **C**, 250 μm in **D**, 20 μm in **E**, 3 Mm in **F,** 200 μm in **G**, 20 μm in **H**, and 50 μm **in J**. See* ***Table S5*** *for statistical test details*.

**Figure S2:**
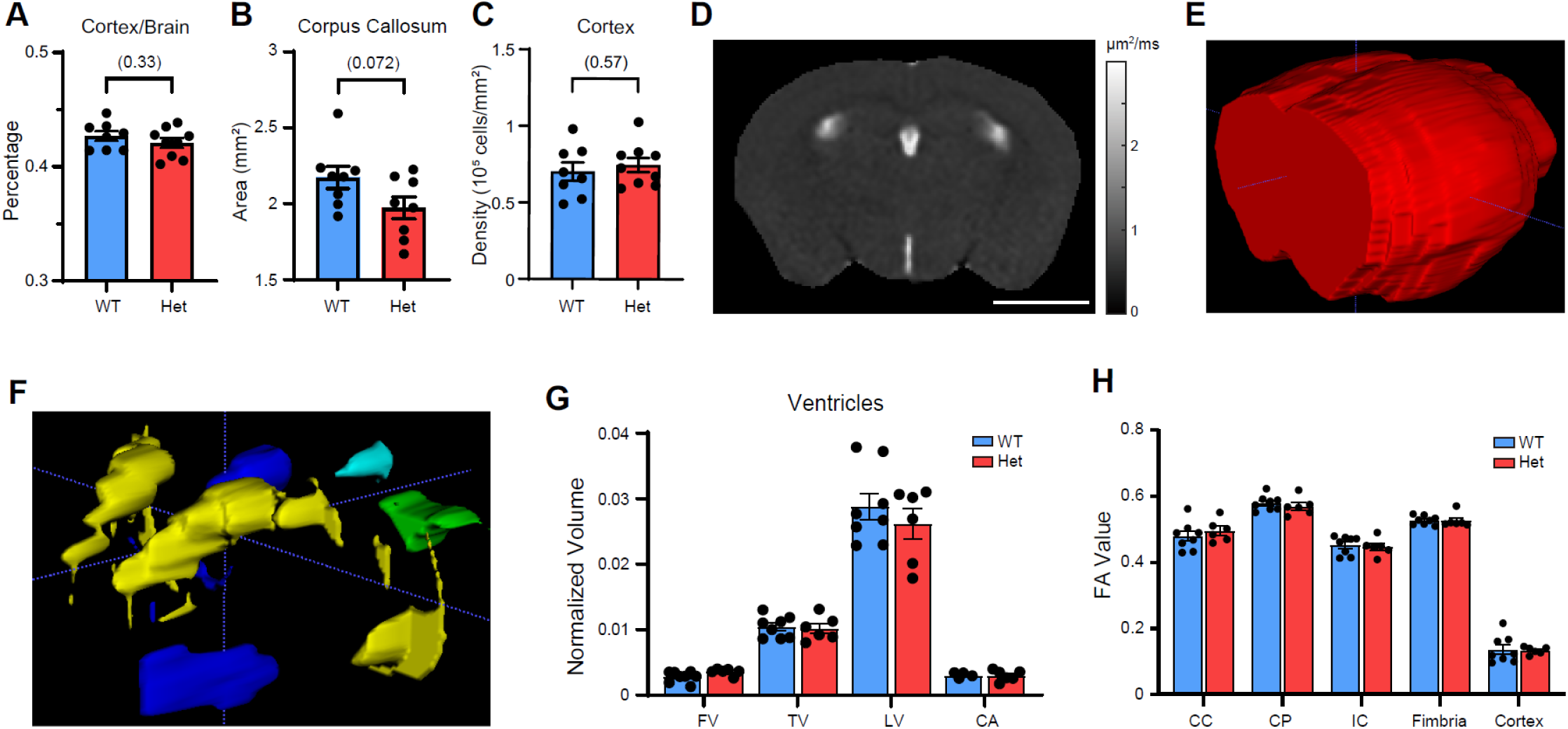
MYT1L haploinsufficiency causes microcephaly and white-matter thinning in corpus callosum. Related to Figure 2. **(A)** The ratio between cortex and brain volume remains unchanged in *Myt1l* Het mice. **(B)** *Myt1l* Het mice had smaller corpus callosum volume (p = 0.072). **(C)** MYT1L loss did not change gross cell density in the brain. **(D)** Apparent diffusion coefficient (ADC) map showing ventricular structures as hyperintense (bright) areas. Scale bar, 0.5 cm. **(E)** 3D reconstruction of the brain contour and **(F)** different ventricles, including the fourth ventricle (FV (green), third ventricle (TV;blue), lateral ventricles (LV;yellow), and cerebral aqueduct (CA;light blue). **(G)** MYT1L loss did not change ventricular sizes. **(H)** FA values were unchanged between Het and WT littermates. *Data are represented as mean ± SEM. See* ***Table S5*** *for statistical test details*.

**Figure S3:**
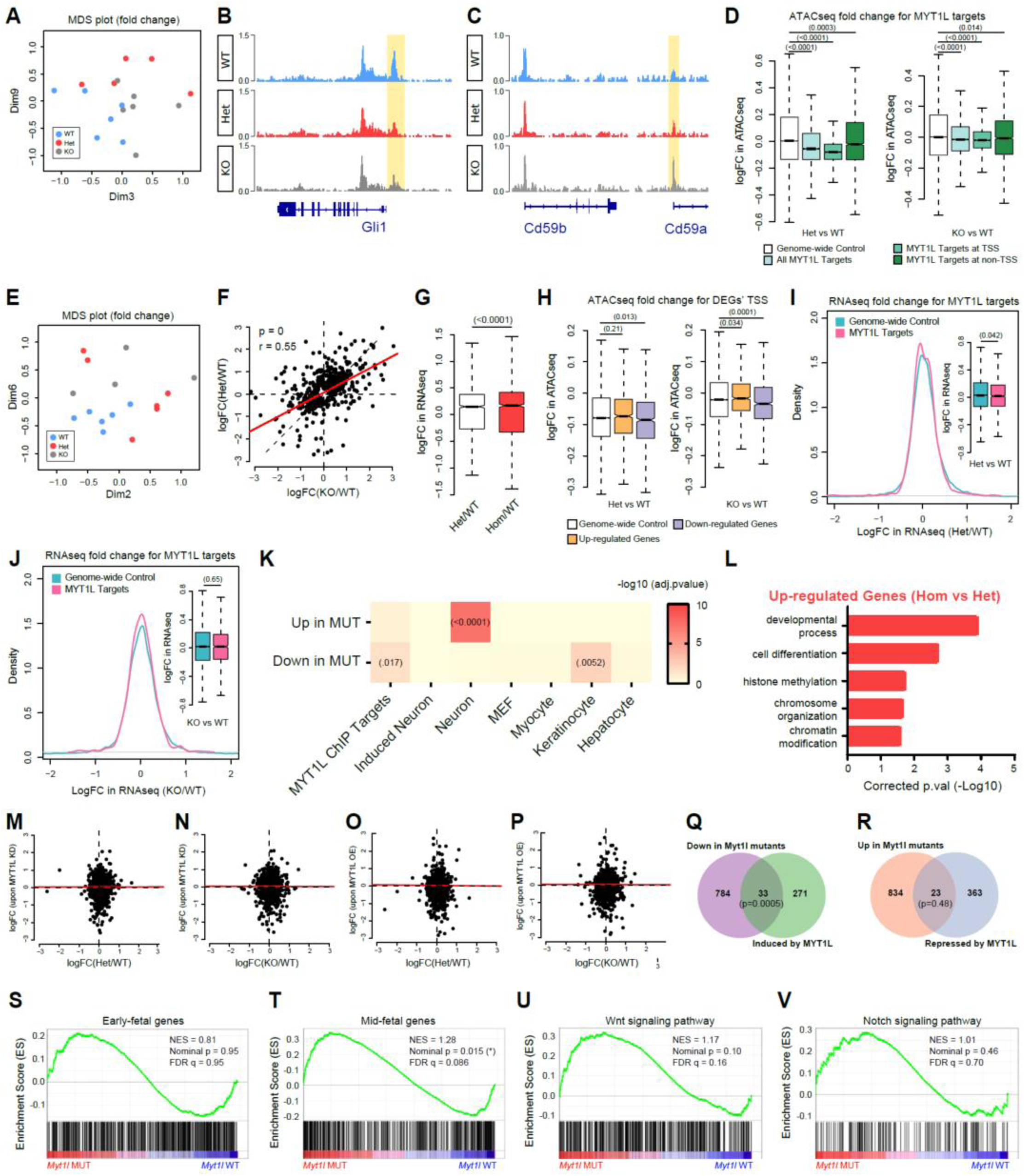
Chromatin Accessibility and RNA-seq analysis define molecular consequences of MYT1L loss in the developing brain. Related to Figure 3. **(A)** MDS plot for ATAC-seq fold changes from individual biological replicates. **(B)** Representative peak of less accessible DAR found in ATAC-seq (highlighted in yellow). **(C)** Representative peak of more accessible DAR found in ATAC-seq (highlighted in yellow). **(D)** ATAC-seq fold changes for MYT1L ChIP targets showed a loss of accessibility at MYT1L bound regions in both Het and KO E14 mice cortex. **(E)** MDS plot for RNA-seq fold changes from individual biological replicates. **(F)** Fold changes of DEGs correlated well between *MYT1L* Het and KO samples. Regression line is shown in red. **(G)** Total loss (KO) of MYT1L had larger effects on fold changes of DEGs compared to partial (Het) MYT1L loss. **(H)** ATAC-seq fold changes for TSS of DEGs identified in RNA-seq. Up-regulated genes in Het E14 mouse cortex had slightly more-accessible TSS while down-regulated genes had significantly less-accessible TSS in ATAC-seq data. The same correlation between DEGs and ATAC-seq was also observed in KO mice with greater significance. **(I)** RNA-seq fold changes for MYT1L targeted genes in (Mall et al., 2017) show subtle downregulation in Het and **(J)** KO E14 CTX expression data sets. **(K)** Overlap of DEGs and different cell-type signature gene lists. Activated genes upon MYT1L loss significantly overlapped neuronal signature genes while genes decreasing expression overlapped with MYT1L embryonic ChIP targets and keratinocyte signature genes. **(L)** GO analysis revealed a further up-regulation of chromatin modification pathways in KO mice compared to Het. **(M)** Correlation of DEGs’ expression changes between Het CTX RNA-seq and primary hippocampal neuron culture MYT1L knockdown (KD) RNA-seq experiments. **(N)** Correlation of DEGs’ expression changes between Het CTX RNA-seq and MEF MYT1L overexpression (OE) RNA-seq experiments. **(O)** Correlation of DEGs’ expression changes between KO CTX RNA-seq and MYT1L KD RNA-seq experiments. **(P)** Correlation of DEGs’ expression changes between KO CTX RNA-seq and MYT1L OE RNA-seq experiments. **(Q)** Venn diagram of overlap between downregulated DEGs in E14 CTX RNA-seq and MYT1L induced genes *in vitro*. **(R)** Venn diagram of overlap between upregulated DEGs in E14 CTX RNA-seq and MYT1L repressed genes *in vitro*. **(S)** GSEA analysis revealed human “early fetal” genes are not categorically affected in mutants while **(T)** “mid-fetal” tended to be up-regulated in mutant E14 mouse cortex. **(U)** GSEA analysis showed no significant categorical shift of Wnt signaling and **(V)** Notch signaling pathway gene expression in *Myt1l* mutant CTX. *Boxplots are plotted with thick horizontal lines as group medians, boxes 25^th^ – 75^th^ percentiles, and whiskers 1.5 x IQR. See* ***Table S5*** *for statistical test details*.

**Figure S4:**
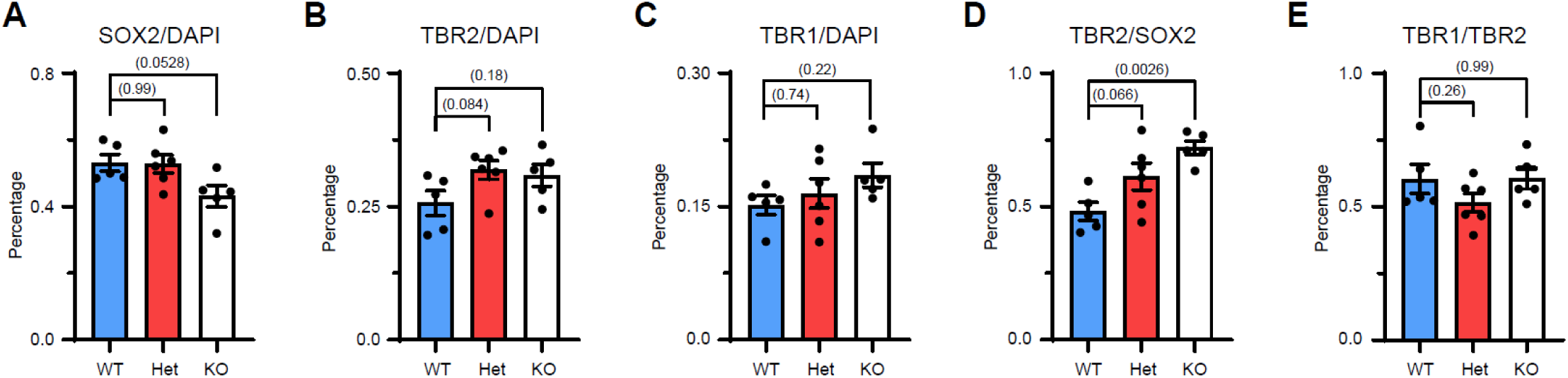
MYT1L loss disrupts progenitor proliferation by precocious cell cycle exit. Related to Figure 4. **(A)** MYT1L KO mice tended to have decreased SOX2(+) cells density compared to WT and Het (*p* = 0.0528) littermates after normalizing to total cell number. **(B)** MYT1L loss did not change normalized TBR2(+) and **(C)** TBR1(+) cell density. **(D)** MYT1L loss altered early progenitor differentiation as shown by TBR2(+)/SOX2(+) ratio. **(E)** MYT1L loss did not alter late progenitor differentiation as shown by TBR1(+)/TBR2(+) ratio. *Data are represented as mean ± SEM and one-way ANOVA was performed with Dunnett correction. See* ***Table S5*** *for statistical test details*.

**Figure S5:**
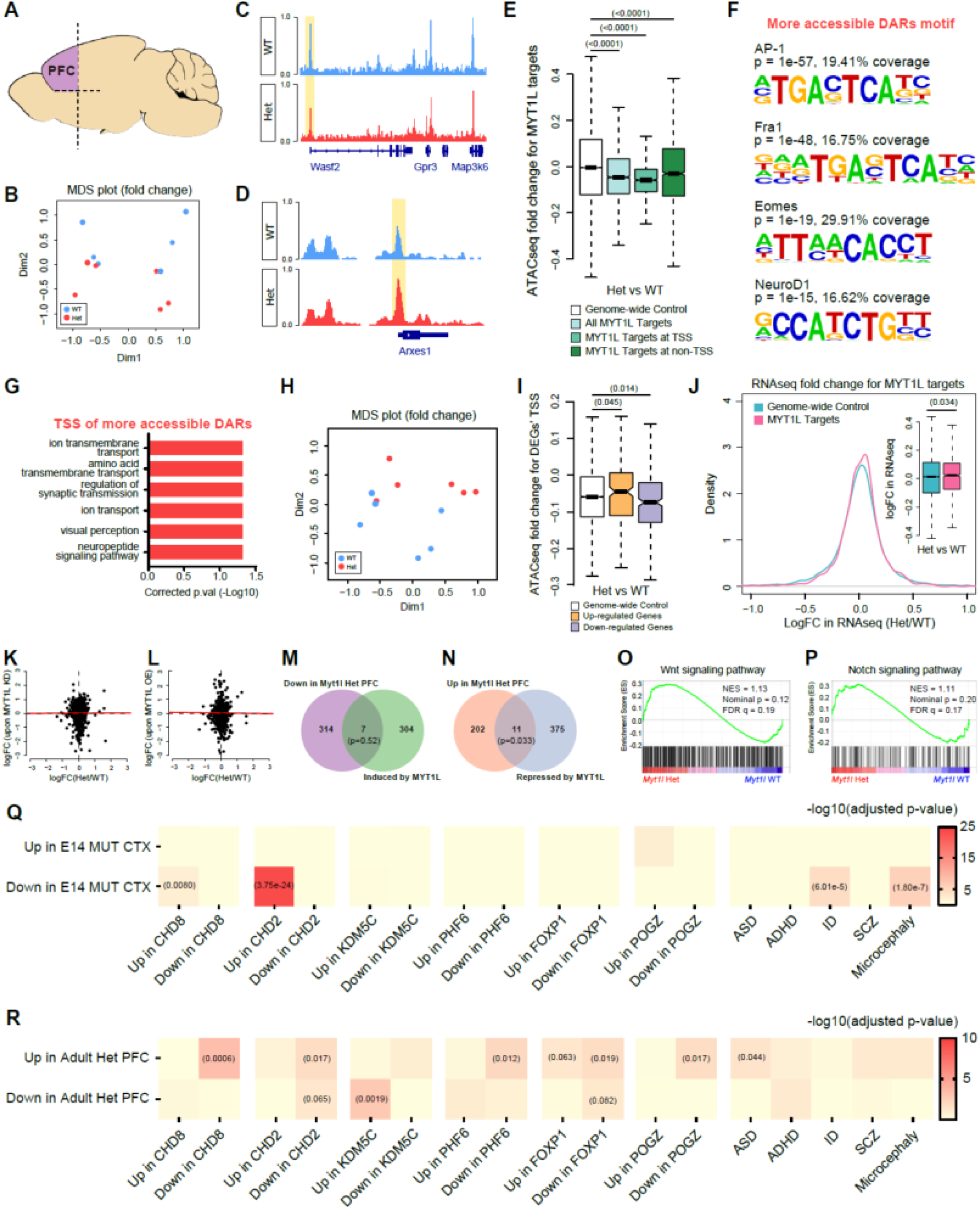
Long term MYT1L deficiency results in arrested maturation of neuronal chromatin and expression patterns. Related to Figure 5. **(A)** Dissection strategy for PFC in adult mouse brain. **(B)** MDS plot for ATAC-seq fold changes from individual biological replicates. **(C)** Representative peak of a DAR with less accessibility in Hets (highlighted in yellow). **(D)** Representative peak of a DAR that was more accessible in Hets (highlighted in yellow). **(E)** ATAC-seq fold changes for MYT1L ChIP targets showed reduced chromatin accessibility at MYT1L bound regions in adult Het mouse PFC. **(F)** Motif analysis comparing more-accessible DARs to less-accessible DARs. **(G)** GO analysis on DAR associated genes showed the dysregulation of neuronal functions in adult Het mouse PFC. **(H)** MDS plot for RNA-seq fold changes from individual biological replicates. **(I)** ATAC-seq fold changes for TSS of DEGs identified in RNA-seq. Upregulated genes in adult Het mouse PFC had more accessible TSS while downregulated genes had less accessible TSS in ATAC-seq data. **(J)** RNA-seq fold changes for MYT1L target genes in (Mall et al., 2017) show subtle upregulation in adult Het mouse PFC. **(K)** Correlation of DEGs’ expression changes between Het PFC RNA-seq and published primary hippocampal neuron culture MYT1L knockdown (KD) RNA-seq experiments. **(L)** Correlation of DEGs’ expression changes between Het PFC RNA-seq and published MEF MYT1L overexpression (OE) RNA-seq experiments. **(M)** Overlap between downregulated DEGs in PFC RNA-seq and MYT1L induced genes *in vitro*. **(N)** Overlap between upregulated DEGs in PFC RNA-seq and MYT1L repressed genes *in vitro*. **(O)** GSEA analysis showed normal expression of Wnt signaling and **(P)** Notch signaling pathway in *Myt1l* Het PFC. **(Q-R)** MYT1L regulated genes were implicated in other ID/ASD mouse models and human genetic data sets. *Boxplots are plotted with thick horizontal lines as group medians, boxes 25^th^ – 75^th^ percentiles, and whiskers 1.5 x IQR. See* ***Table S5*** *for statistical test details*.

**Figure S6:**
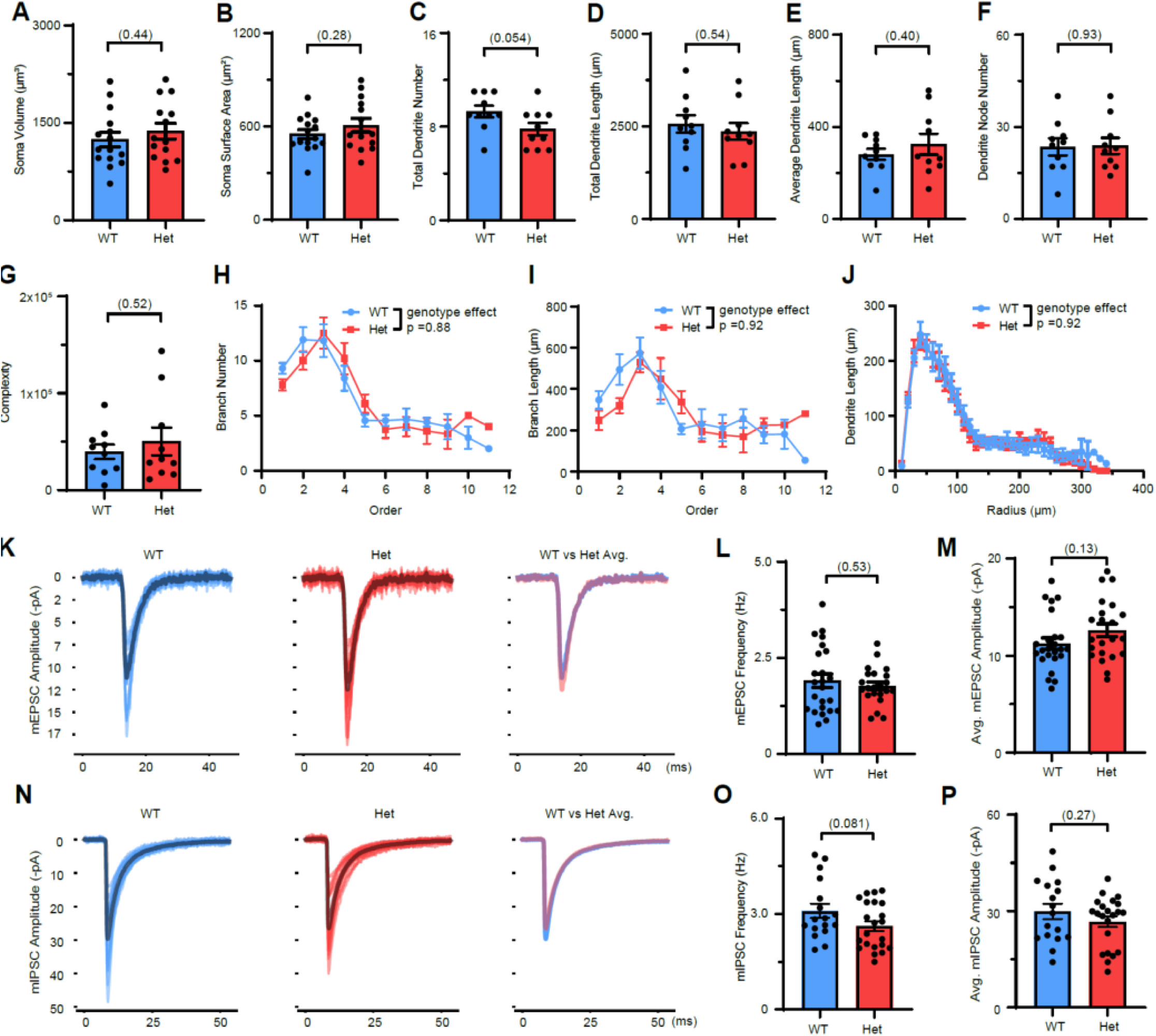
MYT1L haploinsufficiency disrupts baseline neuronal properties and dendritic spine maturity but not neuronal morphology. Related to Figure 6. **(A)** *Myt1l* Het cortical pyramidal neurons had the same soma volume, **(B)** and soma surface area as WT. (C) *Myt1l* Het cortical pyramidal neurons had slightly fewer dendrite numbers compared to WT. (D) *Myt1l* Het cortical pyramidal neurons had the same total dendritic length, **(E)** average dendrite length, **(F)** dendrite node number, **(G)** and dendrite complexity as WT. **(H)** Branch analysis showed no branch number and **(I)** length change in Het neurons. **(J)** Sholl analysis found no dendrite length change across genotypes. **(K)** Overlapped individual mEPSC events of WT (left) neurons, Het (middle) neurons, and averaged mEPSC events (right, blue for WT, red for Het). **(L)** MYT1L loss did not change mEPSC frequency of Het neurons. **(M)** Het neurons have non-significant slightly increased average mEPSC amplitude. **(N)** Overlapped individual mIPSC events of WT (left) neurons, Het (middle) neurons, and averaged mIPSC events (right, blue for WT, red for Het). **(O)** MYT1L loss slightly decreased mIPSC frequency with no influence on **(P)** amplitudes of Het neurons compared to WT. *Data are represented as mean ± SEM. See* ***Table S5*** *for statistical test details*.

**Figure S7:**
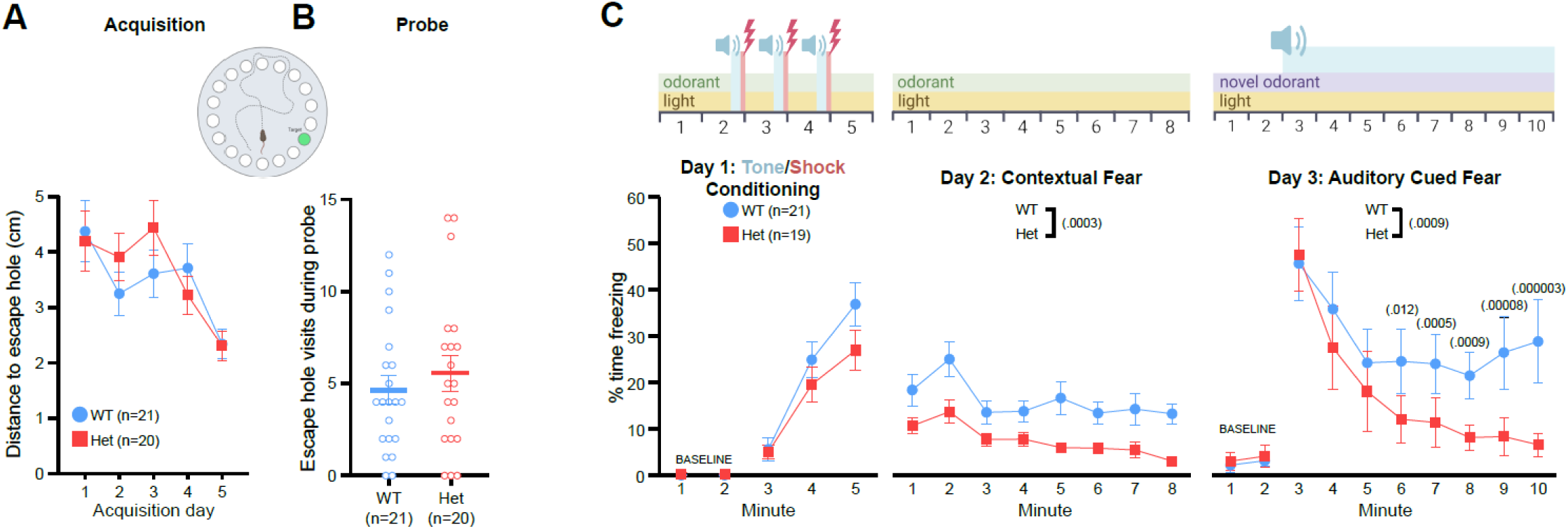
Myt1l haploinsufficiency did not clearly disrupt spatial learning and memory or fear conditioning. Related to Figure 8. **(A)** Distance to reach the escape hole was not different between Hets and WTs during acquisitions trials in the Barnes maze. **(B)** During the Barnes maze probe trial, Hets visited the previously learned escape hole location at a similar frequency to WTs. **(C)** Conditioned fear timeline. Hets froze in response to a pairing of shock and tone/context at a level comparable to WTs on Day 1, yet, froze less during contextual and cued fear tests. No differences were observed for baseline data. *Grouped data are presented as means ± SEM with individual data points as open circles. See also* Figure S8 *and* ***Table S5*** *for statistical test details*.

**Figure S8:**
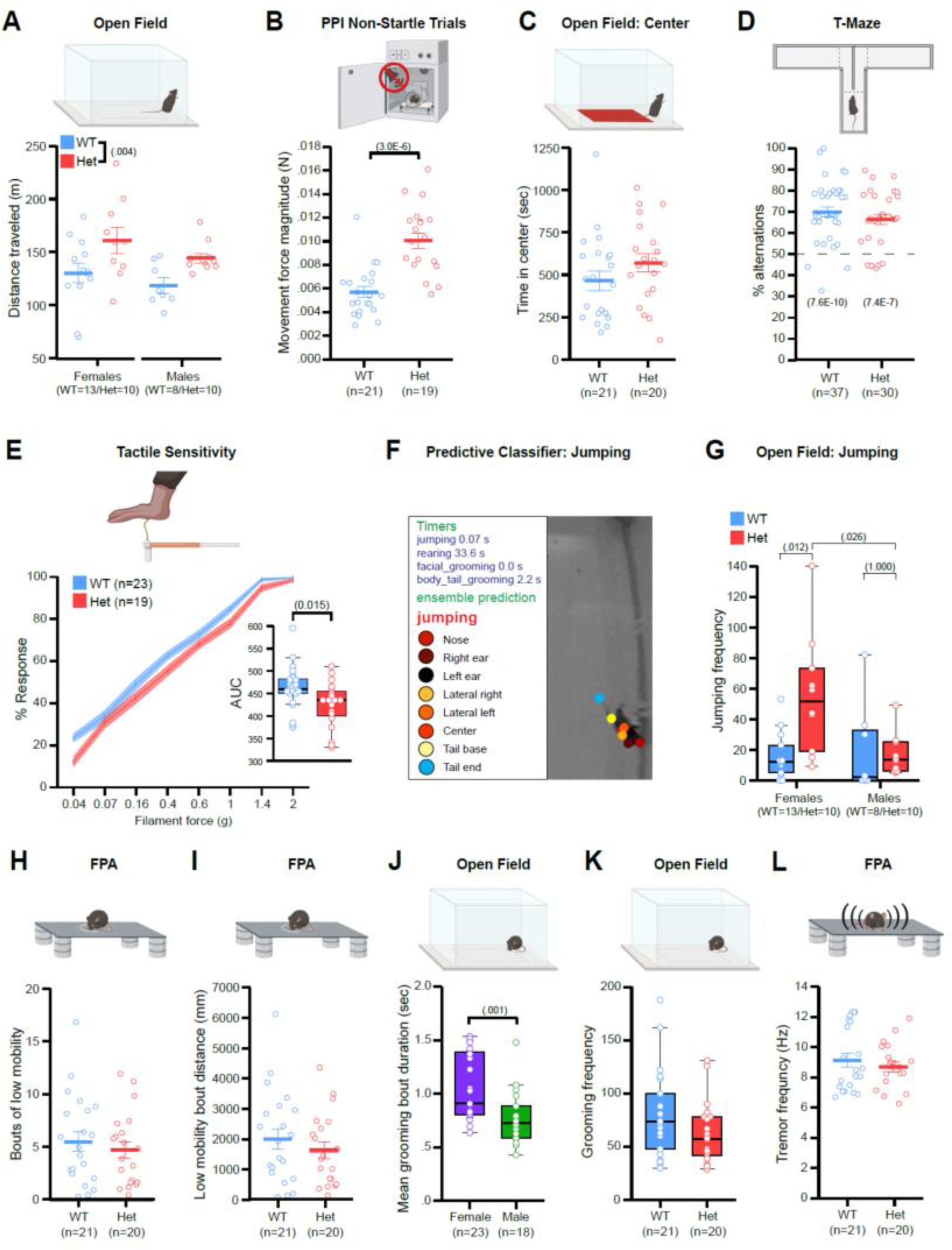
Myt1l haploinsufficiency resulted in hyperactivity and reduced tactile sensitivity without evidence of stereotypies. Related to Figure 8. **(A)** Hets exhibited a longer distance traveled in the open field than WTs. **(B)** During the non-startle trials of the PPI task, Hets exhibited greater movement force magnitude than WTs. **(C)** Hets spent comparable time in the center of the open field chamber to WTs. **(D)** WTs and Hets exhibited % alternations in the T-maze different from chance (50%). **(E)** Het mice responded less than WTs to varying forces of tactile stimulation via von Frey filaments. **(F)** Representative video image frame for SimBA for jumping ensemble prediction. **(G)** Female Hets jumped significantly more than female WTs or male Hets. **(H-I)** In the FPA, Hets demonstrated a comparable number of low mobility bouts and distance traveled during those bouts to WTs. **(J)** Female mice engaged in longer grooming bouts than males. **(K)** Hets did not exhibit altered grooming frequency compared to WTs. **(L)** Hets did not exhibit a difference in tremor frequency in the FPA compared to WTs. *For panels A-F, I, J, and M grouped data are presented as means ± SEM. For panels F (inset), K, and L grouped data are presented as boxplots with thick horizontal lines respective group medians, boxes 25^th^ – 75^th^ percentiles, and whiskers 1.5 x IQR. Individual data points are open circles. See also* Figures 8 and Tab***le S5*** *for statistical test details*.

## Supplemental Information

**Table S1. Characterization of Myt1l Index patient.** The level of functional impairment referable to his autism spectrum disorder (ASD) symptoms was DSM5 Level 1 (“requiring support”). He required frequent verbal (rarely physical) redirection for silliness, perseveration, or engagement in non-preferred tasks. His stereotypic behaviors included repetitive hand wringing, body rocking, stereotypic tensing and vocalizing behaviors. He required a full time paraprofessional in his early school years, but responded significantly to the combination of clonidine and bupropion for improvement in hyperactivity, impulsivity, and aggression, and over time he made incremental gains in composure as well as improved impulse control. A distinct strength was his affable, at times jovial nature; he greatly enjoyed social interactions with people even though his behavior in the context of such interactions could be substantially compromised by immature or perseverative behavior, as well as deficiency in eye gaze. By early adolescence he had mastered enough social interest and competency that he was described as “the mayor” of a summer camp, and he was participating avidly in a musical band. He is able to read, and is very conversational; he has a sense of humor, has made friends, and enjoys telling jokes; his interpersonal exchanges remain silly at times, overly chatty, over-focused on topics of interest to him, and with marginal eye contact and some degree of residual fidgetiness. Medical comorbidities have included obesity and idiopathic scoliosis of adolescence.

**Table S2.** DAR analysis and motif analysis results, Related to Figure 3 and 5

**Table S3.** DGE analysis results and gene lists used for GSEA analysis, Related to Figure 3 and 5

**Table S4.** GO analysis results, Related to Figure 3 and 5

**Table S5.** Statistical analysis results, Related to **Methods**

**Table S6.** Developmental assessment and behavioral testing orders, Related to **Methods**

**Movie S1.** Example DLC pose estimation and SimBA behavior prediction of jumping behavior in Het mice, Related to Figures 7, 8 **and S8.**

